# Nucleoid-binding protein RicO anchors replication origins to the membrane to ensure correct chromosome segregation in *Staphylococcus aureus*

**DOI:** 10.64898/2026.03.06.710092

**Authors:** Simon Schäper, Adrián Izquierdo-Martinez, Qin Liao, António D. Brito, Moritz Sorg, Bruno M. Saraiva, Najwa Taib, Xindan Wang, Mariana G. Pinho

## Abstract

The mechanisms underlying chromosome segregation in bacteria, particularly in non-canonical models, remain incompletely understood. The coccoid bacterium *Staphylococcus aureus* encodes a limited set of conserved proteins involved in chromosome segregation, including the Structural Maintenance of Chromosomes (SMC) complex and an incomplete partition system. Here, we identified a protein of previously unknown function that interacts with the nucleoid and ensures accurate chromosome segregation in *S. aureus*. RicO (Regulator and Insulator of Chromosomal Origins) contains a conserved N-terminal DNA-binding domain that recognizes a specific sequence motif in the origin-proximal region of the chromosome, as well as a C-terminal amphipathic helix that interacts with the cell membrane, thereby bridging DNA and the membrane. RicO localizes in membrane-proximal foci at the tip of each hemisphere (or cell poles). Cells lacking RicO fail to correctly position replication origins at the cell poles and have weakened genomic interactions in the origin-proximal region, resulting in chromosome segregation defects. The combined absence of RicO and a functional SMC complex leads to nearly half of the population appearing as anucleate cells, indicating that these two systems are key contributors to chromosome segregation in *S. aureus.* Collectively, our data support a model in which RicO anchors chromosomal origins at the cell periphery to ensure genome maintenance in this spherical bacterial pathogen.

## Introduction

Faithful chromosome segregation is essential for the survival and proliferation of all life forms. Due to their small size and the absence of a mitotic spindle, bacteria face a particular challenge in maintaining genome integrity while concomitantly condensing and segregating their chromosomes. The mechanisms that orchestrate chromosome segregation and ensure that each daughter cell acquires a complete chromosome copy remain incompletely understood, particularly in non-canonical model bacteria ^1^.

Bacterial chromosome segregation is initiated upon duplication of the replication origin, which subsequently migrates to opposite ends of the cell in many organisms. The Partition (Par) system and Structural Maintenance of Chromosomes (SMC) complex contribute to separate newly replicated origins and organize the two replicated chromosomes into separate cell halves ^2,3^. SMC complexes are present in all eukaryotes and in many bacterial and archaeal species ^4^, while two thirds of all sequenced bacterial genomes contain a *parABS* locus ^5^. ParB binds to origin-proximal *parS* sites generating a large nucleoprotein complex whose migration towards opposite sides of the cell is driven by the motor protein ParA ^6-8^. Upon binding of cytidine triphosphate (CTP), ParB undergoes conformational changes that lead to the formation of a sliding clamp around *parS* sites ^9-11^. Although the Par system is essential in species like *Caulobacter crescentus, Myxococcus xanthus* and *Agrobacterium tumefaciens* ^12-14^, in most cases, inactivation of the Par system causes only moderate defects in the separation of newly replicated origins and the bulk chromosome ^15-19^. SMC complexes are loaded onto the chromosome at *parS* sites in a ParB-dependent manner ^20-23^. Once loaded onto the chromosome, SMC complexes utilize adenosine triphosphate (ATP) to translocate over the DNA and extrude loops, promoting alignment of the two chromosome arms and untangling sister chromosomes ^23-29^. The importance of SMC complexes in chromosome segregation varies across bacterial species, as its loss can lead to temperature-sensitive growth, nucleoid decondensation, and chromosomal partitioning defects in some bacteria ^30-33^, whereas inactivation of the SMC complex in species like *A. tumefaciens*, *Streptococcus pneumoniae, Mycobacterium smegmatis, Corynebacterium glutamicum* and *Deinococcus radiodurans* does not strongly affect chromosome segregation ^14,34-38^. Thus, Par and SMC systems improve the efficiency of chromosome segregation but can be functionally redundant. Furthermore, additional systems and/or passive mechanisms for chromosome segregation may exist in bacteria.

Mechanisms of bacterial chromosome segregation have mainly been studied in rod-shaped bacteria and, to a lesser extent, in cocci. Unlike rod-shaped *Bacillus subtilis,* which encodes a tripartite Par system, the cocci *Staphylococcus aureus* and *S. pneumoniae* encode an incomplete Par system composed only of ParB (also known as Spo0J) and *parS* ^39^. In both species, ParB localizes to the origin of replication by specifically binding to *parS* sites ^34,40,41^. As in *B. subtilis*, the cocci *S. aureus* and *S. pneumoniae* encode an SMC complex composed of a dimer of the eukaryotic condensin homolog SMC, a kleisin homolog ScpA, and a dimer of the accessory protein ScpB ^42,43^. Both ScpA and ScpB are essential for SMC function, forming an SMC-ScpAB ring capable of entrapping DNA ^22^. We have previously shown that deletion of *scpAB* in *S. aureus* results in phenotypic changes similar to those caused by a *smc* mutation ^41^. Notably, individual or combined mutation of *parB* and *smc*/*scpAB* lead to no or only moderate chromosome segregation defects in *S. aureus* and *S. pneumoniae* ^34,41,44,45^, suggesting that additional factors remain to be discovered.

We recently showed that in new-born *S. aureus* cells the partially replicated chromosome adopts a longitudinal (origin-terminus-origin) arrangement, with origins preferably localizing near the tip of each hemisphere, also referred to as ʹcell polesʹ ^41^. However, a mechanism for the tethering of replication origins at this position in *S. aureus* cells has remained unknown thus far. Polar complexes that anchor the origins to the cell pole to lock the chromosome in a longitudinal configuration have been described in various species. In sporulating *B. subtilis* cells, the developmental protein RacA interacts with a centromere-like motif (*ram*) to position the replication origin region near the cell poles ^46-48^. RacA-*ram* nucleoprotein complexes are in turn tethered to the pole by the small peripherical membrane protein DivIVA, which self-associates into an adhesive patch in regions of high membrane negative curvature ^49^. Sporulating cells lacking DivIVA or RacA fail to anchor their origins at the cell poles resulting in pre-spore compartments devoid of a nucleoid ^47,50^. Pole-associated proteins have also been implicated in directly anchoring ParABS in other species, such as PopZ in *Caulobacter crescentus* ^51,52^ and HubP in *Vibrio cholerae* ^53^. This suggests that bacteria have evolved independent systems to maintain their longitudinal organization patterns. A dedicated system for the positioning of replication origins at the periphery of *S. aureus* cells remained to be discovered.

In this study, we identified a previously uncharacterized *S. aureus* protein (RicO, Regulator and Insulator of Chromosomal Origins) and examined its function in chromosome segregation. Cells lacking RicO failed to correctly position replication origins at the cell poles and had defects in chromosome segregation. We show that RicO binds an origin-proximal sequence motif and determines polar localization of replication origins by assembling into membrane-proximal foci at the cell poles. Our findings support a model in which RicO acts as a bridge between replication origins and the cell membrane, thereby ensuring formation of two nucleoid-containing sister cells in *S. aureus*.

## Results

### Identification of RicO as a key protein in *S. aureus* chromosome segregation

To identify new genes required for chromosome segregation in *S. aureus*, we analysed the proportion of cells with nucleoid defects in JE2 mutants of non-essential genes from the Nebraska Transposon Mutant Library (NTML) ^54^, using microscopy images acquired during a previous study ^55^. Out of 1920 mutants, only two showed over 1.75% of cells with nucleoid defects, defined as cells in which fluorescence from the DNA dye Hoechst 33342 was automatically detected below a predetermined threshold in more than a quarter of the cell area. We cannot rule out the possibility that other mutants with nucleoid defects are present in the NTML but escaped our detection due to limitations in the automated quantification. The top mutant contained a transposon insertion in the *scpB* gene, whose deletion in the JE2 strain was previously shown to result in ∼16% anucleate cells ^41,45^, thus validating our screening. The mutant corresponding to the second hit contained a transposon insertion in the previously uncharacterized *SAUSA300_0383* gene. The encoded protein contains an N-terminal putative DNA-binding domain (DBD) and is hereafter referred to as RicO (Regulator and Insulator of Chromosomal Origins) based on our findings described below.

To confirm that the observed phenotype was associated with the *ricO* gene, we first deleted *ricO* in the methicillin-resistant *S. aureus* (MRSA) strains JE2 and COL, as well as in the methicillin-sensitive *S. aureus* (MSSA) strain NCTC8325-4, and manually quantified the fraction of anucleate cells, which resulted in similar numbers (∼4%, ∼5% and ∼3%, respectively) (Fig. 1a; Supplementary Figure 1a). Normal chromosome segregation in the JE2Δ*ricO* strain could be restored with plasmid-encoded *ricO* (Fig. 1a). Despite chromosome segregation defects, growth rate of JE2 was maintained in the absence of RicO (Supplementary Figure 2a).

**Figure 1.**
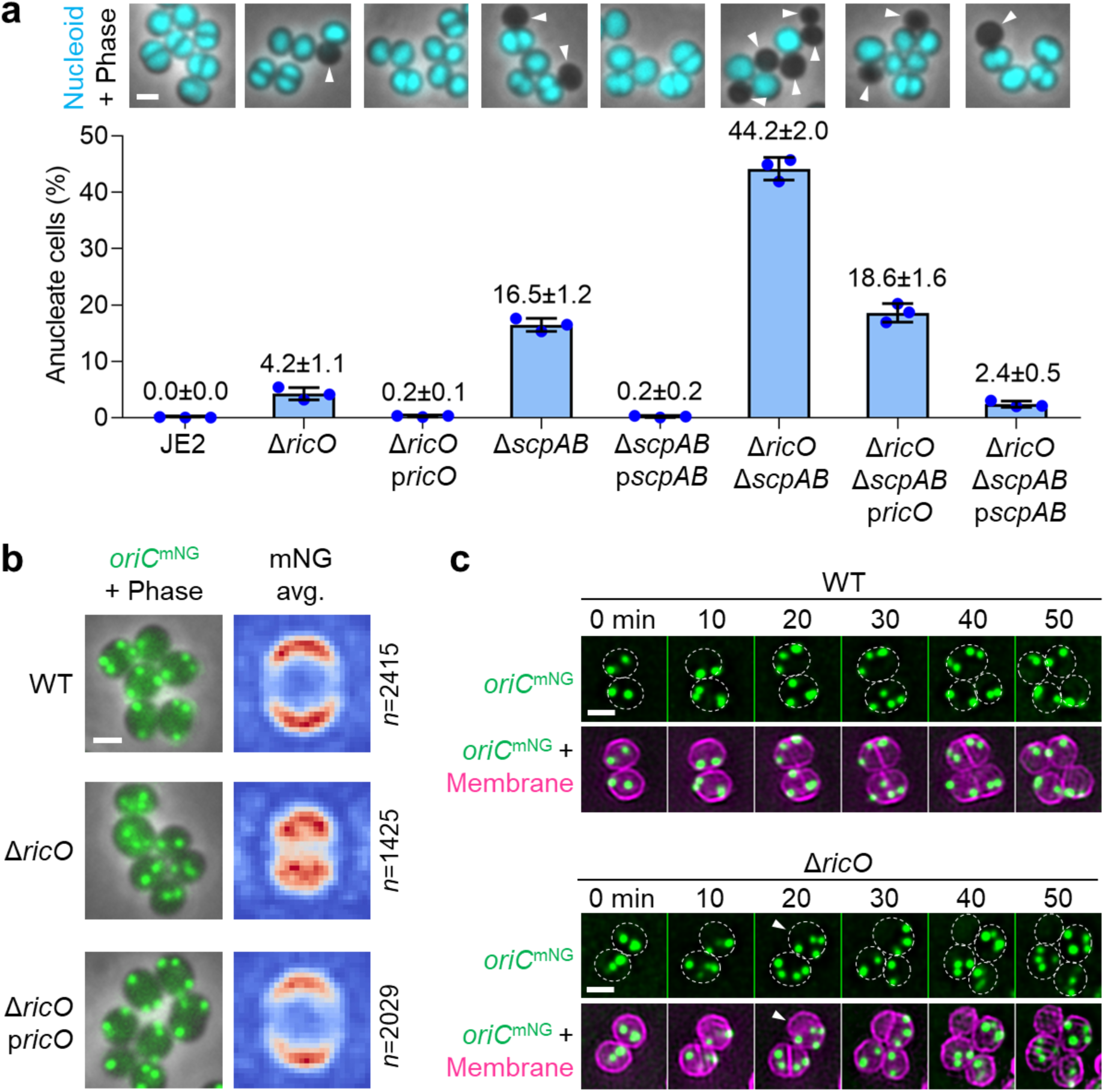
Cells lacking RicO show defects in chromosome segregation and loss of polar localization of chromosomal origins. a) Deletion of *ricO* results in chromosome segregation defects and in synthetic sickness of cells lacking a complete SMC complex. The bar graph shows the percentage of anucleate cells measured for the strains (from left to right) JE2, JE2Δ*ricO,* JE2Δ*ricO*p*ricO,* JE2_Δ*scpAB,* JE2_Δ*scpAB+*ScpAB, JE2Δ*ricO*Δ*scpAB*, JE2Δ*ricO*Δ*scpAB*p*ricO* and JE2Δ*ricO*Δ*scpAB*p*scpAB*. Bars represent the mean and lines indicate the standard deviation of three independent experiments. At least 1170 cells were analysed for each biological replicate. Fluorescence micrographs overlaid with phase contrast images show cells labelled with DNA dye Hoechst 33342 (cyan), and anucleate cells are indicated by white arrowheads. Scale bar, 1 µm. b) RicO is required for membrane-proximal localization of chromosomal origins. Fluorescence micrographs overlaid with phase contrast images of the strains (from top to bottom) JE2_FROS^Ori^ (WT), JE2*oriC*^mNG^Δ*ricO* and JE2*oriC*^mNG^Δ*ricO*p*ricO.* Replication origins (green) were fluorescently labelled by producing TetR-mNG which binds to a *tetO* array introduced at position 358° of the circular genome. Images on the right show heat maps of the average localization of detected fluorescent foci corresponding to replication origins of the same strains as cells shown on the left. Blue-white-red color code in heat maps represents spot density from low to high, respectively. *n*, number of analysed cells. Scale bar, 1 µm. c) Cells lacking RicO fail to equally distribute replicated origins between cell halves. Cells of the strains JE2_FROS^Ori^ (top) and JE2*oriC*^mNG^Δ*ricO* (bottom), each with replication origins fluorescently labelled (green) and membrane stained with CellBrite Fix 640 (magenta), were imaged by time-lapse microscopy. Overlaid fluorescence micrographs show asymmetric distribution of origins between two cell halves of the JE2*oriC*^mNG^Δ*ricO* strain (indicated by white arrowheads). Cell outlines inferred from stained membranes are indicated by dashed white ovals. Scale bar, 1 µm.

Anucleate cells can arise through various mechanisms, such as impaired DNA replication, DNA degradation or incorrect chromosome segregation. To understand the origin of Δ*ricO* anucleate cells, we initially questioned whether cells lacking RicO could have a block in chromosome replication that would result in partially replicated chromosomes that fail to be efficiently segregated. To test this hypothesis, we performed genome-wide marker-frequency analysis (MFA) of exponentially growing cells using next-generation sequencing. MFA has been used in various bacteria, including *S. aureus* ^41,56^, to detect changes in replication by quantifying the relative abundance of DNA sequences across the entire genome. The overall replication profile was similar for the JE2Δ*ricO* deletion and JE2 wild-type strains (Supplementary Figure 3), suggesting that RicO is not involved in chromosome replication.

We then studied the localization and dynamics of chromosomes during the cell cycle of the JE2Δ*ricO* strain by time-lapse fluorescence microscopy. We did not see evidence of septum assembly over the nucleoid, which could cause DNA damage. However, while JE2 wild-type cells visibly directed one copy of the chromosome to each cell half concomitantly with the onset of septum synthesis at mid-cell, leading to the formation of two nucleoid-containing sister cells, a fraction of Δ*ricO* cells failed to equally distribute newly replicated chromosomes between both cell halves before septum completion. This gave rise to asymmetric sister cells, only one of which contained DNA (Supplementary Figure 1b), and led us to test for complementary roles in chromosome segregation of RicO and other proteins with a known role in that process. Specifically, we deleted *scpAB* (components of the SMC complex), *parB* and *ricO* in various combinations in the background of the JE2 strain. Strikingly, the JE2Δ*ricO*Δ*scpAB* double deletion strain showed anucleate cells with a proportion of ∼45%, which was significantly higher than the sum of anucleate cell fractions in JE2Δ*ricO* (∼4%) and JE2_Δ*scpAB* (∼16%) single deletion strains (Fig. 1a). The synthetic sickness resulting from *ricO* and *scpAB* deletions was also reflected in the reduced growth of Δ*ricO*Δ*scpAB* cells (Supplementary Figure 2a). Complementation of the JE2Δ*ricO*Δ*scpAB* strain with either plasmid-encoded *ricO* or *scpAB* restored the fraction of anucleate cells to levels of the corresponding single deletion mutants (Fig. 1a). Deletion of *parB* alone or in combination with *ricO* and *scpAB* deletions did not impair chromosome segregation (Supplementary Figure 4a). Surprisingly, the anucleate cell proportion of the JE2Δ*ricO*Δ*parB* double deletion strain (1.6±0.1%) was approximately two-fold reduced relative to the JE2Δ*ricO* single deletion strain (3.8±0.5%) (Supplementary Figure 4a), indicating that *parB* deletion alleviated defects caused by *ricO* deletion. Taken together, these data suggest that RicO has a function in chromosome segregation.

### RicO determines polar localization of chromosomal origins

To understand the role of RicO in chromosome segregation, we examined the localization of the chromosomal origin of replication (*oriC*) during the cell cycle of Δ*ricO* cells. We probed *oriC* in live *S. aureus* cells using a fluorescence reporter-operator system (FROS), as previously described ^41^. Briefly, an array of 48 *tetO* operator sequences was inserted into the circular JE2 chromosome (at 358° or ∼14 kb from *oriC*) and a gene encoding TetR translationally fused to mNeonGreen (mNG) was expressed from the chromosomal *spa* locus under the control of a cadmium-inducible promoter. Insertion of the (*tetO*)^48^ array near *oriC* and leaky expression of *tetR-mNG* resulted in the formation of bright fluorescent foci without notably affecting cell viability, chromosome segregation or DNA replication ^41^. Visualization of fluorescently labelled origins in live cells of the JE2_FROS^Ori^ strain revealed the previously described *oriC* localization pattern at the cell poles (Fig. 1b) ^41^. Strikingly, the JE2*oriC*^mNG^Δ*ricO* deletion strain showed loss of polar *oriC* localization, as fluorescent foci localized on average over the nucleoid and further away from the membrane relative to wild-type cells (Fig. 1b).

Polar *oriC* localization was recovered in Δ*ricO* cells upon plasmid-encoded expression of *ricO* (Fig. 1b). Noteworthy, membrane-distal *oriC* localization is characteristic for Δ*ricO* cells because no such pattern was observed in previously studied mutants of other genes involved in chromosome segregation, including *smc* and *parB* ^41^. Consistent with unchanged replication profile (Supplementary Figure 3), *ricO* deletion did not change the overall average number of fluorescent *oriC* foci per cell (Supplementary Figure 5). These data indicate that RicO plays a role in anchoring replication origins to the cell poles, without affecting replication.

To further characterize aberrant *oriC* localization in Δ*ricO* cells, we studied the spatiotemporal dynamics of *oriC* in actively growing cells by time-lapse fluorescence microscopy. In cells of the JE2_FROS^Ori^ strain, *oriC* localized as two foci near opposite cell poles and the duplication of each focus was followed by their rapid segregation within cell halves separated by the developing septum (Fig. 1c). Nascent cells of the JE2*oriC*^mNG^Δ*ricO* strain also exhibited two *oriC* foci, which duplicated and initially segregated, but failed to be directed to both cell halves in some cells, giving rise to two asymmetric sister cells, one containing four *oriC* foci and the other none (white arrowheads in Fig. 1c). These data suggest that in the absence of RicO, replicated origins are not always equally distributed between daughter cells, reflecting RicO’s important role in chromosome segregation.

### RicO localizes at the cell poles similarly to chromosomal origins

Next, we studied the subcellular localization of RicO relative to four different chromosomal regions, namely *oriC*, left arm, right arm, and terminus of replication (*terC*). For this, we started by constructing translational fusions of fluorescent proteins to the N- and C-terminal ends of the RicO protein. However, this approach yielded non-functional protein fusions. We then generated a ‘sandwich’ fusion of RicO to mCherry (RicO^SW^-mCherry) by inserting the mCherry sequence between the coiled-coil and disordered regions (between valine 182 and threonine 183; yellow arrowhead in Supplementary Figure 6a). We confirmed that this fusion was functional, as indicated by the absence of anucleate cells in the JE2*ricO^SW^-mcherry* strain, and determined its integrity by in-gel fluorescence detection (Supplementary Figure 7a,b). The RicO^SW^-mCherry fusion formed membrane-proximal, bright fluorescent foci/patches that localized at the cell poles (Fig. 2a). RicO^SW^-mCherry preferentially colocalized with *oriC* rather than with the left and right chromosome arms, or with *terC* (Fig. 2b). As a positive control for colocalization, we used dual labelling of proteins from the SMC complex by generating fusions of ScpA and ScpB to mNG and mCherry, respectively, which co-localized in foci over the nucleoid (Fig. 2a,b). These data suggest that RicO and replication origins occupy the same region near the cell poles for a considerable period during the cell cycle.

**Figure 2.**
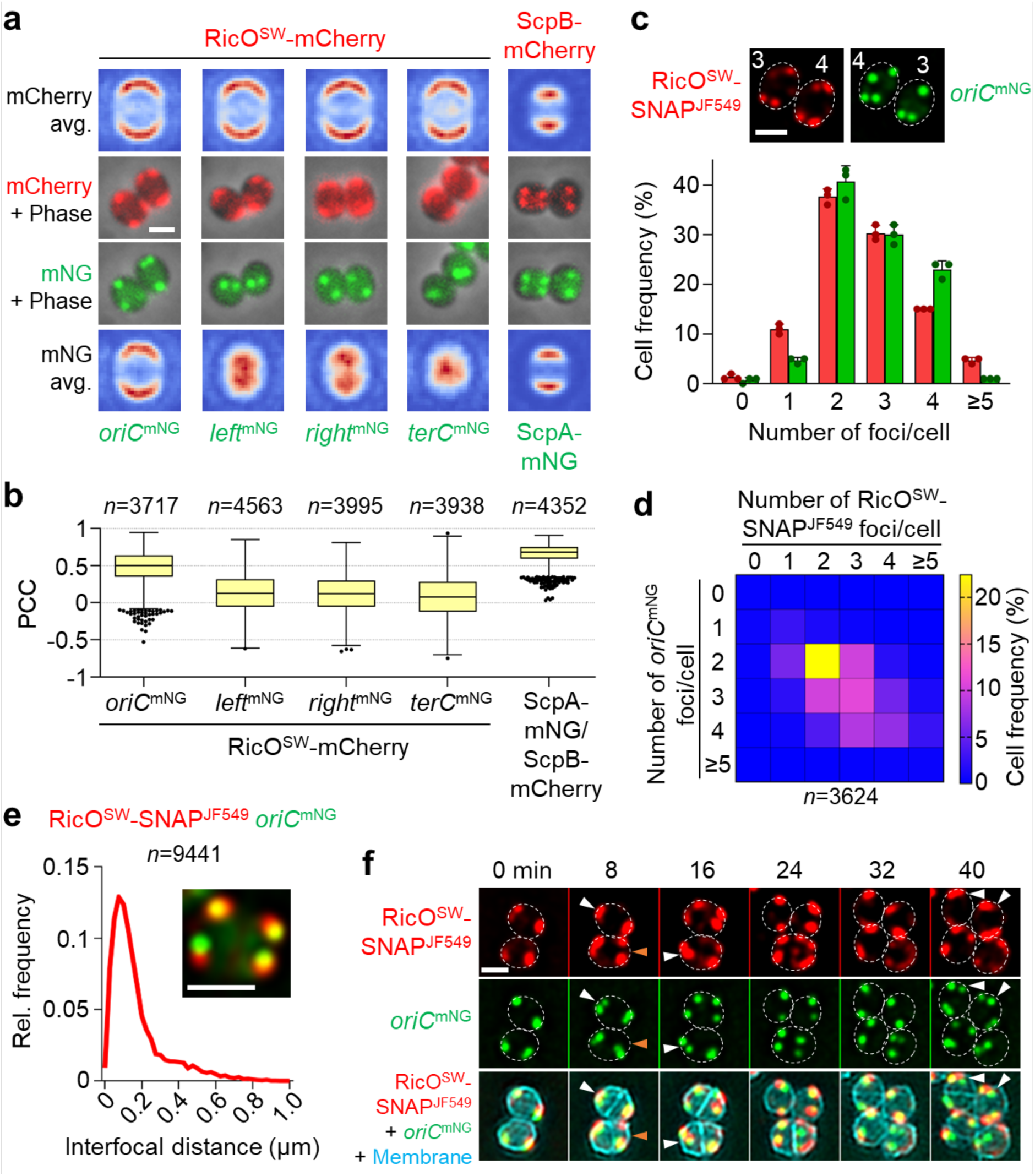
RicO forms membrane-proximal foci in the vicinity of chromosomal origins. a) Polar localization of RicO resembles that of replication origins. Fluorescence micrographs overlaid with phase contrast images of the strains (from left to right) JE2*oriC*^mNG^*ricO^SW^-mcherry*, JE2*left*^mNG^*ricO^SW^-mcherry*, JE2*right*^mNG^*ricO^SW^-mcherry*, JE2*terC*^mNG^*ricO^SW^-mcherry* and JE2*scpA-mNGscpB-mcherry.* Each strain produced mCherry either fused to RicO or to ScpB (both in red), as well as mNeonGreen either fused to TetR or to ScpA (both in green). Strains producing TetR-mNG contained a *tetO* array at positions 358°, 278°, 92° or 182° of the circular genome to fluorescently label the replication origin (*oriC*), left and right chromosome arms (*left*, *right*), or the replication terminus (*terC*), respectively. Top and bottom rows show heat maps of the average localization of detected fluorescent foci corresponding to RicO and ScpB (top), or to different chromosomal loci and ScpA (bottom) of the same strains as cells shown in the middle rows. Blue-white-red color code in heat maps represents spot density from low to high, respectively. Data shown are from three biological replicates and from a minimum of 3564 cells analysed for each strain. Scale bar, 1 µm. b) RicO localizes near replication origins. Pearson Correlation Coefficient (PCC) was used as a measure of colocalization between pairs of fluorescent proteins produced in the same strain. A PCC value close to one denotes colocalization, whereas a negative value denotes anti-colocalization. ScpB-mCherry and ScpA-mNG fusions were used as positive control, as they are part of the SMC complex. Data are represented as box-and-whisker plots in which boxes correspond to the first-to-third quartiles, lines inside the boxes indicate the median, and the ends of whiskers and outliers follow a Tukey representation. PCC values were obtained from the same images as data in panel a. *n*, number of analysed cells. c) RicO and replication origins show a similar distribution for the number of foci per cell. The bar graph shows the frequency of cells classified by the number of fluorescent foci corresponding to RicO^SW^-SNAP (red) and replication origins (green). Fluorescence micrographs show cells of the strain JE2*oriC*^mNG^*ricO^SW^-snap* with RicO labelled with SNAP-tag ligand JF549 (red) and producing TetR-mNG to fluorescently label replication origins (green). Numbers indicate the number of fluorescent foci per cell determined in each fluorescence channel. Bars represent the mean and lines indicate the standard deviation of three independent experiments. At least 1136 cells were analysed for each biological replicate. Cell outlines inferred from stained membranes are indicated by dashed white ovals. Scale bar, 1 µm. d) Number of RicO foci and replication origins in a cell can differ. The heat map shows the frequency of cells displaying the indicated numbers of fluorescent foci corresponding to RicO^SW^-SNAP and replication origins that were detected in the same cell. Percentages were obtained from the same images as data in panel c. *n*, number of analysed cells. e) RicO foci localize in the vicinity of replication origins. Histogram of the distance between a given fluorescent focus of RicO^SW^-SNAP and its nearest replication origin that was detected in the same cell. Bin width, 0.025. Centre of first/last bin, 0/1. Overlaid fluorescence micrograph shows a single cell of the strain JE2*oriC*^mNG^*ricO^SW^-snap* with RicO labelled with SNAP-tag ligand JF549 (red) and with fluorescently labelled replication origins (green). Distances were obtained from the same images as data in panel c. *n*, number of analysed RicO^SW^-SNAP/*oriC*^mNG^ pairs. Scale bar, 1 µm. f) Splitting of RicO foci and separation of newly replicated origins in dividing *S. aureus* cells. Cells of the strain JE2*oriC*^mNG^*ricO^SW^-snap*, producing fluorescently labelled replication origins (green), were labelled with SNAP-tag ligand JF549 (red) and the membrane dye CellBrite Fix 640 (cyan), and imaged by time-lapse microscopy. Initial segregation of replication origins prior and after splitting of RicO^SW^-SNAP foci within cell halves is indicated by white and orange arrowheads, respectively. Cell outlines inferred from stained membranes are indicated by dashed white ovals. Scale bar, 1 µm.

To further study RicO localization relative to replication origins, we used a functional RicO ‘sandwich’ fusion to SNAP-tag (RicO^SW^-SNAP). SNAP-tag is a self-labelling protein tag to specifically and covalently label fusion proteins with small synthetic molecules ^57^. RicO^SW^-SNAP labelled with red-fluorescent, cell-permeable SNAP-tag ligand JF549-cpSTL was brighter than RicO^SW^-mCherry and subject to no apparent proteolytic cleavage (Supplementary Figure 7b). Firstly, we determined the number of fluorescent foci corresponding to RicO^SW^-SNAP and *oriC* in each cell of the strain JE2*oriC*^mNG^*ricO^SW^-snap*. Our previous data indicate that most cells in an exponentially growing culture display two to four *oriC* foci ^41^. Approximately 44% of cells displayed the same number of RicO^SW^-SNAP and *oriC* foci (Fig. 2c,d) and the overall number of foci per cell was similar (2.60±0.04 and 2.73±0.04, respectively). Secondly, we measured the distance between the centre of a given RicO^SW^-SNAP focus to the nearest *oriC* focus detected in the same cell. The distribution of this distance showed a sharp peak at ∼0.1 µm, equivalent to about 1/10 of the cell diameter (Fig. 2e). These data further suggest that RicO localizes in the vicinity of replication origins but does not always colocalize with the origins.

Given that RicO is usually found near, but not always overlapping with chromosomal origins, we hypothesized that movement of RicO foci could precede movement of the origins along the path of chromosome segregation, perhaps promoting origin segregation. To better understand the dynamics underlying RicO localization, we next imaged RicO^SW^-SNAP in actively dividing cells by time-lapse microscopy (Fig. 2f). Most JE2*oriC*^mNG^*ricO^SW^-snap* cells in an early stage of the cell cycle displayed two bright RicO^SW^-SNAP foci at opposite cell poles. During the cell cycle, each focus underwent splitting while maintaining polar localization. However, splitting of RicO^SW^-SNAP foci was observed to occur both prior (orange arrowheads in Fig. 2f), as well as after (white arrowheads in Fig. 2f), initial segregation of newly replicated origins, with the two events following no established order. The asynchronous splitting of foci was also reflected in the proportion of cells (∼40%) displaying an unequal number of RicO^SW^-SNAP and *oriC* foci ranging from two to four (Fig. 2d), indicating that RicO foci could split before *oriC* foci and *vice versa*. Taken together, these data suggest that RicO assembles into membrane-proximal foci at the cell poles where it transiently colocalizes with replication origins but is unlikely to actively drive origin segregation.

### RicO binds a specific sequence motif

Based on the observations above, we hypothesized that RicO may interact directly with the chromosome. To test this hypothesis, we performed chromatin immunoprecipitation sequencing (ChIP-seq) using an antibody raised against purified RicO, whose specificity was confirmed by Western Blot analysis (Supplementary Figure 7c,d). ChIP-seq of the strain JE2Δ*spa* identified nine enrichment peaks within an approximately 700 kb genomic region flanking *oriC,* all of which were lost upon deletion of *ricO* in the JE2Δ*spa* strain (Fig. 3a). Motif discovery algorithms from the MEME Suite ^58^ identified a putative binding site (named *ricS*) that was present within each enrichment peak (Fig. 3b; Supplementary Figure 8a). The JE2Δ*spa* strain lacking the nine *ricS* sites lost the RicO enrichment peaks (Fig. 3a), while cellular levels of RicO were not markedly changed by *ricS* deletions (Supplementary Figure 7d).

**Figure 3.**
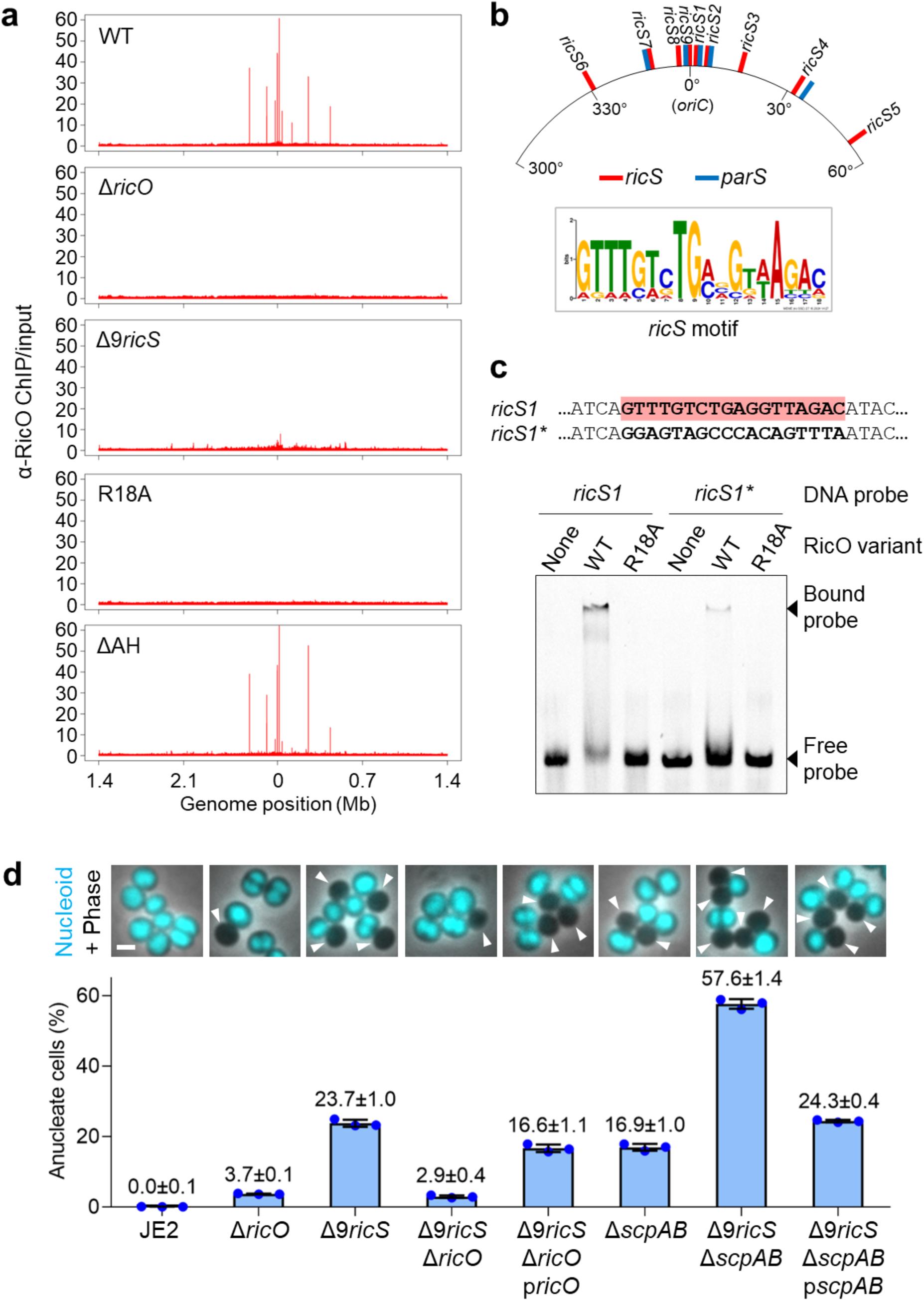
RicO binds to nine specific sites in the origin-proximal region of the chromosome. a) ChIP-seq reveals nine genomic loci targeted by RicO. Enrichment profiles were obtained from anti-RicO ChIP-seq performed in the strains (from top to bottom) JE2Δ*spa* (WT), JE2Δ*ricO*Δ*spa,* JE2Δ9*ricS*Δ*spa*, JE2*ricO*(R18A)Δ*spa* and JE2*ricO*(ΔAH)Δ*spa*, all of which lack the endogenous, antibody-binding protein A. The ChIP enrichment is shown in 1-kb bins. Data shown are representatives of two to four biological replicates. b) RicO binds a conserved sequence motif named *ricS*. Top, illustration of the genomic regions flanking the replication origin indicating the relative positions of *ricS* sites, as well as of *parS* sites (from ^41^). Bottom, sequence motif generated with the nine *ricS* sites shown in Supplementary Figure 8a. c) Purified RicO binds DNA containing *ricS in vitro*. Electrophoretic mobility shift assay using a 54-bp DNA probe containing the *ricS1* sequence (highlighted in red) and recombinant RicO comprising its DNA-binding and coiled-coil domains (residues 1-174). An otherwise identical DNA fragment containing the indicated base substitutions in the central 18-bp region comprising *ricS1* was used as a negative control (*ricS1\**). RicO wild-type (WT), but not its mutant derivative R18A, bound *ricS1* and *ricS1\**, albeit with lower affinity. d) Deletion of the nine identified *ricS* sites results in chromosome segregation defects, as well as in synthetic sickness in cells lacking a functional SMC complex. The bar graph shows the percentage of anucleate cells measured for the strains (from left to right) JE2, JE2Δ*ricO,* JE2Δ9*ricS,* JE2Δ9*ricS*Δ*ricO,* JE2Δ9*ricS*Δ*ricO*p*ricO,* JE2_Δ*scpAB*, JE2Δ9*ricS*Δ*scpAB* and JE2Δ9*ricS*Δ*scpAB*p*scpAB*. Bars represent the mean and lines indicate the standard deviation of three independent experiments. At least 1279 cells were analysed for each biological replicate. Fluorescence micrographs overlaid with phase contrast images show cells labelled with DNA dye Hoechst 33342 (cyan), and anucleate cells are indicated by white arrowheads. Scale bar, 1 µm.

To provide further evidence for specific DNA binding by RicO, we next analysed the RicO-*ricS* interaction by electrophoretic mobility shift assay (EMSA). Recombinant RicO comprising the predicted DNA-binding domain (DBD) and coiled-coil region showed high affinity for a 54-bp DNA fragment containing the 18-bp *ricS1* motif and its flanking regions (Fig. 3c). When the 18-bp *ricS1* motif was mutated in the 54-bp fragment, RicO binding was impaired, but not completely abolished (Fig. 3c; Supplementary Figure 9a-d). Taken together, these data show that RicO preferentially binds a specific sequence motif located in the origin-proximal region of the chromosome but can also non-specifically bind DNA with lower affinity.

### *ricS* sites support RicOʹs function in chromosome segregation

We next asked if *ricS* sites are required for RicOʹs function in chromosome segregation. To address this question, we sequentially deleted the nine identified *ricS* sites in the JE2 background and studied the effects on anucleate cell fraction. Chromosome segregation defects became apparent after deletion of seven *ricS* sites and progressively increased by deleting the two remaining *ricS* sites (Supplementary Figure 8b). Interestingly, the JE2Δ9*ricS* deletion strain produced anucleate cells at a higher frequency (∼24%) than the JE2Δ*ricO* strain (∼4%) (Fig. 3d) and showed moderately impaired cell growth (Supplementary Figure 2b), suggesting that unbound and/or non-specifically bound RicO is deleterious to the cell. In agreement, deletion of *ricO* in the JE2Δ9*ricS* strain reduced the chromosome segregation defects to JE2Δ*ricO* levels (Fig. 3d). A complementation assay using plasmid-encoded *ricO* increased the frequency of anucleate cells of the JE2Δ9*ricS*Δ*ricO* strain back to nearly JE2Δ9*ricS* levels (Fig. 3d). These data suggest that *ricS* sites are required for RicO to properly function in chromosome segregation.

We then asked if the recruitment of RicO by extra *ricS* sequences would be deleterious. For this, we introduced a 97-bp genome fragment containing *ricS1* and its flanking regions into the multicopy plasmid pEPSA5 ^59^. Introduction of the plasmid pEPSA5-*ricS1*, but not of the empty vector pEPSA5, into the parental strain JE2 caused an increase in the proportion of anucleate cells to levels of the JE2Δ*ricO* strain (∼4%) (Supplementary Figure 10a) without substantially affecting cell growth (Supplementary Figure 2c). Contrarily, introducing pEPSA5-*ricS1* into the JE2Δ*ricO* strain did not notably change the percentage of anucleate cells (Supplementary Figure 10a), indicating that chromosome segregation defects of cells with extra *ricS1* copies were mediated by RicO, presumably due to competition with the native *ricS* sites. These data suggest that introducing *ricS* sites on a plasmid sequesters RicO and perturbs RicO’s function in chromosome segregation.

To our surprise, despite causing chromosome segregation defects, mutants lacking *ricS* sites, as well as those carrying the *ricS*-containing plasmid, maintained polar localizations of *oriC* and RicO^SW^-GFP (Supplementary Figures 8c,d & 10b,c), indicating that *ricS* sites contribute to RicO’s function in chromosome segregation but are dispensable for RicO-dependent localization of replication origins.

We next studied the effect of deleting the nine *ricS* sites on anucleate cell fraction of strains lacking ParB and ScpAB. The JE2Δ9*ricS*Δ*scpAB* strain showed a drastic increase in the proportion of anucleate cells (∼58%) relative to the JE2Δ9*ricS* (∼24%) and JE2_Δ*scpAB* (∼17%) strains (Fig. 3d), as well as substantially reduced cell growth (Supplementary Figure 2b). The anucleate cell proportion of the JE2Δ9*ricS*Δ*scpAB* strain was reduced to JE2Δ9*ricS* levels by complementation with plasmid-borne *scpAB* (Fig. 3d). Contrarily, deleting *parB* in the JE2Δ9*ricS* strain reduced the fraction of anucleate cells from ∼23% to ∼9% (Supplementary Figure 4b). Addition of extra RicO binding sites also impacted cells lacking ParB and ScpAB. Introduction of plasmid-borne *ricS1* in the JE2_Δ*scpAB* strain resulted in a doubling of anucleate cells (∼33%) relative to the empty vector control (∼15%) (Supplementary Figure 10a), while the JE2_Δ*parB* strain was less affected by extra *ricS* copies than the JE2 wild-type (1.4±0.2% vs. 3.9±0.2% of anucleate cells, respectively) (Supplementary Figure 4c). In sum, chromosome segregation defects of cells with changes in the number of *ricS* sites were exacerbated by *scpAB* deletion and alleviated by *parB* deletion. This further suggests that *ricS* sites have a role in RicO function, likely underpinning the interplay of RicO, ParB and the SMC complex.

### RicO contributes to origin organization

Our data indicate that RicO-*ricS* acts synergistically with the SMC complex in chromosome segregation (Fig. 1a; Fig. 3d). The SMC complex structurally organizes the chromosome and facilitates juxtaposition of chromosome arms through ParB-dependent loading at origin-proximal *parS* sites ^20-23,25,27^. Hence, we asked if RicO also has a role in chromosome organization. To address this question, we analysed genome-wide DNA-DNA interactions for the JE2Δ*ricO* strain using high-throughput chromosome conformation capture by Hi-C. Hi-C contact maps of JE2 wild-type cells (Fig. 4a) showed the characteristic primary and secondary diagonals, corresponding to short-range interactions between genomically adjacent regions and distant interactions between the two chromosome arms, respectively, as well as additional contacts of origin-flanking regions, as previously described ^41,60-62^. Strikingly, the JE2Δ*ricO* strain showed reduced DNA contacts in the origin-proximal portion of the chromosome (yellow arrows in Fig. 4) and dramatically increased interactions between origin and the rest of the chromosome (black arrows in Fig. 4b), indicating that RicO insulates the origin region from the rest of the chromosome. DNA-DNA interaction patterns surrounding the *oriC* region were restored to wild-type levels by complementation of the JE2Δ*ricO* strain with plasmid-encoded *ricO* (Fig. 4a). These signals were still observed in Hi-C maps for the JE2_Δ*parB* and JE2_Δ*scpAB* strains but were lost upon deletion of *ricO* in these mutants, confirming that *oriC*-proximal interactions are facilitated by RicO and not by ParB or the SMC complex (Supplementary Figure 11). Origin-proximal DNA-DNA interactions were weakened in the JE2Δ9*ricS* strain (Supplementary Figure 12), but not to the extent of JE2Δ*ricO,* indicating that the role of RicO in structurally organizing the origin-proximal region of the chromosome is not totally dependent on specific DNA binding.

**Figure 4.**
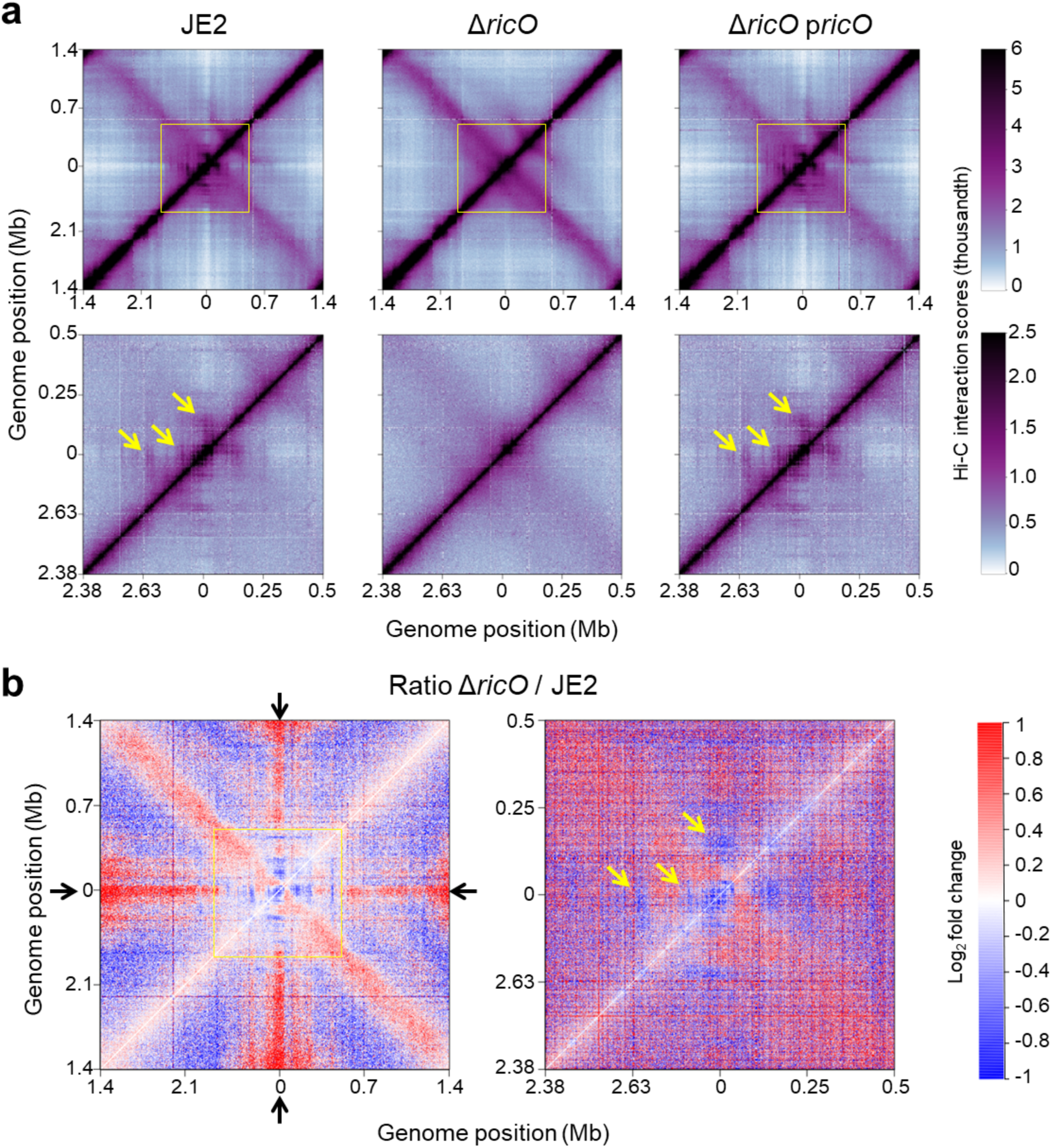
Cells lacking RicO show changes in origin organization. a) RicO facilitates DNA-DNA interactions in the origin-proximal (∼700-kb) region of the chromosome. Normalized Hi-C contact maps of the strains JE2 (from ^41^), JE2Δ*ricO* (middle) and JE2Δ*ricO*p*ricO* (right) displaying contact frequencies for pairs of 5-kb bins. Contact maps in the lower row show a zoom into the 1-Mb region flanking the replication origin (highlighted by yellow squares in the corresponding whole-genome maps) and display contact frequencies for pairs of 1-kb bins. Yellow arrows indicate DNA contacts in the origin-proximal region facilitated by RicO. Data obtained from two biological replicates were combined into one contact map for each strain. b) Ratio plots of the Hi-C contact maps shown in panel a. Red and blue indicate increased and reduced chromosomal interactions, respectively, detected in the strain JE2Δ*ricO* relative to the JE2 wild-type strain. Black arrows point to the vertical and horizontal lines that result from increased interactions of the origin region with the rest of the chromosome in the absence of RicO.

Ratio plots of the Hi-C contact maps of JE2Δ*ricO* versus JE2 showed that short-range interactions (the primary diagonal running from bottom left to top right) were not markedly changed (Fig. 4b), indicating that RicO does not contribute to the local packaging of the chromosome. Finally, in Δ*ricO* cells, SMC-dependent inter-chromosomal arm interactions (the secondary diagonal running from top left to bottom right) were increased, suggesting that RicO may reduce SMC translocation beyond the origin region. Given that *parB* deletion in the JE2Δ*ricO* strain completely abolished long-range interactions between chromosome arms but reduced the frequency of anucleate cells (Supplementary Figures 4a & 11), our data are consistent with a model in which RicO reduces the chance of ParB-loaded SMC to travel beyond the origin region, thereby concentrating SMC near the origin to facilitate chromosome segregation.

### RicO overproduction causes aberrant nucleoid shape

To characterize the effect of an excess of RicO, we generated a *ricO* overexpression construct based on the vector pEPSA5 ^59^ and the constitutive endogenous *amaP* promoter ^63^. P*amaP*-driven ectopic expression of *ricO* from a multicopy plasmid in the JE2*ricO*OE (overexpression) strain resulted in approximately 1.5- to 2-fold increased cellular abundance of RicO relative to the empty vector control (Supplementary Figure 7c). Plasmid-borne *ricO* in the JE2*ricO*OE strain caused a high proportion of anucleate cells (∼7%) and reduced cell growth (Fig. 5a; Supplementary Figure 2c). In Hi-C assays, *ricO*-overexpressing cells showed an increase in DNA-DNA interactions in the origin-proximal region of the chromosome, while inter-chromosomal arm interactions (the secondary diagonal running from top left to bottom right) were reduced relative to the empty vector control (Fig. 5b). Notably, *ricO* overexpression and *ricO* deletion affected the same genomic regions but altered their interactions in opposite ways.

**Figure 5.**
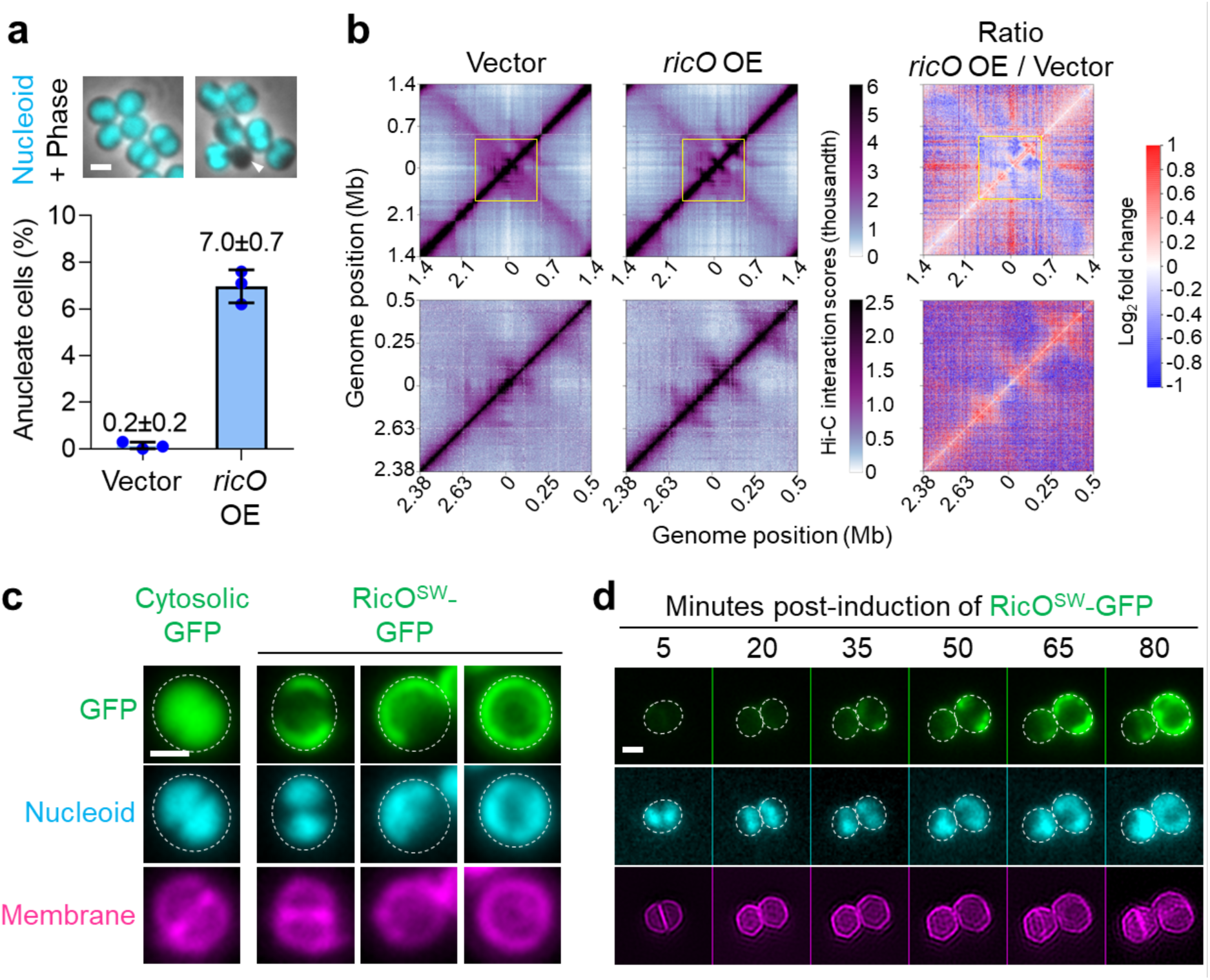
Excess RicO causes anucleate cells and misshaped nucleoids. a) Overexpression of *ricO* results in chromosome segregation defects. Bar graph shows the percentage of anucleate cells measured for the strains JE2vector (empty vector control, left) and JE2*ricO*OE (overexpressing *ricO*, right). Bars represent the mean and lines indicate the standard deviation of three independent experiments. At least 777 cells were analysed for each biological replicate. Fluorescence micrographs overlaid with phase contrast images show cells labelled with DNA dye Hoechst 33342 (cyan), and anucleate cells are indicated by white arrowheads. Scale bar, 1 µm. b) Overexpression of *ricO* alters genomic interactions in the origin-proximal region of the chromosome. Normalized Hi-C contact maps of the strains JE2vector (left) and JE2*ricO*OE (middle) displaying contact frequencies for pairs of 5-kb bins. Contact maps in the lower row show a zoom into the 1-Mb region flanking the replication origin (highlighted by yellow squares in the corresponding whole-genome maps) and display contact frequencies for pairs of 1-kb bins. Data obtained from a single biological replicate is shown for each strain. Ratio plot on the right shows increased (red) and reduced (blue) chromosomal interactions detected in the strain JE2*ricO*OE relative to the JE2vector strain. c) Increased cellular levels of RicO result in aberrant nucleoid shape. Fluorescence micrographs of the strains JE2pBCB50*gfp* (Cytosolic GFP) and JE2pBCB50*ricO^SW^-gfp* (RicO^SW^-GFP) grown in the presence of the inducer Atc to induce production of GFP and RicO^SW^-GFP (both in green), respectively. Cells were stained with the DNA dye Hoechst 33342 (cyan) and the membrane dye FM5-95 (magenta). Cell outlines inferred from stained membranes are indicated by dashed white ovals. Images are representative of two biological replicates. Scale bar, 0.5 µm. d) Formation of misshaped nucleoids coincides with intracellular accumulation of RicO. Fluorescence micrographs show cells of the strain JE2pBCB50*ricO^SW^-gfp* induced with Atc and imaged by time-lapse microscopy. Cells were stained with the DNA dye Hoechst 33342 (cyan) and the membrane dye CellBrite Fix 640 (magenta). Formation of aberrantly shaped nucleoids was observed when cells showed an incremental increase in RicO^SW^-GFP fluorescence (green). Cell outlines inferred from stained membranes are indicated by dashed white ovals. Images are representative of two biological replicates. Scale bar, 1 µm.

To better understand the chromosome segregation defects of *ricO*-overexpressing cells, we localized overproduced RicO^SW^-GFP in single cells. RicO^SW^-GFP, and cytoplasmic GFP as a control, were produced from a construct based on the vector pCN51 ^64^ and the anhydrotetracycline (Atc)-inducible *xyl-tetO* promoter ^65^. Addition of Atc to cells of the JE2pBCB50*ricO^SW^-gfp* strain resulted in RicO^SW^-GFP accumulating at the cell poles and in the cell periphery, whereas non-tagged GFP overproduced in the JE2pBCB50*gfp* strain localized diffusively (Fig. 5c). RicO^SW^-GFP localization near the cell membrane was accompanied by an uneven cellular distribution of the nucleoid, which appeared to be sequestered to sites of highest RicO^SW^-GFP accumulation (Fig. 5c). Using time-lapse microscopy, we observed the formation of misshaped and unevenly distributed nucleoids coinciding with the accumulation of RicO^SW^-GFP at the cell membrane (Fig. 5d). These data confirmed that RicO is associated with the nucleoid and suggest that it can bind DNA non-specifically.

### Conserved domains of RicO are essential for its function and localization

Bioinformatic analysis of the RicO sequence predicted the presence of an N-terminal DBD and a putative membrane-binding C-terminal amphipathic helix (AH) (Supplementary Figure 6a). Structural modelling of RicO in complex with DNA indicated that positively charged arginine at position 18 located in the predicted DNA interface may directly facilitate interaction (Supplementary Figure 6b). To impair DNA and membrane binding by RicO, we replaced the native *ricO* gene with mutant variants encoding either arginine 18 to alanine substitution (R18A) or RicO truncated by 14 residues encompassing its AH (ΔAH), respectively. None of these two mutations led to major changes in cellular RicO levels (Supplementary Figure 7c). As expected, ChIP-seq for the JE2*ricO*(R18A)Δ*spa* strain resulted in loss of the RicO enrichment peaks, confirming loss of specific binding to *ricS* sites, while the enrichment profile for JE2*ricO*(ΔAH)Δ*spa* was similar to that of the parental strain JE2Δ*spa* (Fig. 3a). The R18A mutation also abolished the ability of RicO to bind DNA *in vitro* (Fig. 3c; Supplementary Figure 9e), confirming that DNA binding by RicO is conferred by its DBD. Notably, unlike RicO wild-type protein, the RicO(R18A) mutant also failed to bind non-*ricS* DNA (Fig. 3c). Similar to the JE2Δ*ricO* strain, both JE2*ricO*(R18A) and JE2*ricO*(ΔAH) strains showed an increased proportion of anucleate cells (∼4% and ∼5%, respectively) and weakened genomic interactions in the *oriC* region relative to the JE2 wild-type strain (Fig. 6a,c; Supplementary Figure 11), indicating that RicO binding to both DNA and membrane is required for its function. Moreover, both JE2*oriC*^mNG^*ricO*(R18A) and JE2*oriC*^mNG^*ricO*(ΔAH) strains showed loss of polar *oriC* localization, similar to Δ*ricO* cells (Fig. 6b). Notice that while abolished RicO DNA binding by the R18A mutation resulted in loss of polar *oriC* localization, deletion of nine *ricS* sites did not (Supplementary Figure 8c), confirming that RicO binding to DNA is required for origin positioning, but this binding does not have to be specific. However, specific DNA binding by RicO is required for proper chromosome segregation.

**Figure 6.**
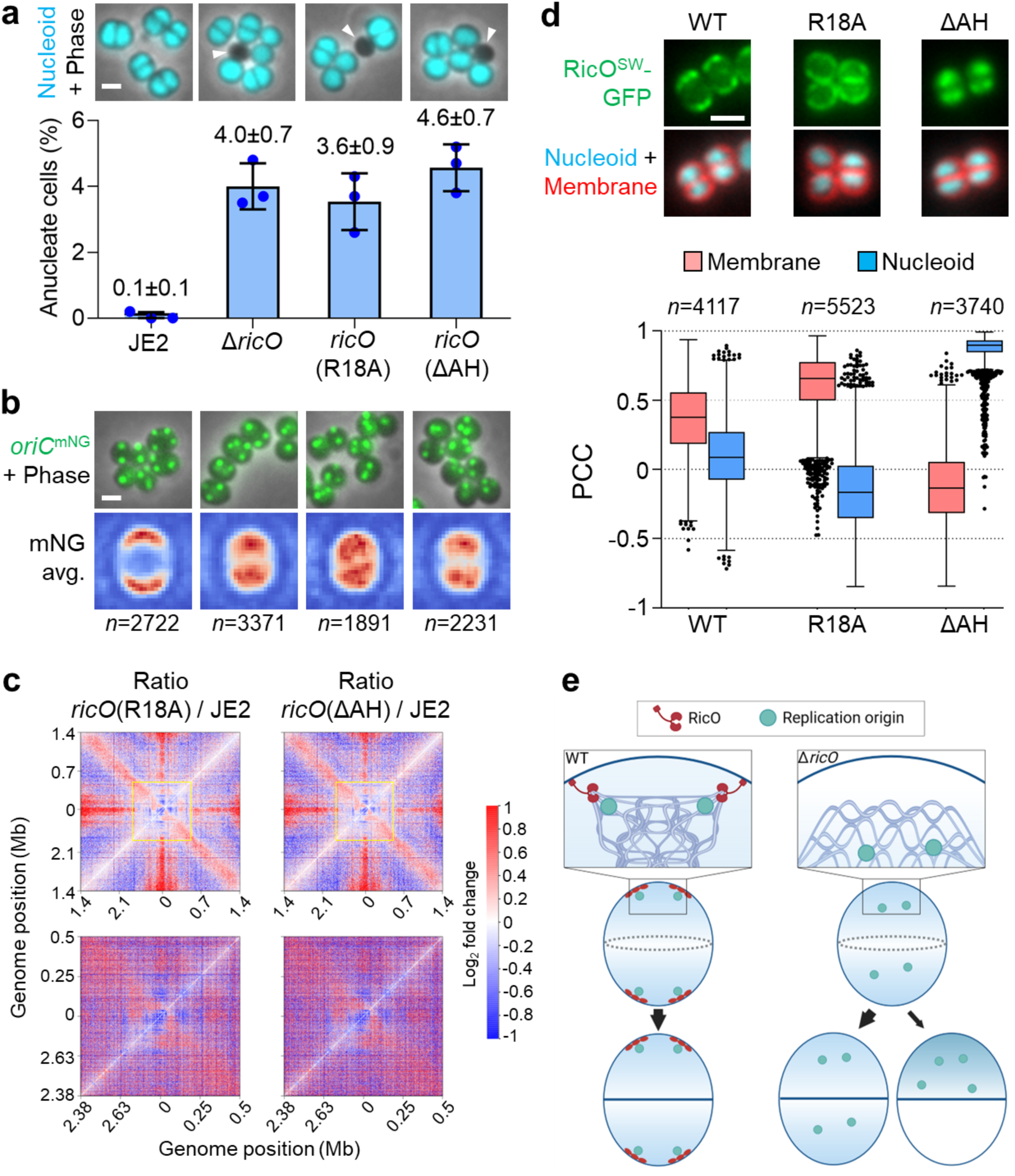
The DNA-binding domain and amphipathic helix of RicO are both required for its function and localization. a) Mutations targeting the DNA-binding domain and amphipathic helix of RicO result in chromosome segregation defects. The bar graph shows the percentage of anucleate cells measured for the strains (from left to right) JE2, JE2Δ*ricO,* JE2*ricO*(R18A) and JE2*ricO*(ΔAH). Bars represent the mean and lines indicate the standard deviation of three independent experiments. At least 882 cells were analysed for each biological replicate. Fluorescence micrographs overlaid with phase contrast images show cells labelled with DNA dye Hoechst 33342 (cyan), and anucleate cells are indicated by white arrowheads. Scale bar, 1 µm. b) Mutations targeting the DNA-binding domain and amphipathic helix of RicO result in loss of polar localization of chromosomal origins. Fluorescence micrographs overlaid with phase contrast images of the strains (from left to right) JE2_FROS^Ori^, JE2*oriC*^mNG^Δ*ricO,* JE2*oriC*^mNG^*ricO*(R18A) and JE2*oriC*^mNG^*ricO*(ΔAH). Replication origins (green) were fluorescently labelled by producing TetR-mNG and introducing a *tetO* array at position 358° of the circular genome. Lower row shows heat maps of the average localization of detected fluorescent foci corresponding to replication origins of the same strains as cells shown in the upper row. Blue-white-red color code in heat maps represents spot density from low to high, respectively. *n*, number of analysed cells. Scale bar, 1 µm. c) Mutations targeting the DNA-binding domain and amphipathic helix of RicO reduce origin-proximal genomic interactions similarly to deletion of *ricO* (compare with Figure 4b). Ratio plots of normalized Hi-C contact maps of the strains JE2, JE2*ricO*(R18A) and JE2*ricO*(ΔAH) displaying contact frequencies for pairs of 5-kb bins. Red and blue indicate increased and reduced chromosomal interactions, respectively, detected in the strains JE2*ricO*(R18A) and JE2*ricO*(ΔAH) relative to the JE2 wild-type strain. Contact maps in the lower row show a zoom into the 1-Mb region flanking the replication origin (highlighted by yellow squares in the corresponding whole-genome maps) and display contact frequencies for pairs of 1-kb bins. Data obtained from two biological replicates were combined into one contact map for the JE2 and JE2*ricO*(R18A) strains, while a single biological replicate was analysed for the JE2*ricO*(ΔAH) strain. d) Mutations of RicOʹs DNA-binding domain and amphipathic helix cause distinct changes in its localization. Fluorescence micrographs show RicO^SW^-GFP and its mutant derivatives (green) localized in the strains JE2*ricO^SW^-gfp* (WT), JE2*ricO*(R18A)*^SW^-gfp* and JE2*ricO*(ΔAH)*^SW^-gfp*, each stained with the DNA dye Hoechst 33342 (cyan) and the membrane dye FM5-95 (red). Pearson Correlation Coefficient (PCC) was used as a measure of colocalization of RicO^SW^-GFP variants with stained membrane or DNA. A PCC value close to one denotes colocalization, whereas a negative value denotes anti-colocalization. Data are represented as box-and-whisker plots in which boxes correspond to the first-to-third quartiles, lines inside the boxes indicate the median, and the ends of whiskers and outliers follow a Tukey representation. Data shown are from three biological replicates. Scale bar, 1 µm. e) Model for RicO associating chromosomal origins with the cell poles in *S. aureus*. In this model, RicO directly binds the cell membrane via its amphipathic helix and the origin-proximal portion of the chromosome via its DNA-binding domain, resulting in polar positioning and structural organization of replication origins. Origin-anchoring by RicO is required to ensure that, upon cell division, each cell acquires a complete copy of the chromosome.

To study the effect of abolished DNA or membrane binding on RicO localization, we introduced the R18A and ΔAH mutations into the JE2*ricO^SW^-gfp* strain, producing RicO^SW^-GFP from its native genomic locus, without causing changes in the stability of the fluorescent mutant derivatives (Supplementary Figure 7b). The two mutations had distinct effects on the localization of RicO^SW^-GFP: whereas the R18A mutation resulted in a redistribution of RicO(R18A)^SW^-GFP around the peripheral membrane, RicO(ΔAH)^SW^-GFP colocalized with the nucleoid and further away from the membrane compared with wild-type RicO^SW^-GFP (Fig. 6d). Taken together, these data are consistent with a model in which RicO binds the nucleoid and the cell membrane through its DBD and AH, respectively, which is strictly required for RicO’s function in origin organization and chromosome segregation.

### Distribution of RicO-like proteins in the Firmicutes

The structure prediction of RicO, together with its role in chromosome positioning, suggests an evolutionary relationship with RocS, a protein promoting the segregation of newly replicated origins in *S. pneumoniae* ^66^, and RacA from *B. subtilis* ^46,47,50^, despite only 26% of sequence identity. All three proteins are predicted to have a small helix-turn-helix DBD, often annotated as belonging to the MerR/MarR domain families, followed by a coiled-coil domain, while only RicO and RocS contain a predicted C-terminal AH (Supplementary Figure 6a). Moreover, the three proteins are involved in anchoring chromosomes, and we thus decided to explore their possible phylogenetic relationship and their taxonomic distribution in other bacteria.

We searched for homologs of RicO, RacA and RocS in a local database containing 500 genomes of Firmicutes (Bacillota) using a Hidden Markov Model (HMM) built from these three proteins and close homologs. Using an e-value cut-off of 10^-4^, we retrieved 137 sequences, which we used to infer a maximum likelihood tree (Supplementary Figure 13). For these, we used the prediction tool CoCoNat ^67^ to predict the presence of coiled-coil regions and their oligomeric state. In the maximum likelihood tree, we observed a distinct clade containing only sequences of proteins predicted to form parallel dimers (rarely present in the rest of the tree), and we decided to tentatively root the tree between this clade and another clade containing the rest of the sequences. In the clade containing parallel dimers, we found the sequences belonging to Lactobacillales (including RocS), Clostridia and Erysipelotrichia, in agreement with a previous report on RocS forming parallel dimers ^68^. Sequences from Clostridia can also be found outside of this clade, in some cases predicted to have coiled-coil regions forming antiparallel dimers. However, these belong to species absent in the parallel dimer clade, except for *Blautia hydrogenotrophica*, which has two homologs, one in the clade of the parallel dimers and one outside it. In contrast, the sequences belonging to the order Bacillales (including RicO and RacA) are predicted to bear coiled-coil regions that form tetramers or trimers. This could indicate that the ancestral version of the protein (before the Bacillales-Lactobacillales split) formed parallel dimers, but the Bacillales ancestor already carried a tetrameric/trimeric version. We can speculate that this change affected the properties of the protein, particularly its interaction with DNA: RocS (dimer) is thought to bind sequences based on their topology ^66^, whereas RicO and RacA (tetramers) can bind specific DNA sequences (Fig. 3) ^46,48^. In conclusion, this analysis supports that RicO, RacA and RocS form part of a family of proteins widespread among the Firmicutes. Future work characterizing more proteins from this family will allow us to shed light on their evolutionary history, how it affected their roles in chromosome positioning, and their distribution inside and outside of the Firmicutes.

## Discussion

Bacterial chromosomes undergo dynamic changes in condensation and conformation during the cell cycle and development ^69^. Chromosome segregation in some rod-shaped bacteria involves specialised systems for remodelling and anchoring the chromosomes to the cell poles ^47,51-53^. In this work, we identified a previously uncharacterized protein in the coccoid bacterium *S. aureus*, named RicO, and show it has an important role in anchoring of the staphylococcal chromosome (Fig. 6e). RicO can bind both specifically and non-specifically to the *S. aureus* chromosome. Specific binding occurs at nine *ricS* sites with a 18-bp conserved motif, distributed within ∼700 kb (equivalent to ∼25% of the genome) of the origin-proximal portion of the chromosome. The *ricS* motif has a balanced GC-content and does not contain a repeat or palindrome, thus it does not resemble the centromere-like binding sequence *ram* in *B. subtilis* ^46^ or other *cis*-acting elements for chromosome segregation in *B. subtilis* ^70^ and *Escherichia coli* ^71^. The *B. subtilis* developmental protein RacA preferentially binds 25 *ram* sites within a ∼600 kb flanking region of the replication origin, as well as in a dispersed manner throughout the chromosome, causing a high degree of DNA compaction ^46^. Contrary to the location of *ram* sites inside protein-coding sequences, all nine *ricS* sites mapped to intergenic regions. Interestingly, one of the *ricS* sites (*ricS5*) is located in the upstream region of *ricO*, the last gene of a predicted polycistronic transcriptional unit. However, a strain lacking all nine *ricS* sites produced RicO at about wild-type levels, suggesting that RicO binding does not significantly affect its gene expression or stability. This strain had chromosome segregation defects that, surprisingly, were more severe than those resulting from the absence of RicO, suggesting that specific DNA binding may constitute a mechanism to sequester RicO at specific regions in the genome, thereby keeping its activity in check.

Unexpectedly, RicO-mediated positioning and structural organization of chromosomal origins does not only rely on its specific binding to *ricS* sites. Deletion of all nine identified *ricS* sites maintained polar localization of replication origins and most of RicO-dependent origin-proximal genomic interactions, despite causing severe chromosome segregation defects (Fig. 3d; Supplementary Figures 8c & 12). Loss of polar localization and structural organization of replication origins was only observed when RicO binding to DNA was completely abolished by amino acid substitution R18A in the predicted N-terminal DNA-binding domain (DBD). This suggests that wild-type RicO was able to bind non-specific DNA *in vitro*, albeit with lower affinity when compared to specific binding to *ricS* (Fig. 3c; Supplementary Figure 9). We provide three lines of evidence for the ability of RicO to bind non-specific DNA *in vivo*: (i) deletion of all nine *ricS* sites to abolish specific DNA binding by RicO resulted in weak DNA enrichment by ChIP in the origin-proximal portion of the chromosome, which was not observed for cells lacking RicO or producing RicO(R18A) (Fig. 3a); (ii) removal of RicOʹs amphipathic helix (AH) to abolish membrane binding resulted in its colocalization with the nucleoid, suggesting scattered binding of RicO across the chromosome (Fig. 6d); (iii) accumulation of overproduced RicO^SW^-GFP at the peripheral membrane coincided in space and time with the formation of misshaped nucleoids, suggesting that RicO is capable of associating entire chromosomes with the cell membrane (Fig. 5c,d). The ability of RicO to bind non-specific DNA could explain why polar localization and organization of origins were maintained in the Δ9*ricS* mutant. Interestingly, structural analysis with MADOKA ^72^ revealed high similarity between the predicted N-terminal DBD of RicO and the helix-turn-helix domain of *B. subtilis* RacA required for DNA binding (Supplementary Figure 6b). *B. subtilis* RacA forms a stable complex even with non-specific DNA, which is required for the remodelling of the chromosomes into an axial filament during sporulation ^46^. The mode of RacA binding to DNA is unique in the sense that the same residues can be adjusted to bind DNA either specifically or non-specifically. Arginine 19 in RacA is an adaptable base and phosphate-interacting residue that can make bipartite contacts by recognising the *ram* site and by interacting with phosphates in non-*ram* DNA through side chain rotations. Similar to the R18A mutation in RicO, the equivalent mutation R19A in RacA completely abolished DNA binding ^48^. Based on their high structural similarity, we speculate that RicO and RacA employ a similar adjustable DNA docking mode and mechanism for DNA compaction. RacA was reported to cause strong DNA compaction depending on self-interaction presumably through its coiled-coil structures in its C-terminal region ^46^. Likewise, RicO molecules bound to DNA might achieve the high degree of *oriC*-proximal DNA-DNA interactions by interacting with each other through the central coiled-coil domain to form a single, highly condensed, higher order structure.

Reminiscent of RicO, ParB binds to multiple sites clustered in the origin region of the chromosome. It has been previously shown that *S. aureus* ParB binds to five 16-bp *parS* sites distributed over ∼400 kb flanking the replication origin ^5,41^. In *S. aureus*, cells lacking ParB show normal chromosome segregation and polar localization of replication origins ^41^, hence no direct evidence exists for ParB to actively segregate origins or act as an anchor for the origin region in this organism, which encodes no ParA homolog. ParB promotes the co-linearity of chromosome arms in *S. aureus* ^41,62^, presumably by loading SMC complexes onto the chromosome at origin-proximal *parS* sites ^20-23,25,27^. Our data indicate that the chromosome segregation defects observed for cells with impaired RicO function were partially dependent on ParB since detrimental effects caused by *ricO* deletion, multiple *ricS* deletions or extra *ricS* copies were partially mitigated by *parB* deletion (Supplementary Figure 4). Based on our finding that inter-chromosomal arm interactions were increased in cells lacking RicO, we can envision two possible hypotheses. First, in the absence of RicO, ParB might load SMC complexes at an abnormally high rate. The clustering of *parS* and *ricS* sites in the origin-proximal portion of the chromosome (Fig. 3b) may generate an overlap between ParB-occupied genomic regions and *ricS* sites, thus generating a potential conflict between RicO and ParB for DNA binding. In *S. aureus*, the average genomic distance from *parS* to its closest *ricS* site is 12 kb, with the smallest distance of 5 kb between the *ricS-parS* pair located nearest to the origin. Given that *S. aureus* ParB slides over DNA to occupy ∼8-10 kb of the genome to form a large nucleoprotein complex ^41^, while the genomic regions occupied by RicO were approximately 0.25 kb in size, ParB could spread into *ricS* sites, but RicO likely does not spread into *parS* sites. It is tempting to speculate that RicO limits ParB sliding on the DNA and thereby limits the loading of SMC complexes at *parS* sites. Secondly, RicO tethering the chromosome to the membrane might simply serve as a barrier for SMC translocation. This will allow SMC to concentrate near the replication origin to help origin segregation, rather than allowing SMC to rampantly translocate down the chromosome arms. Since *S. aureus* is a coccus rather than a rod, origin-localized SMC might be more effective at segregating the chromosome than the aligned chromosome arms.

An interesting remaining question is which factors determine the localization of RicO at the cell poles. The requirement for RicO to directly associate with the cell membrane contrasts with that of *B. subtilis* RacA, which is recruited to the membrane at the cell poles by DivIVA. DivIVA forms a meshwork at negatively curved membranes recognized by its N-terminal domain ^49,73^. Our results indicate that both the conserved DBD and amphipathic helix (AH) are required for RicO assembly into foci at the cell poles. The RicO(R18A) mutant deficient in DNA binding localized in a dispersed manner at the peripheral membrane, suggesting that the chromosome itself may have a role in concentrating RicO at the cell poles. On the other hand, neither multiple deletions nor extra copies of *ricS* sites resulted in sequestration of RicO away from the cell membrane. RicO’s association with the cell membrane is facilitated by its C-terminal AH. This region shares high similarity to the AH domains of *S. pneumoniae* RocS ^66^, *E. coli* MinD ^74^ and the *S. aureus* plasmid partitioning protein Par ^75^. These proteins are thought to be inserted into the interior of the bilayer through the hydrophobic surface of their AH. RocS’s AH forms a kink-helix motif that mediates specific lipid interactions, and a dimer is sufficient to facilitate membrane association, but it does not confer specificity for localization and can be substituted with unrelated transmembrane domains ^68,76^. Our data indicate that membrane binding by RicO is crucial for its function in chromosome segregation, as a RicO mutant variant lacking its AH was unable to associate with the membrane and hence failed to correctly position chromosomal origins at the cell poles. Replication origins associated with the membrane by RicO likely have reduced ability to interact with physically distant loci of the longitudinally configured chromosome, which might contribute to their insulation. Interestingly, RicO lacking its AH also failed to structurally organize the replication origin region while maintaining its ability to bind *ricS* sites (Figs. 3a & 6c). This highlights that DNA binding alone is not sufficient and that membrane association by RicO is required for anchoring and insulating chromosomal origins.

The predicted structure of RicO is reminiscent of that of RocS (Supplementary Figure 6), a protein required for proper chromosome partitioning in *S. pneumoniae* ^66^. RocS’ MerR-type winged helix-turn-helix facilitates interaction with DNA and the coiled-coil region mediates self-interaction, which in turn is required for association with the membrane via the AH ^68^. RocS is involved in the initial segregation of newly replicated origins from mid-cell towards the equatorial rings of *S. pneumoniae*, a function that becomes essential in cells lacking either ParB or SMC ^66^. In *S. aureus*, synthetic sickness was caused only by simultaneous removal/inactivation of RicO and SMC, but not of RicO and ParB (Fig. 1a; Supplementary Figure 4a). Unlike RocS in *S. pneumoniae*, RicOʹs function in chromosome segregation of *S. aureus* does not seem to employ an active mechanism for separating newly replicated origins. The number of diffraction-limited foci corresponding to replication origins remained unchanged in cells lacking RicO and the segregation of RicO foci did not follow a specific spatiotemporal choreography with origins (Fig. 2f; Supplementary Figure 5). Both *S. aureus* and *S. pneumoniae* lack a ParA homolog and a Min system, while only *S. aureus* encodes a functional homolog of the nucleoid occlusion protein Noc ^39,56,77^. Despite shared features and structural similarity, we propose that *S. aureus* RicO and *S. pneumoniae* RocS have evolved distinct functions in chromosome segregation. Whereas RocS protects the nucleoid from premature truncation during cell division in *S. pneumoniae*, similarly to Noc in *S. aureus*, the primary role of RicO is to structurally organize and facilitate polar localization of chromosomal origins.

In summary, our study shows that the coccoid bacterium *S. aureus* employs a dedicated mechanism for anchoring chromosomal replication origins at the cell periphery. We demonstrate that this mechanism is based on the ability of newly identified RicO to directly interact with both the nucleoid and the cell membrane, acting as a bridge between the two. In addition to the condensin-like SMC complex, RicO serves as a major factor for chromosome segregation in *S. aureus*.

## Methods

### Bacterial growth conditions

Strains and plasmids used in this study are listed in Supplementary Tables 1 and 2. Plasmids containing DNA sequences from *S. aureus* were designed based on information from the strain-specific genome browser of *Aureo*Wiki ^78^. *E. coli* strains were grown on Luria-Bertani agar (VWR) or in Luria-Bertani broth (VWR) with aeration at 37 °C. *S. aureus* strains were grown on tryptic soy agar (VWR) or in tryptic soy broth (TSB, Difco) at 200 rpm with aeration at 37 °C or at 30 °C. When necessary, culture media were supplemented with antibiotics (100 μg/ml ampicillin, Sigma-Aldrich; 10 μg/ml erythromycin, Apollo Scientific; 10 μg/ml chloramphenicol, Sigma-Aldrich). 5-bromo-4-chloro-3-indolyl β-d-galactopyranoside (X-Gal, Apollo Scientific) was used at 100 μg/ml and anhydrotetracycline (Atc, Sigma-Aldrich) was used at 50 ng/ml.

### Construction of *S. aureus* strains

Oligonucleotides used in this study were purchased from Metabion and Integrated DNA Technologies and are listed in Supplementary Table 3. Cloning of RicO mutants and tagged protein fusions in *S. aureus* was done using the following general strategy: plasmids were propagated in *E. coli* strain DC10B and purified from overnight cultures supplemented with the relevant antibiotics. Plasmids were then introduced into electrocompetent *S. aureus* RN4220 cells as previously described ^79^ and transduced to JE2, COL or NCTC8325-4 using phage 80α ^80^. Antibiotic marker-free allelic replacements in the *S. aureus* chromosome were performed using plasmids that allow for double homologous recombination events at selected genome sites. Constructs were confirmed by polymerase chain reaction (PCR) and by Sanger sequencing.

Strains with deletions of *ricO, ricO*(AH) and *ricS* were constructed using the pMAD ^81^ and pIMAY-Z ^82^ vectors that allow for double homologous recombination events at selected genome sites. 600-bp upstream and downstream regions of *ricO* from the JE2 genome were amplified using the primer pairs 6983/6984 and 6981/6982, followed by cloning both fragments into SmaI-digested pMAD via Gibson Assembly (NEB), resulting in pMAD-Δ*ricO*. To delete 14 codons at the 3ʹ-end of the native *ricO* gene, a 687-bp upstream region of *ricO* codon 305 and a 753-bp downstream region of *ricO* codon 304 from the JE2 genome were amplified using the primer pairs 10687/10766 and 10679/10676, respectively. Note that primer 10766 contained a stop codon for the termination of *ricO* translation after codon 304. The upstream and downstream regions were digested with EcoRI/BamHI and BamHI/SalI, respectively, joined by ligation and the full constructs were cloned into EcoRI/SalI-digested pMAD, resulting in pMAD-*ricO*(ΔAH). To delete genomic *ricS* sites with a length of 18 bp, approximately 600-800-bp flanking regions from the JE2 genome (JE2*ricO^SW^-snap* genome for the *ricS5* downstream region) were amplified using two primer pairs for *ricS1* (9951/9952; 9953/9954), *ricS2* (10878/10879; 10880/10881), *ricS3* (11245/11246; 11247/11248), *ricS4* (9963/9964; 9965/9966), *ricS5* (11249/11250; 11251/10771), *ricS6* (10882/11243; 11244/10885), *ricS7* (9959/9960; 9961/9962), *ricS8* (11253/11254; 11255/11256), and *ricS9* (9955/9956; 9957/9958). Upstream regions were digested with BamHI/BglII and downstream regions were digested with BglII/EcoRI (BglII/SalI for *ricS5* and *ricS8*, BglII/SmaI for *ricS7*), and corresponding fragments were joined by ligation and the full constructs were cloned into pMAD pre-digested with BamHI/EcoRI (BamHI/SmaI for *ricS7* and BamHI/SalI for *ricS8*) and pIMAY-Z pre-digested with BamHI/XhoI (for *ricS5*). The procedure to generate pMAD-Δ*ricS5*Δ*ricO* was the same as that for pMAD-Δ*ricO*, except that for the former the template DNA to amplify the upstream region of *ricO* was the JE2Δ9*ricS* genome. pMAD and pIMAY-Z derivatives were electroporated into RN4220 and transduced to JE2, COL and NCTC8325-4 derivative strains. Integration excision by a double homologous recombination event was performed at 30 °C and targeted genomic loci were amplified by PCR. Clones with *ricS* and *ricO*(AH) deletions were identified by digestion of the PCR product with BglII and BamHI, respectively, followed by sequencing to verify the absence of deleted sequences.

RicO(R18A) mutant strains were constructed using the pIMAY-Z vector ^82^. A *ricO* allele encoding the point mutation R18A (by substitution of CGT for GCT at bases 52-54) was generated by amplifying a 562-bp upstream and a 922-bp downstream region of codon 18 from the JE2 and JE2*ricO^SW^-snap* genomes using the primers 10675/10163 and 10162/10771, respectively. The two fragments were joined by overlap PCR using the primers 10675/10771, digested with EcoRI/SalI and cloned into EcoRI/XhoI-digested pIMAY-Z. The resulting plasmid pIMAY-Z-*ricO*(R18A) was electroporated into RN4220 and transduced to JE2. Integration excision by a double homologous recombination event was performed at 30 °C and the *ricO* genomic region was sequenced to confirm the presence of the R18A mutation.

Strains producing RicO^SW^-mCherry, RicO^SW^-SNAP, RicO^SW^-GFP, ScpA-mNG and ScpB-mCherry fusions from the native genomic locus were constructed using the pMAD vector ^81^. To insert protein tags into the genomic *ricO* locus, first a 486-bp upstream region of *ricO* codon 183 and a 411-bp downstream region of *ricO* codon 182 from the JE2 genome were amplified using the primer pairs 9436/10684 and 10685/10678, respectively. Secondly, the *mcherry*, *gfp* and *snap-tag* coding sequences from pMAD-rodAmCh ^83^, pMAD-ezrAsgfp ^83^ and pSNAP-tag (T7)-2 (NEB) were amplified using the primer pairs 10888/10889, 10886/10887 and 9630/9632, respectively. Sequences encoding protein tags were then inserted into *ricO* by overlap PCR using the primer pair 9436/10678. The full constructs were digested with EcoRI/SalI and cloned into equally digested pMAD. To insert sequences encoding fluorescent proteins into the genomic *scpAB* locus, DNA fragments with approximately 600 bp spanning 3′ ends (excluding stop codons) of the *scpA* and *scpB* genes from the JE2 genome were amplified using the primer pairs 7476/7477 and 7480/7483, respectively. The *mNG* and *mcherry* coding sequences from pBCBSS135 ^84^ and pMAD-rodAmCh ^83^ were amplified using the primer pairs 7494/7495 and 7456/7457, respectively. The downstream region of the *scpA* and *scpB* genes from the JE2 genome were amplified using the primer pairs 7472/7475 and 7478/7479, respectively. Corresponding DNA fragments were cloned into SmaI-digested pMAD via Gibson assembly (NEB). pMAD derivatives were electroporated into RN4220 and transduced to JE2 derivative strains. Integration excision by a double homologous recombination event was performed at 30 °C and selected genomic loci were amplified by PCR to identify clones that contained the inserted sequences.

Strains harbouring plasmid-borne *ricO* and *ricS1* were constructed using pEPSA5 ^59^, a replicative plasmid that allows gene expression under control of the xylose-inducible T5X promoter, or its derivatives pEPSA-P*tcaB* and pEPSA-P*amaP* containing endogenous *S. aureus* promoters. To construct pEPSA5-*ricS1*, a DNA fragment containing *ricS1* (5ʹ-GTTTGTCTGAGGTTAGAC-3ʹ) and its genomic flanking regions (39 bp and 40 bp in length) was generated by annealing the two oligonucleotides 10133/10134, followed by ligation of the 97-bp fragment into BamHI/EcoRI-digested pEPSA5. pEPSA5-*ricS1* and the empty vector pEPSA5 were electroporated into RN4220 and transduced to JE2, JE2Δ*ricO,* JE2_Δ*parB* and JE2_Δ*scpAB.* To construct pEPSA-P*tcaB* and pEPSA-P*amaP*, the transcriptional terminator sequence from pCN51 vector ^64^ and an approximately 200-bp upstream genomic region of the *tcaB* and *amaP* genes from the JE2 genome were amplified using the primer pairs 10191/10192, 10255/10256 and 10189/10190, respectively. DNA fragments containing the transcriptional terminator and promoter sequences were digested with SmaI/XhoI and XhoI/EcoRI, respectively. Fragments were joined by ligation, and the full constructs were cloned into EcoRV/EcoRI-digested pEPSA5. The *ricO* coding sequence was then amplified from the JE2 genome using the primer pair 10123/10124, digested with EcoRI/BamHI and cloned into equally digested pEPSA-P*tcaB* and pEPSA-P*amaP.* The resulting plasmids pEPSA-P*tcaB-ricO* and pEPSA-P*amaP-ricO* were electroporated into RN4220 and transduced to JE2 derivative strains. pEPSA-P*tcaB-ricO* was used to complement *ricO* deletion phenotypes and pEPSA-P*amaP-ricO* was used to assess *ricO* overexpression phenotypes.

Strains producing plasmid-borne RicO^SW^-GFP and GFP were constructed using pBCB50, a derivative of the pCN51 vector ^64^ that allows gene expression under control of the Atc-inducible *xyl-tetO* promoter. To construct pBCB50, the pIMAY-Z region encompassing *tetR* and P*xyl-tetO* was amplified using the primer pair 9626/9627, digested with PaeI/PstI and cloned into equally digested pCN51. Note that primer 9627 contains a *tetO* sequence for an improved TetR repression. The *gfp* and *ricO^SW^-gfp* coding sequences were amplified from pMAD-ezrAsgfp ^83^ and the JE2*ricO^SW^-gfp* genome, respectively, digested with PstI/EcoRI and cloned into equally digested pBCB50. The resulting plasmids pBCB50-*gfp* and pBCB50-*ricO^SW^-gfp* were electroporated into RN4220 and transduced to JE2.

To construct pBCBAIM012 and pBCBAIM009 for heterologous production of recombinant RicO in *E. coli*, the *ricO* coding sequence from the JE2 genome was amplified using the primer pairs 9906/10108 (codons 1-174) and 9906/9907 (codons 1-319), respectively. The resulting DNA fragments were digested with SacI/BamHI and cloned into equally digested pTB146. To introduce the *ricO*(R18A) mutation into the plasmid pBCBAIM012, an inverse PCR was performed using the primer pair 10163/10162, resulting in plasmid pBCBAIM015. The presence of the R18A mutation was verified by sequencing.

### Growth curves of *S. aureus* strains

To assess growth of JE2 derivative strains, overnight cultures in TSB (supplemented with chloramphenicol where necessary) at 37 °C were back-diluted to an OD_600_ of 0.01 in 20 ml TSB rich medium without antibiotics, in 100-ml Erlenmeyer flasks. Cells were grown with shaking at 200 rpm at 37 °C in triplicate. 1-ml samples were taken every 60 min to record the OD_600_ using a spectrophotometer (Biochrom Ultrospec 2100 Pro).

### Fluorescence microscopy

To quantify the anucleate cell fraction of *S. aureus* strains, overnight cultures of JE2, COL and NCTC8325-4 derivative strains were inoculated in TSB (supplemented with chloramphenicol for strains harbouring pEPSA5 or derivatives) in triplicates from independent single colonies and incubated with shaking at 37 °C. Cultures were diluted 1:200 into fresh TSB without antibiotics, followed by incubation with shaking at the same temperature. A total of 1 ml of mid-exponential growth phase cells (OD_600_ of 0.6-0.8) was mixed with Hoechst 33342 (1 μg/ml, Invitrogen) and incubated with shaking for 5 min at 37 °C. Cells were pelleted by centrifugation for 1 min at 9,300*g*, re-suspended in phosphate-buffered saline (PBS; 137 mM NaCl, 2.7 mM KCl, 10 mM Na_2_HPO_4_, 1.8 mM KH_2_PO_4_, pH 7.4) and spotted on a microscope slide covered with a thin layer of 1.5% TopVision agarose (Thermo Fisher) in PBS. Images were acquired with a Zeiss Axio Observer microscope equipped with a Plan-Apochromat 100×/1.4 oil Ph3 objective, a Retiga R1 CCD camera (QImaging), a white-light source HXP 120 V (Zeiss) and the software ZEN blue v2.0.0.0 (Zeiss). For image acquisition, the filter (Semrock) Brightline DAPI-1160A (Hoechst 33342) was used.

To visualize replication origins in live *S. aureus* cells and generate average localization heat maps, the strain JE2_FROS^Ori^ and its derivatives were grown overnight in TSB (supplemented with chloramphenicol for strains harbouring pEPSA5 or derivatives) and diluted 1:200 in fresh TSB without antibiotics, followed by incubation with shaking at 37 °C. Cells were grown in the absence of inducer, as leaky expression of *tetR-mNG* from the *cad* promoter was sufficient for imaging. After cells reached mid-exponential growth phase (OD_600_ of 0.6-0.8), they were pelleted by centrifugation for 1 min at 9,300*g*, re-suspended in PBS and spotted on a microscope slide covered with a thin layer of 1.5% TopVision agarose (Thermo Fisher) in PBS. Images were acquired with a Zeiss Axio Observer microscope using the filter (Semrock) Brightline GFP-3035B (mNG).

To localize RicO^SW^-GFP in cells with changes in the number of *ricS* sites and generate average localization heat maps, the strains JE2*ricO^SW^-gfp,* JE2Δ9*ricSricO^SW^-gfp*, JE2*ricO^SW^-gfp*pEPSA5 and JE2*ricO^SW^-gfp*pEPSA5-*ricS1* were grown overnight in TSB (supplemented with chloramphenicol for the latter two strains) and diluted 1:200 in fresh TSB without antibiotics, followed by incubation with shaking at 37 °C. Cells in mid-exponential growth phase (OD_600_ of 0.6-0.8) were pelleted by centrifugation for 1 min at 9,300*g*, re-suspended in PBS and spotted on a microscope slide covered with a thin layer of 1.5% TopVision agarose (Thermo Fisher) in PBS. Images were acquired with a Zeiss Axio Observer microscope using the filter (Semrock) Brightline GFP-3035B (mNG).

To colocalize RicO^SW^-GFP with the membrane and the nucleoid, overnight cultures of the strains JE2*ricO^SW^-gfp,* JE2*ricO*(R18A)*^SW^-gfp*, JE2*ricO*(ΔAH)*^SW^-gfp,* JE2pBCB50*gfp* and JE2pBCB50*ricO^SW^-gfp* were inoculated in TSB (supplemented with erythromycin for the latter two strains) from independent single colonies and incubated with shaking at 37 °C. Cultures were diluted 1:200 into fresh TSB without antibiotics (supplemented with Atc for the strains JE2pBCB50*gfp* and JE2pBCB50*ricO^SW^-gfp*), followed by incubation with shaking at the same temperature. A total of 1 ml of mid-exponential growth phase cells (OD_600_ of 0.6-0.8) was mixed with FM5-95 (1 µg/ml, Invitrogen) and Hoechst 33342 (1 µg/ml, Invitrogen), followed by incubation with shaking for 5 min at 37 °C. Cells were pelleted by centrifugation for 1 min at 9,300*g*, re-suspended in PBS and spotted on a pad of 1.5% TopVision agarose (Thermo Fisher) in PBS. Images were acquired with a Zeiss Axio Observer microscope using the filters (Semrock) Brightline DAPI-1160A (Hoechst 33342), Brightline GFP-3035B (GFP) and Brightline TXRED-4040B (FM5-95).

To colocalize RicO^SW^-mCherry with TetR-mNG (labelling four different chromosomal loci) or ScpB-mCherry with ScpA-mNG, overnight cultures in TSB of the strains JE2*oriC*^mNG^*ricO^SW^-mcherry*, JE2*left*^mNG^*ricO^SW^-mcherry*, JE2*right*^mNG^*ricO^SW^-mcherry*, JE2*terC*^mNG^*ricO^SW^-mcherry* and JE2*scpA-mNGscpB-mcherry* were inoculated in triplicates from independent single colonies and incubated with shaking at 37 °C. After cells reached mid-exponential growth phase (OD_600_ of 0.6-0.8), they were pelleted by centrifugation for 1 min at 9,300*g*, re-suspended in PBS and spotted on a microscope slide covered with a thin layer of 1.5% TopVision agarose (Thermo Fisher) in PBS. Images were acquired with a Zeiss Axio Observer microscope using the filters (Semrock) Brightline GFP-3035B (mNG) and Brightline TXRED-4040B (mCherry).

To determine the number of foci per cell and measure the distance between foci corresponding to RicO^SW^-SNAP and replication origins, the strains JE2_FROS^Ori^, JE2*oriC*^mNG^Δ*ricO* and JE2*oriC*^mNG^*ricO^SW^-snap* were grown overnight in TSB and diluted 1:200 in fresh TSB followed by incubation with shaking at 37 °C. A total of 0.6 ml of mid-exponential growth phase cells (OD_600_ of 0.6-0.8) was mixed with 1 µl of 1000x CellBrite Fix 640 (Biotium) for 20 min at 37 °C and with Hoechst 33342 (1 μg/ml, Invitrogen) for 5 min at 37 °C. JE2*oriC*^mNG^*ricO^SW^-snap* cells were additionally labelled with JF549-cpSTL (83 nM, red-fluorescent Janelia Fluor 549 cell-permeable SNAP-tag ligand) for 20 min at 37 °C. Cells were pelleted by centrifugation for 1 min at 9,300*g*, washed one time, re-suspended in TSB and spotted on a microscope slide covered with a thin layer of 1.5% TopVision agarose (Thermo Fisher) in TSB:PBS (1:3). Imaging was performed in a DeltaVision OMX SR microscope (GE Healthcare) equipped with an Olympus 60X PlanApo N/1.42 oil differential interference contrast objective and two PCO Edge 5.5 sCMOS cameras (one for Hoechst 33342 and mNG; one for JF549 and CellBrite Fix 640). The software AcquireSR v4.4 (GE Healthcare) was programmed to acquire *Z*-stacks of three images with a step size of 500 nm using a 405 nm laser (100 mW, at 30% maximal power), a 488 nm laser (100 mW, at 20% maximal power), a 568 nm laser (100 mW, at 50% maximal power) and a 640 nm laser (100 mW, at 30% maximal power), each with an exposure time of 100 ms. The software SoftWorRx v7.2.1 was used for fluorescence channel alignment, maximum intensity projection (MIP) of three images from each *Z*-stack (for mNG and JF549), and deconvolution of projected mNG and JF549 images.

To visualize segregating chromosomes in actively growing *S. aureus* cells, the strains JE2, JE2Δ*ricO* and JE2pBCB50*ricO^SW^-gfp* were grown overnight in TSB (supplemented with erythromycin for the latter strain) and diluted 1:200 in fresh TSB without antibiotics, followed by incubation with shaking at 37 °C. A total of 0.1 ml of mid-exponential growth phase cells (OD_600_ of 0.6-0.8) was mixed with 1 µl of 1000x CellBrite Fix 640 (Biotium) for 20 min at 37 °C and with Hoechst 33342 (5 μg/ml, Invitrogen) for 5 min at 37 °C. Cells were pelleted by centrifugation for 1 min at 9,300*g*, re-suspended in TSB and spotted on a microscope slide covered with a thin layer of 1.5% TopVision agarose (Thermo Fisher) in TSB:PBS (1:3) supplemented with 2 μg/ml Hoechst 33342. For the strain JE2pBCB50*ricO^SW^-gfp,* the agarose was additionally supplemented with 50 ng/ml Atc to induce production of RicO^SW^-GFP. Images were acquired in a DeltaVision OMX SR microscope equipped with an environmental control module set to 37 °C. Epifluorescence images (for Hoechst 33342 and GFP) and Structured Illumination Microscopy (SIM) images (for CellBrite Fix 640; three phase shifts and three grid rotations) were acquired every 15 min for 75 min using a 405 nm laser (100 mW, at 30% maximal power), a 488 nm laser (100 mW, at 20% maximal power) and a 640 nm laser (100 mW, at 30% maximal power), respectively, each with an exposure time of 25 ms. CellBrite Fix 640 images were reconstructed using SoftWoRx v7.2.1, time frames were aligned using NanoJ v2.1RC1 drift correction ^85^, and fluorescence channels were manually aligned using ImageJ/Fiji v1.53 ^86^.

To visualize the movement dynamics of replication origins during the cell cycle, the strains JE2_FROS^Ori^ and JE2*oriC*^mNG^Δ*ricO* were grown overnight in TSB and diluted 1:200 in fresh TSB followed by incubation with shaking at 37 °C. A total of 0.1 ml of mid-exponential growth phase cells (OD_600_ of 0.6-0.8) was mixed with 1 µl of 1000x CellBrite Fix 640 (Biotium) for 20 min at 37 °C. Cells were pelleted by centrifugation for 1 min at 9,300*g*, re-suspended in TSB and spotted on a microscope slide covered with a thin layer of 1.5% TopVision agarose (Thermo Fisher) in TSB:PBS (1:3). Images were acquired in a DeltaVision OMX SR microscope equipped with an environmental control module set to 37 °C. Three epifluorescence images in *Z*-stacks with a step size of 500 nm (for mNG) and SIM images (for CellBrite Fix 640; three phase shifts and three grid rotations) were acquired every 10 min for 50 min using a 488 nm laser (100 mW, at 10% maximal power) with an exposure time of 100 ms and a 640 nm laser (100 mW, at 30% maximal power) with an exposure time of 25 ms, respectively. SIM reconstruction of CellBrite Fix 640 images, MIP of mNG images from each *Z*-stack, and deconvolution of projected mNG images was performed for each time frame using SoftWoRx v7.2.1. Time frames and fluorescence channels were aligned as described above.

To colocalize RicO^SW^-SNAP with chromosomal origins in actively growing *S. aureus* cells, the strain JE2*oriC*^mNG^*ricO^SW^-snap* was grown overnight in TSB and diluted 1:200 in fresh TSB followed by incubation with shaking at 37 °C. A total of 0.6 ml of mid-exponential growth phase cells (OD_600_ of 0.6-0.8) was mixed with 1 µl of 1000x CellBrite Fix 640 (Biotium) and 2 µl of 25 µM JF549-cpSTL (83 nM final concentration) for 20 min at 37 °C. Cells were pelleted by centrifugation for 1 min at 9,300*g*, washed one time, re-suspended in TSB and spotted on a microscope slide covered with a thin layer of 1.5% TopVision agarose (Thermo Fisher) in TSB:PBS (1:3). Images were acquired in a DeltaVision OMX SR microscope equipped with an environmental control module set to 37 °C. Three epifluorescence images in *Z*-stacks with a step size of 500 nm (for mNG and JF549) and SIM images (for CellBrite Fix 640; three phase shifts and three grid rotations) were acquired every 8 min for 40 min using a 488 nm laser (100 mW, at 10% maximal power), a 568 nm laser (100 mW, at 20% maximal power) and a 640 nm laser (100 mW, at 30% maximal power), respectively, each with an exposure time of 100 ms. SIM reconstruction of CellBrite Fix 640 images, MIP of mNG and JF549 images from each *Z*-stack, and deconvolution of projected mNG and JF549 images was performed for each time frame using SoftWoRx v7.2.1. Time frames and fluorescence channels were aligned as described above.

### Microscopy image analysis

Generation of average localization heat maps and quantification of fluorescent foci was performed as previously described ^41^. To generate heat maps, cell segmentation was performed using eHooke software version 1.1 ^87^ for phase contrast images. To quantify fluorescent foci, cell segmentation was performed using an in-house fine-tuned StarDist model ^88^ for images with fluorescence signal from membrane labelling, including 1-pixel mask dilation. A PCA transform was applied to the coordinates of the pixels that constitute the cell outline to calculate the orientation of the major axis of each cell. Then, cell crops were aligned by their major axes as previously described ^84^. Spot detection in fluorescence channels was performed in TrackMate 7.11.1 ^89^ using the Laplacian of Gaussian filter with subpixel localization. The blob radius was set to 125 nm, the quality threshold was adjusted for each field of view (automatic adjustment for heat maps and manual adjustment for foci quantification), and the results were exported as xml files. In each cell crop, spots were represented as a circle of 1 pixel radius, an intensity of 1, and the same relative coordinates as the spot, in a rectangle with the same dimensions as the cell crop (model image), with background set to 0. All model images were then resized to a common width and height equal to the median of the width and height of all cell crops. Heat maps were generated by averaging all model images and coloured using the coolwarm colormap provided by matplotlib ^90^. The distance between red- and green-fluorescent foci was measured based on the spot coordinates determined for a given RicO^SW^-SNAP focus (red channel) relative to its nearest *oriC* focus (green channel) located in the same cell. Colocalization of two fluorescent proteins or of a fluorescent protein with membrane and DNA dyes was assessed by measuring the Pearson Correlation Coefficient (PCC) as previously described ^91^.

To perform automated quantification of cells with nucleoid defects, DNA-stained cells were segmented using the software eHooke as previously described ^87^. For each cell, the percentage of the cell area with an intensity in the DNA channel below a predetermined threshold was calculated. The threshold was calculated for each field of view. Briefly, the pixel values in the DNA channel were binned and the most common bin was calculated. Every intensity value that was higher than the largest value of the most common bin was assigned the value of the most common bin. The final threshold was calculated from the rescaled DNA image using the ISODATA clustering algorithm^92^. A cell was considered as having nucleoid defects when the DNA signal was below this threshold in at least 25% of the cell area.

### Protein in-gel fluorescence detection and Western blot analysis

To assess the integrity of RicO translationally fused to mCherry, SNAP-tag and GFP, as well as its fluorescent mutant derivatives R18A and ΔAH, the strains JE2, JE2*ricO^SW^-mcherry,* JE2*ricO^SW^-snap,* JE2*ricO^SW^-gfp,* JE2*ricO*(R18A)*^SW^-gfp* and JE2*ricO*(ΔAH)*^SW^-gfp* were grown to mid-exponential phase (OD_600_ of 0.6-0.8) in 50 ml TSB at 37 °C. Cells were cooled on ice and harvested by centrifugation for 10 min at 7,200*g* at 4 °C. JE2 and JE2*ricO^SW^-snap* cells were re-suspended in 0.3 ml fresh TSB and labelled with 83 nM of JF549-cpSTL for 20 min at 37 °C. Cells were cooled on ice, washed one time and re-suspended in 0.3 ml of PBS supplemented with complete mini protease Inhibitor Cocktail (Roche). Cell suspensions were transferred to lysis tubes containing glass beads and subjected to mechanical disruption in a FastPrep-24 bead beating grinder and lysis system (MP Biomedicals) programmed to two 45-s cycles. Glass beads and cell debris were removed in two steps of centrifugation each for 1 min at 3,400*g* at 4°C. A total of 20 μl of non-boiled protein sample were loaded on 12% Mini-Protean TGX pre-cast gels (Bio-Rad) and the proteins were separated by sodium dodecyl-sulfate polyacrylamide gel electrophoresis (SDS-PAGE). Gels were imaged in a FujiFilm FLA-5100 imaging system. RicO^SW^-GFP fluorescence was detected using 473 nm laser/Cy3 filter, RicO^SW^-mCherry and JF549-labelled RicO^SW^-SNAP fluorescence was detected using 532 nm laser/LPB filter, and pre-stained molecular weight marker (PageRuler, Thermo Scientific) was detected using 635 nm laser/LPFR filter settings. Gels were post-stained with InstantBlue Coomassie stain (Abcam).

To detect RicO wild-type protein in various genetic backgrounds, as well as its mutant derivatives R18A and ΔAH, JE2 derivative strains were grown to mid-exponential phase (OD_600_ of 0.6-0.8) in 50 ml TSB at 37 °C. Cells were cooled on ice, harvested by centrifugation and re-suspended in 0.3 ml PBS supplemented with complete mini protease Inhibitor Cocktail (Roche). Whole-cell protein extracts were obtained and processed as described above, except that additional protein denaturation was performed at 100 °C for 10 min before gel loading. A total of 20 μl of protein sample were loaded on 8% polyacrylamide gels and the proteins were separated by SDS-PAGE. Separated proteins were then transferred to a nitrocellulose membrane using a Trans-Blot Turbo RTA Mini 0.2 μm Nitrocellulose Transfer Kit and Trans-Blot Turbo system (Bio-Rad). The membrane was cut to separate the regions above and below approximately 100 kDa. The top part of the membrane was stained with Sypro-Ruby stain (Invitrogen) following the manufacturer’s instructions to label high-molecular weight proteins as gel loading control. The bottom part of the membrane containing RicO was blocked with 5% milk, followed by consecutive incubations with an anti-RicO antibody (1:1,000 dilution) for 16 h at 4 °C and with a secondary fluorescent antibody (StarBright anti-rabbit diluted 1:50,000; BioRad) for 1 h at room temperature. StarBright and Sypro-Ruby fluorescence detection was performed in an iBright Imaging System (Invitrogen). Uncropped images of gels and blotted membranes can be found in Supplementary Figure 14.

### Purification of RicO and antibody production

A total of three recombinant RicO variants were produced as 6xHis-SUMO fusions using pTB146 as backbone (see above for construction details). The expression was performed using *E. coli* Rosetta(DE3) (Novagen). Cultures of Rosetta(DE3) harbouring the pTB146 derivatives pBCBAIM009 (encoding full-length RicO), pBCBAIM012 (encoding C-terminally truncated RicO) or pBCBAIM015 (encoding C-terminally truncated RicO with the R18A mutation) were grown in LB medium supplemented with ampicillin at 37 °C with shaking until they reached an OD_600_ of 0.6. Cultures were cooled to 18 °C and isopropyl β-d-1-thiogalactopyranoside (IPTG, NZYtech) was added to a final concentration of 1 mM, followed by overnight incubation with shaking. Cells were pelleted by centrifugation at 5,000*g* at 4 °C, washed with buffer BZ3 (50 mM Tris/HCl pH 8.0, 300 mM NaCl, 30 mM imidazole, 10% glycerol), and centrifuged again at 5,000*g* at 4°C. The cell pellets were stored at -80 °C. When required, cell pellets of the desired strain were thawed on ice and re-suspended in buffer BZ3^HS^ (50 mM Tris/HCl pH 8.0, 1 M NaCl, 30 mM imidazole, 10% glycerol) supplemented with 5 mM MgCl_2_, 1 mM phenylmethanesulfonyl fluoride (Sigma-Aldrich), 1 µl of Benzonase (NZYtech) and 0.5 mg/ml of lysozyme (Sigma-Aldrich). The cell suspension was sonicated using a Branson sonicator at 35% power in eight 10-s pulses, with cooling on ice in between. The cell lysate was centrifuged at 25,000*g* for 30 min at 4 °C and the supernatant was passed through a filter with a 0.22-µm pore size. The filtered lysate was then loaded onto a 5 ml HisTrap HP column (Cytiva) pre-equilibrated with BZ3^HS^ buffer, using an AKTA Start chromatographic system (Cytiva). After loading, the column was washed with ten column volumes of buffer BZ3^HS^ before the bound protein was eluted with a linear gradient of BZ3^HS^-BZ4^HS^ (50 mM Tris/HCl pH 8.0, 1 M NaCl, 250 mM imidazole, 10% glycerol). Fractions enriched in recombinant RicO were pooled and the 6xHis-Ulp1 protease (purified from *E. coli* BL21(DE3) carrying plasmid pTB145) was added in a 1:1000 ratio to the protein content in the solution. The mixture was dialyzed in 2 L of CB buffer (50 mM Tris/HCl pH 8.0, 300 mM NaCl, 10% glycerol) for at least 16 h before loading it again onto the 5 ml HisTrap HP column pre-equilibrated with BZ3^HS^ buffer. Fractions from the flow-through enriched in free RicO protein (cleaved from the 6xHis-SUMO) were pooled and dialyzed against storage buffer (50 mM HEPES/NaOH pH 7.4, 300 mM NaCl, 10% glycerol) for at least 16 h. The protein solution was then concentrated using Amicon centrifugal filters (Millipore), aliquoted, snap-frozen in liquid nitrogen and stored at -80 °C until further use.

To produce specific antibodies against RicO, purified full-length RicO was loaded onto a polyacrylamide gel and stained with Coomassie Instant Blue (Abcam). Gel bands containing the RicO protein were sent to Eurogentech where the immunization of rabbits was carried out. Serum from the final bleed was tested for specificity and used directly where required.

### Electrophoretic mobility shift assay (EMSA)

The DNA fragments *ricS1* and *ricS1\** were generated by hybridization of the complementary oligonucleotides 10207/10208 and 10209/10210, respectively. Oligonucleotides were mixed in an equimolar concentration, creating a 50 mM stock that was subjected to three cycles of 10 min at 98 °C, followed by slow cooling to room temperature. For fluorescence detection of the DNA fragments, one of the two oligonucleotides was synthetized with a 6-FAM fluorophore attached to its 5’-end. EMSA reactions were performed in a buffer of 10 mM HEPES/NaOH pH 7.2, 150 mM NaCl, 10 mM MgCl_2_ and incubated at room temperature for 12 min. Unless stated otherwise, all reactions were performed with 0.2 µM of DNA. Purified RicO variants were used at various concentrations ranging from 0 to 40 µM. Glycerol was added to each sample at a final concentration of 6% and the samples were analyzed in a 6% polyacrylamide gel in 0.5x TBE buffer (45 mM Tris, 45 mM boric acid, 1 mM EDTA). Gels were run for 65 min at 70 V at room temperature and then imaged using an iBright Imaging System (Invitrogen).

### Whole-genome sequencing (WGS) and Marker-frequency analysis (MFA)

Genomic DNA extraction, sequencing and MFA were performed as previously described ^41^. Briefly, cells of the strains JE2 and JE2Δ*ricO* were grown in TSB at 37 °C with shaking until the early-exponential phase (OD_600_ 0.3-0.4) and centrifuged for 10 min at 7,200*g* at 4 °C. Genomic DNA was isolated using the DNeasy Blood & Tissue Kit (Qiagen), fragmented using a Qsonica Q800R2 water bath sonicator, prepared using the NEBNext UltraII kit (E7645), and sequenced at the Indiana University Center for Genomics and Bioinformatics using NextSeq2000. The reads were mapped to the genome of *S. aureus* JE2 (NCBI Reference Sequence GCF_002085525.1) using CLC Genomics Workbench (CLC Bio, QIAGEN). The mapped reads were normalized by the total number of reads. Plotting and analysis were performed using R scripts.

### Chromatin immunoprecipitation sequencing (ChIP-seq) and ChIP-seq analysis

ChIP-seq was performed as previously described ^93^. Briefly, cells of the *spa* deletion strains JE2Δ*spa*, JE2Δ*ricO*Δ*spa,* JE2Δ9*ricS*Δ*spa*, JE2*ricO*(R18A)Δ*spa* and JE2*ricO*(ΔAH)Δ*spa* were grown in TSB at 37 °C with shaking until they reached early-exponential phase (OD_600_ 0.3-0.4). Cells were crosslinked with formaldehyde (3%, Sigma-Aldrich) at room temperature for 30 min, followed by quenching with 125 mM glycine (Sigma-Aldrich). Cells were centrifuged for 10 min at 7,200*g* at 4 °C and washed three times in ice-cold PBS. The cell pellets were re-suspended in 500 μl ChIP buffer A (12.5 mM Tris pH 8.0, 12.5 mM EDTA pH 8, 62.5 mM NaCl, 25% sucrose), snap-frozen in liquid nitrogen and stored at -80°C until further use. Cells were lysed using lysozyme (4 mg/ml, VWR, 97062-136) and lysostaphin (200 µg/ml, Sigma-Aldrich, L9043). Crosslinked chromatin was sheared to an average size of 170 bp by sonication using a Qsonica Q800R2 waterbath sonicator. The lysate was precleared using Protein A magnetic beads (GE Healthcare/Cytiva 28951378) and was then incubated with anti-RicO antibodies overnight at 4 °C. Next day, the lysate was incubated with Protein A magnetic beads for one hour at 4 °C. After washes and elution, the immunoprecipitate was incubated at 65 °C overnight to reverse the crosslinks. The DNA was further treated with RNase A, Proteinase K, extracted with phenol/chloroform/isoamylalcohol (25:24:1) (PCI), re-suspended in 100 µl of Qiagen EB buffer and used for library preparation with the NEBNext UltraII kit (E7645). The library was sequenced at the Indiana University Center for Genomics and Bioinformatics using NextSeq500 or NextSeq2000. The sequencing reads were mapped to the genome of *S. aureus* JE2 (NCBI Reference Sequence GCF_002085525.1) using CLC Genomics Workbench (CLC Bio, QIAGEN). Sequencing reads from each ChIP and input sample were normalized by the total number of reads. The ChIP enrichment (ChIP/Input) was plotted and analyzed using R scripts.

### Hi-C and Hi-C analysis

Chromosome conformation capture (Hi-C) was performed as previously described ^41,94^. Briefly, cells of the required strains were grown in TSB at 37 °C with shaking until they reached early-exponential phase (OD_600_ 0.3-0.4). Cells were crosslinked with formaldehyde (7%, Sigma-Aldrich) at room temperature for 30 min, followed by quenching with glycine (125 mM, Sigma-Aldrich). Cells were centrifuged for 10 min at 7,200*g* at 4°C and washed three times in ice-cold PBS. The cell pellets were snap-frozen in liquid nitrogen and stored at -80°C until further use. Cells were lysed using Ready-Lyse Lysozyme (Epicentre, R1802M) and lysostaphin (200 µg/ml, Sigma-Aldrich, L9043), followed by treatment with SDS (1% final concentration). Solubilized chromatin was digested with DpnII and the digested ends were filled in with Klenow and Biotin-14-dATP, dGTP, dCTP, dTTP. The products were ligated with T4 DNA ligase, followed by reversal of cross-links. The DNA was then extracted twice with PCI, precipitated with ethanol, and re-suspended in 20 µl of 0.1x TE buffer (10 mM Tris-HCl, 1 mM EDTA). Biotin from non-ligated ends was removed using T4 polymerase, followed by extraction with PCI. The DNA was then sheared by sonication using a Qsonica Q800R2 water bath sonicator and used for library preparation with the NEBNext UltraII kit (E7645). Biotinylated DNA fragments were purified using streptavidin beads and DNA-bound beads were used for PCR. PCR products were purified using Ampure beads (Beckman, A63881) and sequenced at the Indiana University Center for Genomics and Bioinformatics using NextSeq500. Paired-end sequencing reads were mapped to the genome of *S. aureus* JE2 (NCBI Reference Sequence GCF_002085525.1) using the same pipeline as previously described ^93^. The genome was divided into 5-kb or 1-kb bins. Subsequent analysis and visualization were done using R scripts. Hi-C scores, which quantify the interaction between loci and correct for biases in the abundance of the different bins in each experiment, were calculated as previously described ^93^.

### Bioinformatic analysis of RicO conservation

The NCBI BLAST tool ^95^ and the sequences of RicO, RacA and RocS (Uniprot accession codes Q2G0Z2_STAA8, RACA_BACSU and ROCS_STRR6, respectively) were used to search for homologs in closely related species. 15 proteins were identified that showed both sequence and structure similarities as predicted by AlphaFold 3 ^96^ (Uniprot accession codes: A0A662Z4U2_9STAP, Y2353_STAES, A0A078M2D1_9STAP, A0A0C2DMP9_9STAP, A0A240AF17_9STAP, Y1595_STRP6, A0A7X1ZBY2_9LACT, A0A841C711_9LACT, A0A1E8GJF8_9LACT, RACA_GEOKA, S0JDT9_9ENTE, RACA_BACAC, RACA_BACCR, D5DZ83_PRIM1). All 18 protein sequences were aligned using T-COFFEE Expresso mode ^97^ and the alignment was used to generate a Hidden Markov Model (HMM) of a potential protein family using the suite HMMer v.3.4 ^98^. This HMM profile was then used to search with HMMSEARCH potential homologs in a local databank of Firmicutes containing one proteome per genus as described in Luhur *et al.* ^99^. We selected hits with an e-value below 10^-4^, yielding 137 sequences of potential homologs. The sequences were aligned using MAFFT v.7 with the L-INS-i algorithm^100^ and the subsequent alignment was trimmed using trimAL ^101^ with the gappyout mode. The trimmed alignment (185 amino acid positions) was used to infer a maximum likelihood tree using IQtree 3.0.1 ^102^, with the substitution model Q.PFAM+F+G4 chosen by model finder ^103^. Branch supports were assessed using ultra-fast bootstrapping (UFBoot) ^104^. The Pfam domain database (v. 38.0) and HMMSCAN from the HMMer suite were used to identify domains. The CoCoNat server ^67^ was used to identify coiled-coil regions with a minimum length of ten amino acids. Tree representation and annotation were done using iTOL v.7.2 ^105^.

## Data availability

The Hi-C, ChIP-seq and WGS data generated in this study have been deposited in the NCBI Gene Expression Omnibus database under accession code GSE318607 [https://www.ncbi.nlm.nih.gov/geo/query/acc.cgi?acc=GSE318607] Source data are provided with this paper.

## Code availability

The codes used to plot the WGS, ChIP-seq and Hi-C results were deposited to github [https://github.com/xindanwanglab/Schaper-2026-RicO]. The codes used to analyse fluorescence microscopy images and create average localization heatmaps were deposited to github [https://github.com/BacterialCellBiologyLab/eHooke and https://github.com/BacterialCellBiologyLab/AverageCellLoc].

## Acknowledgements

We thank members of the Pinho lab, P. Pereira (ITQB-NOVA) and S. Filipe (FCT-NOVA) for stimulating discussions and support; H. Veiga (ITQB-NOVA) for sharing strain JE2Δ*spa*; T. Bernhardt (Harvard Medical School, Boston) for donating the plasmids pTB145 and pTB146; L. Lavis (Janelia Research Campus, Ashburn) for the generous gift of JF549-cpSTL; and C. Tesseur (UCLouvain, Brussels) for collecting preliminary data. We thank Indiana University Center for Genomics and Bioinformatics for high throughput sequencing. This study was funded by the European Research Council (ERC) through grant 101096393 (to M.G.P.); by the Fundação para a Ciência e a Tecnologia (FCT) through MOSTMICRO-ITQB R and D Unit (UID/04612/2025, UID/PRR/4612/2025 to ITQB-NOVA), LS4FUTURE Associated Laboratory (LA/P/0087/2020 to ITQB-NOVA), grant 2022.01678.PTDC (to S.S.) and contract 2022.03033.CEECIND (to S.S.). Research in the Wang laboratory was supported by National Institutes of Health R01GM141242, R01GM143182, and R01AI172822 (to X.W.). This research is a contribution of the GEMS Biology Integration Institute, funded by the National Science Foundation DBI Biology Integration Institutes Program, Award #2022049 (to X.W.). Figure 6e was created with BioRender.

## Author contributions

S.S., A.I.M., X.W. and M.G.P. designed the research; S.S., A.I.M. and Q.L. performed the experiments; S.S., A.I.M. and M.S. constructed strains; A.D.B. and B.M.S. developed software; A.I.M. performed the phylogenetic analysis with input from N.T.; all authors analysed data; S.S. and M.G.P. wrote the original draft; all authors revised the manuscript.

## Competing interests

The authors declare no competing interests.

## Notes

### Competing Interest Statement

The authors have declared no competing interest.

