## Supplementary Information for "Nucleoid-binding protein RicO anchors replication origins to the membrane to ensure correct chromosome segregation in *Staphylococcus aureus*"

**Table of contents**

- Supplementary Tables 1-3
- Supplementary Figures 1-14
- Supplementary References

### Table S1. Strains used in this study.

| Name | Description | Reference |
|---|---|---|
| ***Escherichia coli*** | | |
| DC10B | Δ*dcm* in DH10B background; Dam methylation only; for cloning | [1] |
| Rosetta(DE3) | $F^-$ *ompT hsdS*$_B$ ($r_B^-$ $m_B^-$) *gal dcm* (DE3) pRARE2 ($Cm^r$); for protein production | Novagen |
| BL21(DE3) | $F^-$ *ompT hsdS*$_B$ ($r_B^-$ $m_B^-$) *gal dcm* (DE3); for protein production | Novagen |
| ***Staphylococcus aureus*** | | |
| RN4220 | Restriction-negative derivative of NCTC8325-4 | [2] |
| JE2 | CA-MRSA strain | [3] |
| COL | HA-MRSA strain | [4] |
| NCTC8325-4 | MSSA strain | R. Novick |
| JE2Δ*spa* | JE2 with markerless deletion of *spa* | H. Veiga |
| JE2_Δ*scpAB* | JE2 with markerless deletion of *scpAB* | [5] |
| JE2_Δ*scpAB*+ScpAB | JE2_Δ*scpAB* harbouring pBCBSS327; $Cm^r$ | [5] |
| JE2_Δ*parB* | JE2 with markerless deletion of *parB* | [5] |
| JE2_FROS$^{Ori}$ | JE2 with *spa* replaced by P*cad-tetR-mNG* and *tetO48* inserted between *SAUS300_2631* and *SAUS300_2632* to label the origin of replication | [5] |
| JE2_FROS$^{Left}$ | JE2 with *spa* replaced by P*cad-tetR-mNG* and *tetO48* inserted between *SAUS300_1984* and *SAUS300_1985* to label the left chromosome arm | [5] |
| JE2_FROS$^{Right}$ | JE2 with *spa* replaced by P*cad-tetR-mNG* and *tetO48* inserted between *SAUS300_0660* and *SAUS300_0661* to label the right chromosome arm | [5] |
| JE2_FROS$^{Ter}$ | JE2 with *spa* replaced by P*cad-tetR-mNG* and *tetO48* inserted between *SAUS300_1326* and *SAUS300_1327* to label the terminus of replication | [5] |
| JE2_FROS$^{Ori}$_Δ*scpAB* | JE2_FROS$^{Ori}$ with markerless deletion of *scpAB* | [5] |
| JE2Δ*ricO* | JE2 with markerless deletion of *ricO* | This study |
| COLΔ*ricO* | COL with markerless deletion of *ricO* | This study |
| NCTC8325-4Δ*ricO* | NCTC8325-4 with markerless deletion of *ricO* | This study |
| JE2Δ*ricO*p*ricO* | JE2Δ*ricO* harbouring pEPSA-P*tcaB-ricO*; $Cm^r$ | This study |
| JE2Δ*ricO*Δ*scpAB* | JE2Δ*ricO* with markerless deletion of *scpAB* | This study |
| JE2Δ*ricO*Δ*scpAB*p*ricO* | JE2Δ*ricO*Δ*scpAB* harbouring pEPSA-P*tcaB-ricO*; $Cm^r$ | This study |
| JE2Δ*ricO*Δ*scpAB*p*scpAB* | JE2Δ*ricO*Δ*scpAB* harbouring pBCBSS327; $Cm^r$ | This study |
| JE2Δ*ricO*Δ*parB* | JE2Δ*parB* with markerless deletion of *ricO* | This study |
| JE2Δ*ricO*Δ*scpAB*Δ*parB* | JE2Δ*ricO*Δ*parB* with markerless deletion of *scpAB* | This study |
| JE2*oriC*$^{mNG}$Δ*ricO* | JE2_FROS$^{Ori}$ with markerless deletion of *ricO* | This study |
| JE2*oriC*$^{mNG}$Δ*ricO*p*ricO* | JE2*oriC*$^{mNG}$Δ*ricO* harbouring pEPSA-P*tcaB-ricO*; $Cm^r$ | This study |
| JE2*oriC*$^{mNG}$*ricO*$^{SW}$*-snap* | JE2_FROS$^{Ori}$ with a *ricO* translational sandwich fusion to *snap* | This study |
| JE2*oriC*$^{mNG}$*ricO*$^{SW}$*-mcherry* | JE2_FROS$^{Ori}$ with a *ricO* translational sandwich fusion to *mcherry* | This study |
| JE2*left*$^{mNG}$*ricO*$^{SW}$*-mcherry* | JE2_FROS$^{Left}$ with a *ricO* translational sandwich fusion to *mcherry* | This study |
| JE2*right*$^{mNG}$*ricO*$^{SW}$*-mcherry* | JE2_FROS$^{Right}$ with a *ricO* translational sandwich fusion to *mcherry* | This study |
| JE2*terC*$^{mNG}$*ricO*$^{SW}$*-mcherry* | JE2_FROS$^{Ter}$ with a *ricO* translational sandwich fusion to *mcherry* | This study |
| JE2*scpA-mNGscpB-mcherry* | JE2 with a *scpA* translational C-terminal fusion to *mneongreen* and a *scpB* translational C-terminal fusion to *mcherry* | This study |
| JE2vector | JE2 harbouring pEPSA-P*amaP*; $Cm^r$ | This study |
| JE2*ricO*OE | JE2 harbouring pEPSA-P*amaP-ricO*; $Cm^r$ | This study |
| JE2pBCB50*gfp* | JE2 harbouring pBCB50-*gfp*; $Em^r$ | This study |
| JE2pBCB50*ricO*$^{SW}$*-gfp* | JE2 harbouring pBCB50-*ricO*$^{SW}$*-gfp*; $Em^r$ | This study |
| JE2Δ*spa*Δ*ricO* | JE2Δ*ricO* with markerless deletion of *spa* | This study |
| JE2Δ1*ricS* | JE2 with markerless deletion of *ricS9* | This study |
| JE2Δ2*ricS* | JE2Δ1*ricS* with markerless deletion of *ricS1* | This study |
| JE2Δ3*ricS* | JE2Δ2*ricS* with markerless deletion of *ricS7* | This study |
| JE2Δ4*ricS* | JE2Δ3*ricS* with markerless deletion of *ricS4* | This study |
| JE2Δ5*ricS* | JE2Δ4*ricS* with markerless deletion of *ricS2* | This study |
| JE2Δ6*ricS* | JE2Δ5*ricS* with markerless deletion of *ricS6* | This study |
| JE2Δ7*ricS* | JE2Δ6*ricS* with markerless deletion of *ricS8* | This study |
| JE2Δ8*ricS* | JE2Δ7*ricS* with markerless deletion of *ricS3* | This study |
| JE2Δ9*ricS* | JE2Δ8*ricS* with markerless deletion of *ricS5* | This study |
| JE2Δ*ricS5* | JE2 with markerless deletion of *ricS5* | This study |
| JE2Δ9*ricS*Δ*spa* | JE2Δ9*ricS* with markerless deletion of *spa* | This study |
| JE2Δ9*ricSricO*$^{SW}$*-gfp* | JE2Δ9*ricS* with a *ricO* translational sandwich fusion to *gfp* | This study |
| JE2*ricO*(R18A) | JE2 with native *ricO* replaced by a *ricO*(R18A) variant | This study |
| JE2*ricO*(R18A)Δ*spa* | JE2*ricO*(R18A) with markerless deletion of *spa* | This study |
| JE2*ricO*(ΔAH) | JE2 with native *ricO* replaced by a *ricO*(ΔAH) variant | This study |
| JE2*ricO*(ΔAH)Δ*spa* | JE2*ricO*(ΔAH) with markerless deletion of *spa* | This study |
| JE2Δ9*ricS*Δ*ricO* | JE2Δ9*ricS* with markerless deletion of *ricO* | This study |
| JE2Δ9*ricS*Δ*ricO*p*ricO* | JE2Δ9*ricS*Δ*ricO* harbouring pEPSA-P*tcaB-ricO*; $Cm^r$ | This study |
| JE2Δ9*ricS*Δ*scpAB* | JE2Δ9*ricS* with markerless deletion of *scpAB* | This study |
| JE2Δ9*ricS*Δ*scpAB*p*scpAB* | JE2Δ9*ricS*Δ*scpAB* harbouring pBCBSS327; $Cm^r$ | This study |
| JE2Δ9*ricS*Δ*parB* | JE2Δ9*ricS* with markerless deletion of *parB* | This study |
| JE2*oriC*$^{mNG}$Δ4*ricS* | JE2Δ4*ricS* with *tetO48* inserted between *SAUS300_2631-2* (358°) and *spa* replaced by P*cad-tetR-mNG* | This study |
| JE2*oriC*$^{mNG}$Δ5*ricS* | JE2*oriC*$^{mNG}$Δ4*ricS* with markerless deletion of *ricS2* | This study |
| JE2*oriC*$^{mNG}$Δ6*ricS* | JE2*oriC*$^{mNG}$Δ5*ricS* with markerless deletion of *ricS6* | This study |
| JE2*oriC*$^{mNG}$Δ7*ricS* | JE2*oriC*$^{mNG}$Δ6*ricS* with markerless deletion of *ricS8* | This study |

| | | |
|---|---|---|
| JE2*oriC*$^{mNG}$Δ8*ricS* | JE2*oriC*$^{mNG}$Δ7*ricS* with markerless deletion of *ricS3* | This study |
| JE2*oriC*$^{mNG}$Δ9*ricS* | JE2*oriC*$^{mNG}$Δ8*ricS* with markerless deletion of *ricS5* | This study |
| JE2*oriC*$^{mNG}$*ricO*(ΔAH) | JE2_FROS$^{Ori}$ with native *ricO* replaced by a *ricO*(ΔAH) variant | This study |
| JE2*oriC*$^{mNG}$*ricO*(R18A) | JE2*ricO*(R18A) with *tetO48* inserted between *SAUS300_2631-2* (358°) and *spa* replaced by P*cad-tetR-mNG* | This study |
| JE2*ricO*$^{SW}$-*mcherry* | JE2 with a *ricO* translational sandwich fusion to *mcherry* | This study |
| JE2*ricO*$^{SW}$-*snap* | JE2 with a *ricO* translational sandwich fusion to *snap* | This study |
| JE2*ricO*$^{SW}$-*gfp* | JE2 with a *ricO* translational sandwich fusion to *gfp* | This study |
| JE2*ricO*(R18A)$^{SW}$-*gfp* | JE2*ricO*(R18A) with a *ricO* translational sandwich fusion to *gfp* | This study |
| JE2*ricO*(ΔAH)$^{SW}$-*gfp* | JE2*ricO*$^{SW}$-*gfp* with a C-terminal truncation of 14 aa | This study |
| JE2pEPSA5 | JE2 harbouring pEPSA5; Cm$^{r}$ | This study |
| JE2pEPSA5-*ricS1* | JE2 harbouring pEPSA5-*ricS1*; Cm$^{r}$ | This study |
| JE2Δ*parB*pEPSA5 | JE2_Δ*parB* harbouring pEPSA5; Cm$^{r}$ | This study |
| JE2Δ*parB*pEPSA5-*ricS1* | JE2_Δ*parB* harbouring pEPSA5-*ricS1*; Cm$^{r}$ | This study |
| JE2Δ*ricO*pEPSA5 | JE2Δ*ricO* harbouring pEPSA5; Cm$^{r}$ | This study |
| JE2Δ*ricO*pEPSA5-*ricS1* | JE2Δ*ricO* harbouring pEPSA5-*ricS1*; Cm$^{r}$ | This study |
| JE2Δ*scpAB*pEPSA5 | JE2_Δ*scpAB* harbouring pEPSA5; Cm$^{r}$ | This study |
| JE2Δ*scpAB*pEPSA5-*ricS1* | JE2_Δ*scpAB* harbouring pEPSA5-*ricS1*; Cm$^{r}$ | This study |
| JE2*oriC*$^{mNG}$pEPSA5 | JE2*oriC*$^{mNG}$ harbouring pEPSA5; Cm$^{r}$ | This study |
| JE2*oriC*$^{mNG}$pEPSA5-*ricS1* | JE2*oriC*$^{mNG}$ harbouring pEPSA5-*ricS1*; Cm$^{r}$ | This study |
| JE2*ricO*$^{SW}$-*gfp*pEPSA5 | JE2*ricO*$^{SW}$-*gfp* harbouring pEPSA5; Cm$^{r}$ | This study |
| JE2*ricO*$^{SW}$-*gfp*pEPSA5-*ricS1* | JE2*ricO*$^{SW}$-*gfp* harbouring pEPSA5-*ricS1*; Cm$^{r}$ | This study |

### Table S2. Plasmids used in this study.

| Name | Description | Reference |
|---|---|---|
| pSNAP-tag (T7)-2 | *E. coli* expression vector encoding the SNAP-tag protein; Amp$^r$ | NEB |
| pIMAY-Z | *E. coli-S. aureus* shuttle vector with a thermosensitive origin of replication for Gram-positive bacteria; Cm$^r$, *lacZ* | [6] |
| pCN51 | *E.coli-S. aureus* shuttle replicative vector containing the cadmium-inducible *cad* promoter; Amp$^r$, Ery$^r$ | [7] |
| pEPSA5 | *E.coli-S. aureus* shuttle replicative vector containing a xylose-inducible T5X promoter; Amp$^r$, Cm$^r$ | [8] |
| pMAD | *E. coli-S. aureus* shuttle vector with a thermosensitive origin of replication for Gram-positive bacteria; Amp$^r$, Ery$^r$, *lacZ* | [9] |
| pMAD-ezrAsgfp | pMAD derivative containing a *ezrA-sgfp* fusion and the downstream region of *ezrA* | [10] |
| pMAD-rodAmCh | pMAD derivative containing a *rodA-mcherry* fusion and the downstream region of *rodA* | [10] |
| pBCBSS135 | pMAD derivative containing *3xflag-mNG* under P*spac* control | [11] |
| pMAD-Δ*spa* | pMAD containing *spa* upstream and downstream regions | [11] |
| pBCBSS289 | pMAD containing *tetO48* between the *SAUS300_2631-2* genes | [5] |
| pBCBAIM018 | pMAD derivative containing *tetR-mNG* under P*cad* control | [5] |
| pBCBMS086 | pMAD containing *scpA* upstream and *scpB* downstream regions | [5] |
| pBCBMS073 | pMAD containing *parB* upstream and downstream regions | [5] |
| pBCBSS327 | pEPSA5 containing the *scpA* and *scpB* coding sequences | [5] |
| pTB145 | Vector for production of the 6xHis-Ulp1 protease; Amp$^r$ | T. Bernhardt |
| pTB146 | Vector for production of N-terminal fusions to 6xHis-SUMO; Amp$^r$ | T. Bernhardt |
| pBCBAIM009 | pTB146 containing the *ricO* coding sequence from 1 to 957 bp | This study |
| pBCBAIM012 | pTB146 containing the *ricO* coding sequence from 1 to 522 bp | This study |
| pBCBAIM015 | pBCBAIM012 containing DNA-binding mutation R18A | This study |
| pBCB50 | pCN51 derivative containing *tetR* and tetracycline-inducible P*xyl-tetO* | This study |
| pBCB50-*gfp* | pBCB50 containing the *gfp* coding sequence | This study |
| pBCB50-*ricO*$^{SW}$-*gfp* | pBCB50 containing a *ricO* sandwich fusion to *gfp* | This study |
| pIMAY-Z-*ricO*(R18A) | pIMAY-Z containing *ricO*(R18A) mutation | This study |
| pEPSA5-*ricS1* | pEPSA5 containing the *ricS1* sequence with flanking regions | This study |
| pEPSA-P*tcaB* | pEPSA5 derivative containing the *tcaB* promoter region | This study |
| pEPSA-P*tcaB-ricO* | pEPSA-P*tcaB* containing the *ricO* coding sequence | This study |
| pEPSA-P*amaP* | pEPSA5 derivative containing the *amaP* promoter region | This study |
| pEPSA-P*amaP-ricO* | pEPSA-P*amaP* containing the *ricO* coding sequence | This study |
| pMAD-Δ*ricO* | pMAD containing *ricO* upstream and downstream regions | This study |
| pMAD-Δ*ricS1* | pMAD containing *ricS1* upstream and downstream regions | This study |
| pMAD-Δ*ricS2* | pMAD containing *ricS2* upstream and downstream regions | This study |
| pMAD-Δ*ricS3* | pMAD containing *ricS3* upstream and downstream regions | This study |
| pMAD-Δ*ricS4* | pMAD containing *ricS4* upstream and downstream regions | This study |
| pIMAY-Z-Δ*ricS5* | pMAD containing *ricS5* upstream and downstream regions | This study |
| pMAD-Δ*ricS6* | pMAD containing *ricS6* upstream and downstream regions | This study |
| pMAD-Δ*ricS7* | pMAD containing *ricS7* upstream and downstream regions | This study |
| pMAD-Δ*ricS8* | pMAD containing *ricS8* upstream and downstream regions | This study |
| pMAD-Δ*ricS9* | pMAD containing *ricS9* upstream and downstream regions | This study |
| pMAD-Δ*ricS5*Δ*ricO* | pMAD containing *ricO* upstream and downstream regions including Δ*ricS5* | This study |
| pMAD-*ricO*(ΔAH) | pMAD containing *ricO*(AH) upstream and downstream regions | This study |
| pMAD-*ricO*$^{sw}$-*mcherry* | pMAD containing a *ricO* sandwich fusion to *mcherry* | This study |
| pMAD-*ricO*$^{sw}$-*snap* | pMAD containing a *ricO* sandwich fusion to *snap* | This study |
| pMAD-*ricO*$^{SW}$-*gfp* | pMAD containing a *ricO* sandwich fusion to *gfp* | This study |
| pMAD-*scpB-mcherry* | pMAD containing a *scpB-mcherry* fusion and the *scpB* downstream region | This study |
| pMAD-*scpA-mNG* | pMAD containing a *scpA-mNG* fusion and the *scpA* downstream region | This study |

### Table S3. Oligonucleotides used in this study.

| No. | Name | 5'-3' sequence |
|---|---|---|
| 5467 | PstI_RBS_sfGFP_Fwd | AAACTGCAGAAAAAATAAGGAGGAAAAAAAATGAGTAAAGGAGAAGAAC |
| 6282 | sfGFP STOP_EcoRI | GCTGAATTCTTATTTGTATAGTTCATCCATGCC |
| 6981 | DOWN ΔRS02045 fw | CGATGCATGCCATGGTACCCCTTGAATCGAGATGTGGCTATG |
| 6982 | DOWN ΔRS02045 rv | TGCAGAAAGGACTTTAAAGTACTTGTGATTTGAACAC |
| 6983 | Up ΔRS02045 fw | TTTGTGTTCAAATCACAAGTACTTTAAAGTCCTTTCTGC |
| 6984 | Up ΔRS02045 rv | CTTCTAGAATTCGAGCTCCCGTCAGTCTAGCAACTGCAC |
| 7456 | 5aaL-mCherry P1 | TTACTTGTACAGCTCGTCCATGCC |
| 7457 | 5aaL-mCherry P2 | TCCTGCGGCGCCTCCGCTATCATTAAAGAGTTCATGCGC |
| 7472 | scpA-fusion P1 fw | CGATGCATGCCATGGTACCCAGCTATACGTTTTTGTAATCG |
| 7475 | scpA-mNG P2 rv | GATGAGTTATACAAATAACATTATTAGAGGAGTGAACTACC |
| 7476 | scpA-mNG P3 fw | ACGGAGGCGCCGCAGGATCCAAAATGGTAGTTCACTCC |
| 7477 | scpA-fusion P4 rv | CTTCTAGAATTCGAGCTCCCGTTCATGCAATGAAACAGCTTG |
| 7478 | scpB-fusion P1 fw | CGATGCATGCCATGGTACCCTGACCAAAATGTTCGAACATTC |
| 7479 | scpB-mCherry P2 rv | GACGAGCTGTACAAGTAACAATCAAAAAGGAGAAAATAATGACTAAAG |
| 7480 | scpB-mCherry P3 fw | AGCGGAGGCGCCGCAGGAGTCATTATTTTCTCCTTTTTG |
| 7483 | scpB-fusion P4 rv | CTTCTAGAATTCGAGCTCCCATATTGAGCAGCAACGTAATTTTG |
| 7494 | 5aaL-mNG P1 | TTATTTGTATAACTCATCCATGC |
| 7495 | 5aaL-mNG P2 | TCCTGCGGCGCCTCCGTATCAAAAGGTGAAGAAGATAATATG |
| 9436 | SAUSA300_0383_+61_Eco_fwd | ATATGAATTCGTTCAACATTTAGAAGAACGTGG |
| 9626 | tetR_Pae_fwd | ATATGCATGCTTAAGACCCACTTTCACATTTAAG |
| 9627 | Pxyl-tetO_Pst_rev1 | ATATCTGCAGACTCTATCAATGATAGAGAGCTCGATCTACTCTATCAATGATAGAGAGCTTATTTTAATTATACTCTATCA |
| 9630 | 15aa-SNAP_fwd | CAAGGACCAGGATCTGGTCAAGGTTCTGGTGACAAAGATTGCGAAATGAAACG |
| 9632 | SNAP-10aa_rev | AGAGCCACCTCCGCCAGAACCGCCTCCACCTCCCAGACCCGGTTTACCCAG |
| 9906 | OBCBAIM067 | AGATTGGTGGTAGAAGAGCAGAGCTCATGTTAACTAAAGAATTTGCACAACG |
| 9907 | OBCBAIM068 | CTTTCGGGCTTTGTTAGCAGCCGGATCCTTACAGGTTGAATAAACGTGCGAA |
| 9951 | 0009_Bam_fwd | ATATGGATCCGGTTACCAAGCTACAATGAT |
| 9952 | 0009_BglII_rev | ATATAGATCTATACCGGAGTCTTTAGTCAGC |
| 9953 | 0010_BglII_fwd | ATATAGATCTTGATGTCTAATGTTATTGCTTAG |
| 9954 | 0010_Eco_rev | ATATGAATTCGAGTTATCATTGTTCCATCATC |
| 9955 | 2646_Bam_fwd | ATATGGATCCTGTAGATAAACCTTCACGCA |
| 9956 | 2646_BglII_rev | ATATAGATCTGTAAGACTGACCTTAGGGTTATA |
| 9957 | 2647_BglII_fwd | ATATAGATCTTTCGGGATGTGTTGAGTGGTTAATG |
| 9958 | 2647_Eco_rev | ATATGAATTCCAGAGTCTGCATAATCTGGTA |
| 9959 | 2571_Bam_fwd | ATATGGATCCCTAACTTGAGTACACGTGAT |
| 9960 | 2571_BglII_rev | ATATAGATCTTACTCGGCAGTCTAAAATCATTAC |
| 9961 | 2572_BglII_fwd | ATATAGATCTCTCACACTCCTCTCATCTAATTAC |
| 9962 | 2572_Sma_rev | ATATCCCGGGGTGATGGTGATGGTCGCACAT |
| 9963 | 0214_Bam_fwd | ATATGGATCCCAGCCTACTCTGAAATTTATG |
| 9964 | 0214_BglII_rev | ATATAGATCTTTCAGACTAACCTCGAGTTCG |
| 9965 | 0215_BglII_fwd | ATATAGATCTTGGTGGCAGTGTTGAATAAATGC |
| 9966 | 0215_Eco_rev | ATATGAATTCCACCCAAATTACTTCTTAAACATC |
| 10108 | OBCBAIM080 | TCCTTTCGGGCTTTGTTAGCAGCCGGATCCTTATGTTGATTCAGCTAATGCTTTTG |
| 10123 | SAUSA300_0383_-25_Eco_fwd | ATATGAATTCGTATATGCAGAAAGGACTTTAAAGT |
| 10124 | SAUSA300_0383_Bam_rev | ATATGGATCCTTACAGGTTGAATAAACGTGCGAAG |
| 10133 | 0009_oligo_fwd | GATCCAATAAAAAGTTAATATGAAGCTGACTAAAGACTCCGGTAT**GTCTAACCTCAGACAAAC**TGATGTCTAATGTTATTGCTTAGGGTATAGAACTGTATTG |
| 10134 | 0009_oligo_rev | AATTCAATACAGTTCTATACCCTAAGCAATAACATTAGACATCA**GTTTGTCTGAGGTTAGAC**ATACCGGAGTCTTTAGTCAGCTTCATATTAACTTTTTATTG |
| 10162 | OBCBAIM090 | GCGTAGAACTAAGTGAGAAGCAAGTAGCTAAAATTGTTCAACATTTAGAAG |
| 10163 | OBCBAIM091 | TAAATGTTGAACAATTTTAGCTACTTGCTTCTCACTTAGTTCTACGC |
| 10189 | PamaP_Xho_fwd | ATATCTCGAGTTAATAATGAGGTTTTATTTACAAAAACTA |
| 10190 | PamaP_Eco_rev | ATATGAATTCCCTTAGTTAAAGTTAATTTAAAAAC |
| 10191 | TT_Nde_Sma_fwd | ATATCATATGCCCGGGCTAATGCATAATAAATACTG |
| 10192 | TT_Xho_rev | ATATCTCGAGTGTCACTTTGCTTGATATATGAG |
| 10207 | OBCBAIM120 | GACTAAAGACTCCGGTAT**GTCTAACCTCAGACAAAC**TGATGTCTAATGTTATTG |
| 10208 | OBCBAIM121 | CAATAACATTAGACATCA**GTTTGTCTGAGGTTAGAC**ATACCGGAGTCTTTAGTC |
| 10209 | OBCBAIM122 | GACTAAAGACTCCGGTAT*TAAACTGTGGGCTACTCC*TGATGTCTAATGTTATTG |
| 10210 | OBCBAIM123 | CAATAACATTAGACATCA*GGAGTAGCCCACAGTTTA*ATACCGGAGTCTTTAGTC |
| 10254 | SAUSA300_0383_Eco_rev | ATATGAATTCTTACAGGTTGAATAAACGTGCGAAG |
| 10255 | PtcaB_Xho_fwd | ATATCTCGAGTACAATTAGTGTGAATTACTTTGAAG |
| 10256 | PtcaB_Eco_rev | ATATGAATTCTTATATAATGATTAGTTTATAACT |
| 10675 | 0383up_Eco_fwd | ATATGAATTCGTTAATGACGCAAACATTACATC |
| 10676 | 0383down_Sal_rev | ATATGTCGACTGAAGCAGGGCATTAAATGCAA |
| 10678 | 0383C_Sal_rev | ATATGTCGACTTACAGGTTGAATAAACGTGCGAAG |
| 10679 | 0383down_Bam_fwd | ATATGGATCCACTTGTGATTTGAACACAAATTTA |
| 10684 | 0383int-15aa_rev | ACCAGAACCTTGACCAGATCCTGGTCCTTGTCCTGATCCTTCCAAAACAGCATTCTTTTCAACTTTATTTG |
| 10685 | 0383int-10aa_fwd | GGTGGAGGCGGTTCTGGCGGAGGTGGCTCTACTGAAGATAAAGCTGACAGTAA |
| 10687 | 0383+226_Eco_fwd | ATATGAATTCAAAGACTTCCTGCAAGTCATTGTCA |
| 10766 | 0383_+912_+stop_Bam_rev | ATATGGATCCTTAAGGTTTTGGTTCTGACATTTCTGAC |
| 10771 | snap_int_Sal_rev | ATATGTCGACCGGCGGCGGTGGCCGCCGGATTA |

| | | |
|---|---|---|
| 10878 | 0030_Bam_fwd | ATATGGATCCGTTGGAGTTAGATGTTGCAAT |
| 10879 | 0030_BglII_rev | ATATAGATCTGTAAGACTCACCTCTAACTTTC |
| 10880 | 0031_BglII_fwd | ATATAGATCTTTACCTTTAACATGTTACATAC |
| 10881 | 0031_Eco_rev | ATATGAATTCCAAACATGATGATGGCTATTAATG |
| 10882 | 2447_Bam_fwd | ATATGGATCCAATGATTCCACAACTTTTGGATG |
| 10885 | 2448_Eco_rev | ATATGAATTCGTTTACTGAAGCATTGGTGGGAGC |
| 10886 | 15aa-sGFP_fwd | GGACAAGGACCAGGATCTGGTCAAGGTTCTGGTAGTAAAGGAGAAGAACTTTTCACTG |
| 10887 | sGFP-10aa_rev | AGAGCCACCTCCGCCAGAACCGCCTCCACCTTTGTATAGTTCATCCATGCCATGT |
| 10888 | 15aa-mCh_fwd | GGACAAGGACCAGGATCTGGTCAAGGTTCTGGTGCTATCATTAAAGAGTTCATGCG |
| 10889 | mCh-10aa_rev | AGAGCCACCTCCGCCAGAACCGCCTCCACCCTTGTACAGCTCGTCCATGCCA |
| 11094 | SAUSA300_0383_st7_Pst-Sal_fwd | ATATCTGCAGGTCGACAAAAAATAAGGAGGAAAAAAAATGTTAACTAAAGAATTTGCACAAC |
| 11243 | 2447_BglII_rev2 | CAGGTCAGACTAACAGATCTTGACGGGAGTCTAAAAAAATGTG |
| 11244 | 2448_BglII_fwd2 | GTCAAGATCTGTTAGTCTGACCTGATAAATATTT |
| 11245 | 0106_Bam_fwd | ATATGGATCCTGTTTATCACGCACCTGGTGTTG |
| 11246 | 0106_BglII_rev | CAGTGTCCGTCAAGATCTGAAGCATTATTATATCAGTG |
| 11247 | 0107_BglII_fwd | CTTCAGATCTTGACGGACACTGTTCTCGG |
| 11248 | 0107_Eco_rev | ATATGAATTCGGTGTTATTGATGATTCGAGTTGGT |
| 11249 | 0384_Bam_fwd | ATATGGATCCGAGATTACTGTACTGTCTAATTTAAG |
| 11250 | 0384_BglII_rev | GTATACTGACGTTTAGATCTGTCAGTCTTGCTTCATACA |
| 11251 | 0383_BglII_fwd | TGACAGATCTAAACGTCAGTATACATAAAG |
| 11253 | 2627_Bam_fwd | ATATGGATCCGAATCAATTGTTAGGCAAGCCCATTCA |
| 11254 | 2627_BglII_rev | ATACGTCTGTCAAGATCTCTCAGTCTCACCTCATCATG |
| 11255 | 2628_BglII_fwd | TGAGAGATCTTGACAGACGTATAAATTTG |
| 11256 | 2628_Sal_rev | ATATGTCGACCAGGTGCTTGGAACACATTAGTATG |

Underlined sequences indicate restriction sites used for cloning. The *ricS1* sequence is highlighted in bold and the modified *ricS1* sequence (*ricS1**) is italicised.

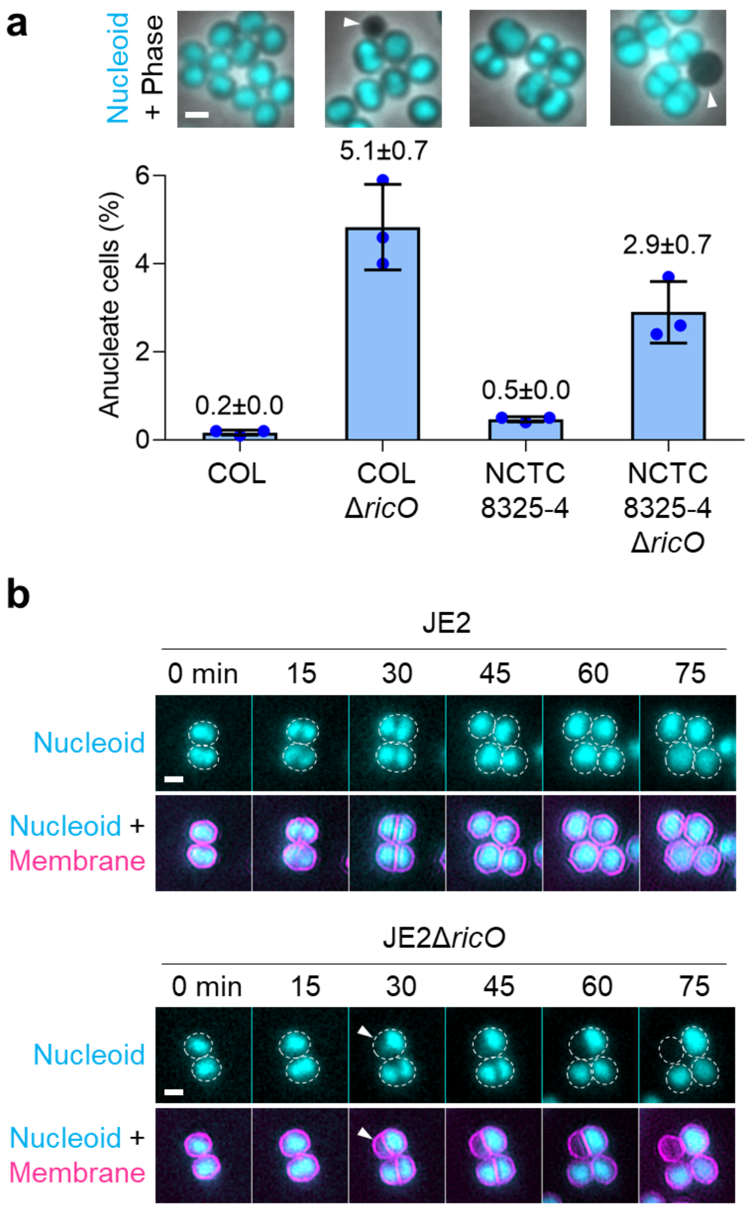

**Figure S1. Deletion of *ricO* causes chromosome segregation defects in different *S. aureus* strains.**

a) *ricO* deletion in the *S. aureus* strains COL and NCTC8325-4 leads to the formation of anucleate cells. The bar graph shows the percentage of anucleate cells measured for the indicated strains, each labelled with DNA dye Hoechst 33342. Bars represent the mean and lines indicate the standard deviation of three independent experiments. At least 1239 cells were analysed for each biological replicate. Fluorescence micrographs overlaid with phase contrast images show cells with labelled DNA (cyan), and anucleate cells are indicated by white arrowheads. Scale bar, 1 µm.

b) *ricO* deletion in the *S. aureus* strain JE2 leads to impaired chromosome segregation in a fraction of cells. Cells of the JE2 wild-type and JE2Δ*ricO* strains, each stained with DNA dye Hoechst 33342 (cyan) and membrane dye CellBrite Fix 640 (magenta), were imaged by time-lapse microscopy. Overlaid fluorescence micrographs show normal nucleoid segregation during the cell cycle and impaired nucleoid segregation in JE2Δ*ricO* cells (indicated by white arrowheads). Cell outlines inferred from stained membranes are indicated by dashed white ovals. Scale bar, 1 µm.

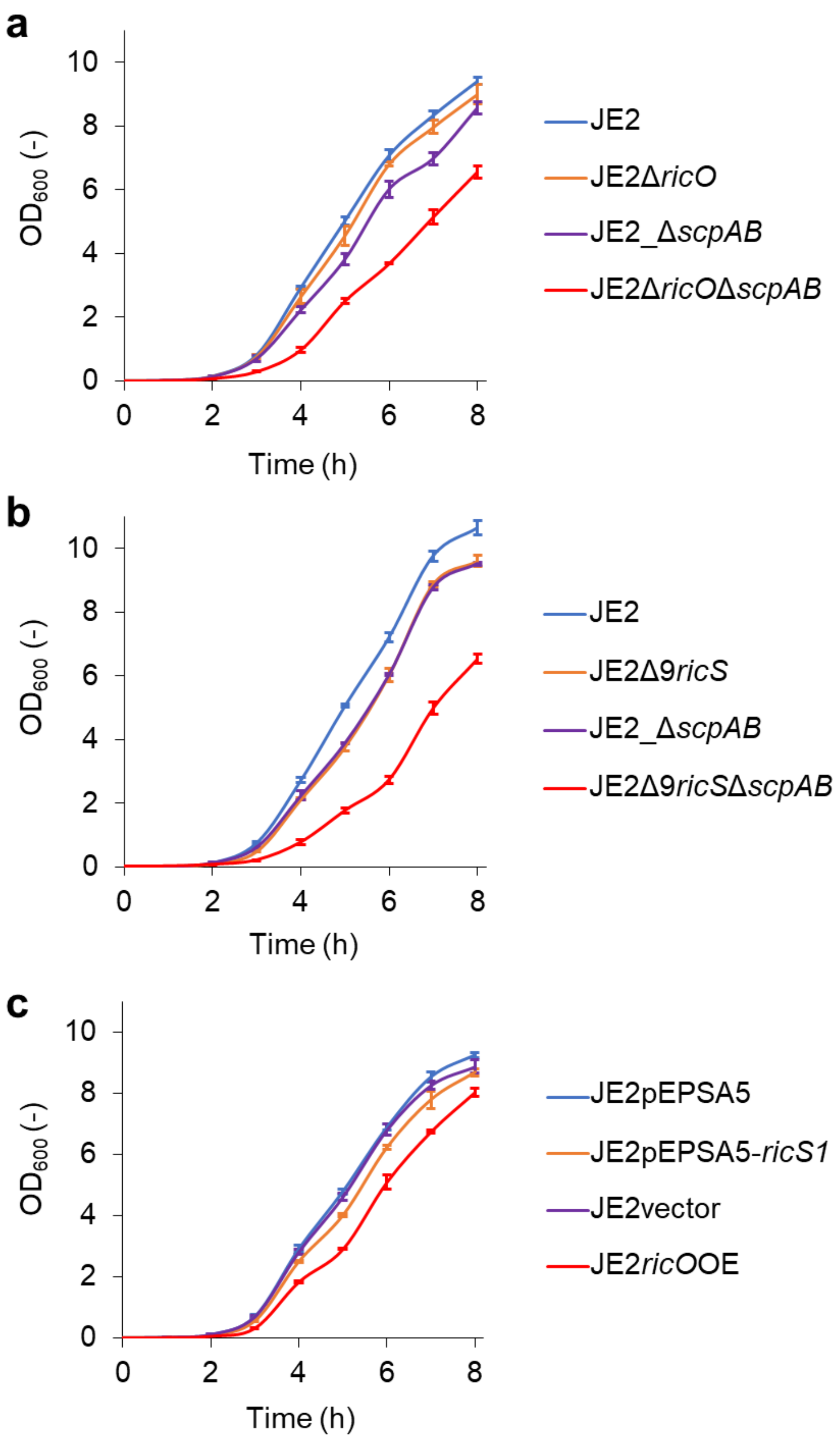

**Figure S2. Growth curves of *S. aureus* strains used in this study.**

a-c) Mean optical density over time measured for the indicated strains, each grown in TSB rich medium at 37°C. Error bars represent the standard deviation of three biological replicates.

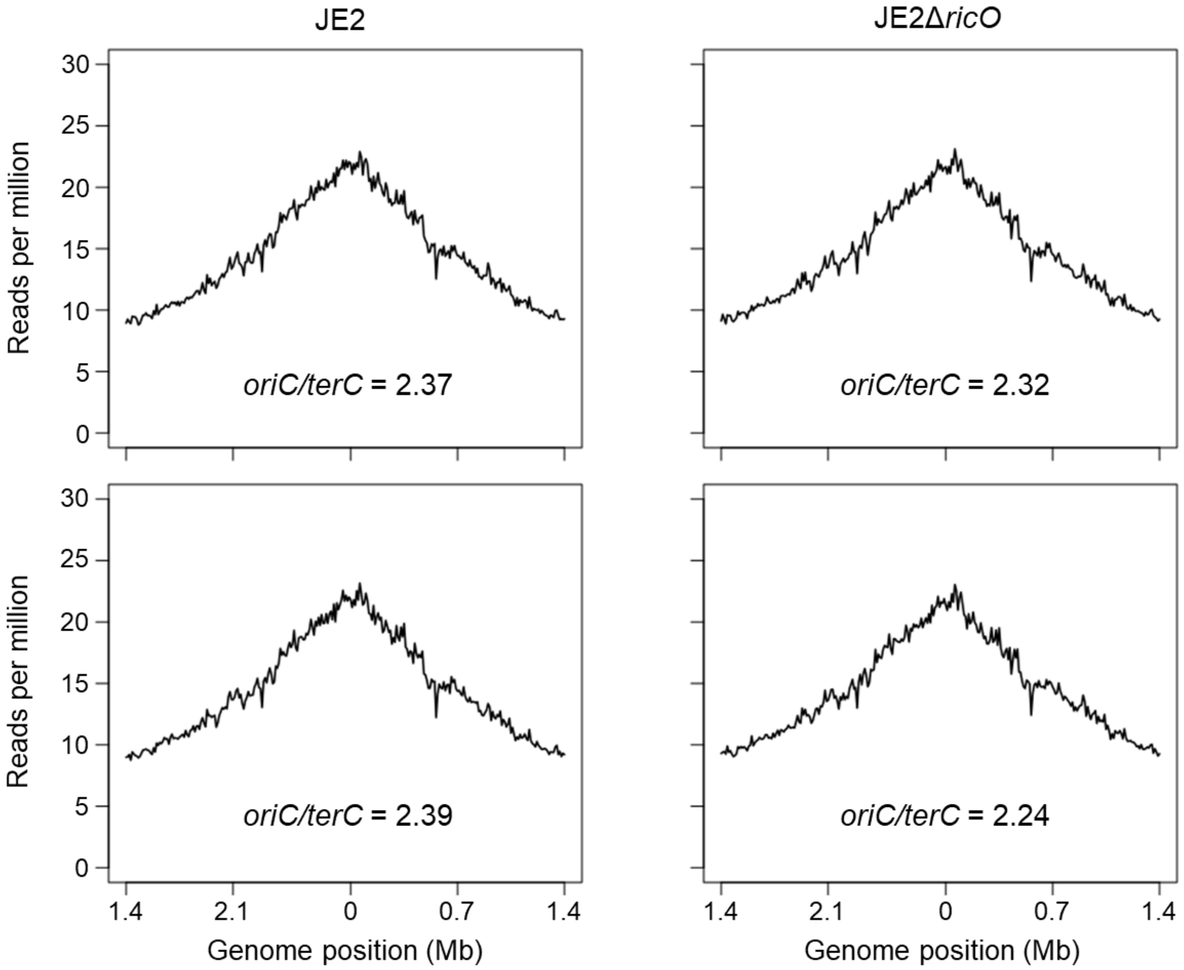

**Figure S3.** ***S. aureus*** **cells lacking RicO are not affected in DNA replication.**

The indicated strains were grown to early-exponential phase in TSB rich medium at 37°C and analysed by whole-genome sequencing. Marker frequency analysis confirmed a similar relative number of reads determined for the replication origin and terminus regions (*oriC/terC* ratio) in JE2 wild-type and JE2Δ*ricO* strains. Data from two biological replicates are shown.

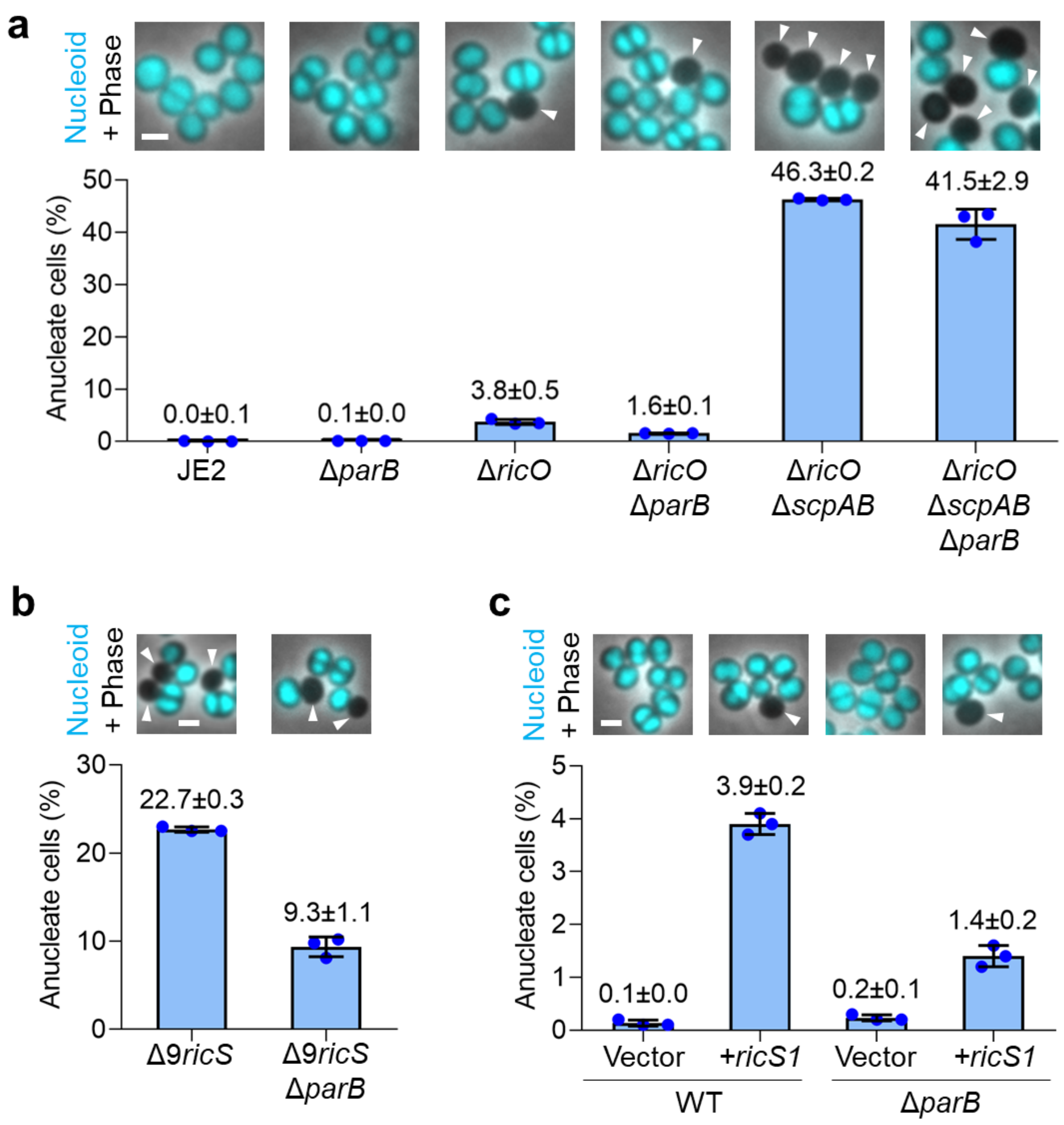

**Figure S4. Deletion of *parB* alleviates defects caused by deletion of *ricO*, deletion of nine *ricS* sites or the presence of extra *ricS* copies.**

a-c) ParB is partially responsible for the chromosome segregation defects resulting from impaired RicO function. Bar graphs show the percentage of anucleate cells measured for each strain. Strains in panel a: JE2, JE2_Δ*parB*, JE2Δ*ricO*, JE2Δ*ricO*Δ*parB*, JE2Δ*ricO*Δ*scpAB*, JE2Δ*ricO*Δ*scpAB*Δ*parB.* Strains in panel b: JE2Δ9*ricS*, JE2Δ9*ricS*Δ*parB.* Strains in panel c: JE2pEPSA5, JE2pEPSA5-*ricS1*, JE2Δ*parB*pEPSA5 and JE2Δ*parB*pEPSA5-*ricS1*. Bars represent the mean and lines indicate the standard deviation of three independent experiments. At least 982 cells were analysed for each biological replicate. Fluorescence micrographs overlaid with phase contrast images show cells labelled with DNA dye Hoechst 33342 (cyan), and anucleate cells are indicated by white arrowheads. Scale bars, 1 µm.

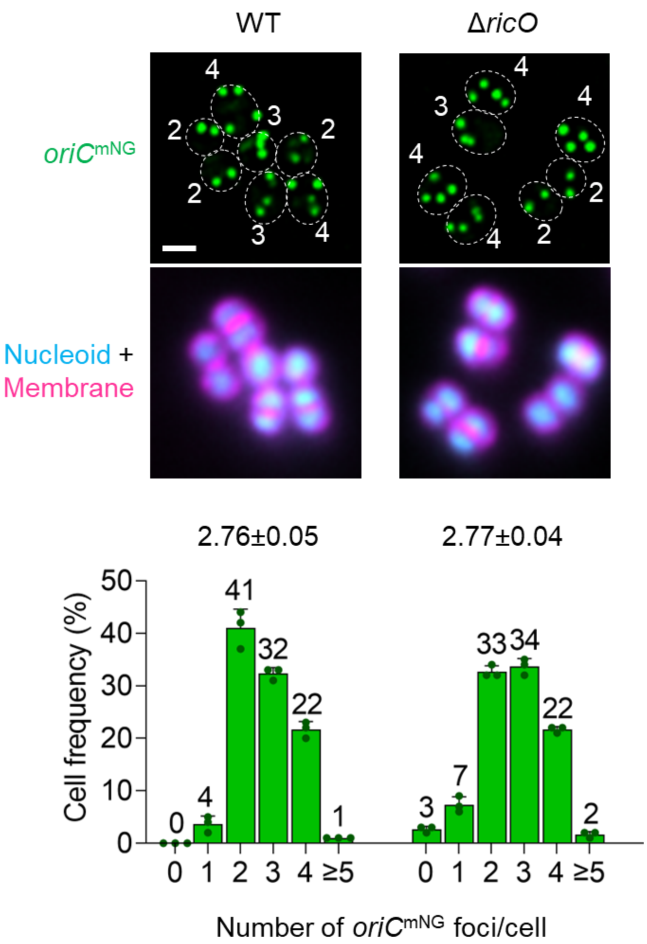

**Figure S5. Deletion of *ricO* in *S. aureus* does not impede initial segregation of newly replicated origins.**

The number of replication origins per cell is similar in the presence and absence of RicO. The bar graph shows the number of fluorescent foci corresponding to replication origins determined for cells of the strains JE2_FROS$^{Ori}$ (WT, left) and JE2*oriC*$^{mNG}$Δ*ricO* (right). Bars represent the mean and lines indicate the standard deviation of three independent experiments. At least 470 cells were analysed for each biological replicate. The mean and standard deviation of the origin copy number shown on top was calculated from cells displaying at least one fluorescent focus. Replication origins (green) were fluorescently labelled by producing TetR-mNG and introducing a *tetO* array at position 358° of the circular genome. The number of diffraction-limited foci is indicated for each cell in the fluorescence micrographs. Cells were labelled with the membrane dye CellBrite Fix 640 (magenta) and DNA dye Hoechst 33342 (cyan). Cell outlines inferred from stained membranes are indicated by dashed white ovals. Scale bar, 1 µm.

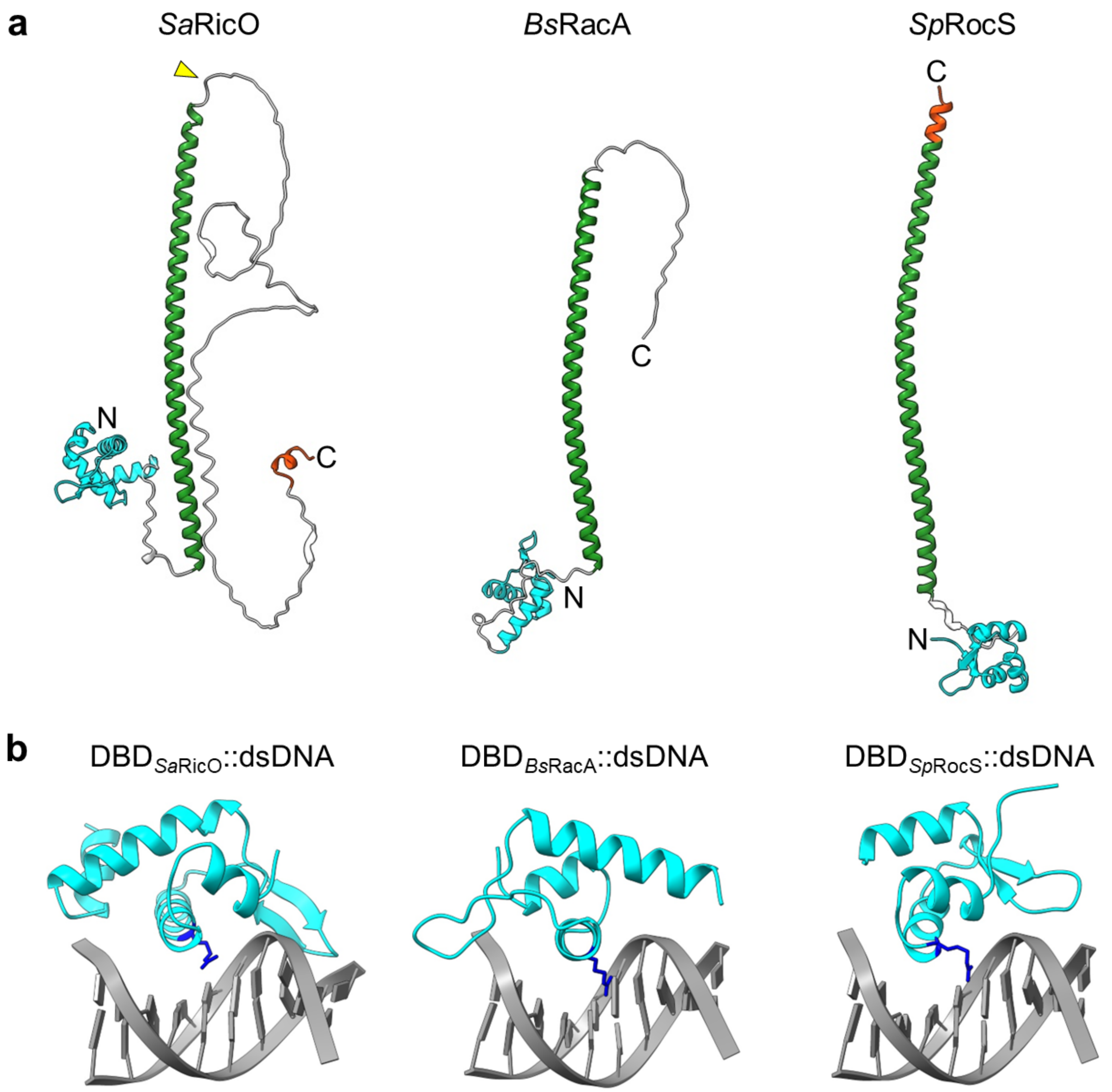

**Figure S6. *Staphylococcus aureus* RicO, *Bacillus subtilis* RacA and *Streptococcus pneumoniae* RocS share similar structural features.**

a) AlphaFold [12] structure predictions of *Sa*RicO, *Bs*RacA and *Sp*RocS with DNA-binding domains highlighted in cyan, coiled-coil domains highlighted in green, and amphipathic helices highlighted in orange. The insertion site for fluorescent protein tagging of RicO is indicated by a yellow arrowhead.

b) AlphaFold 3 [13] structure predictions of the *Sa*RicO, *Bs*RacA and *Sp*RocS DNA-binding domains (cyan) in complex with double-stranded DNA (gray). The positively charged residues *Sa*RicO(R18), *Bs*RacA(R19) and *Sp*RocS(R18) located in the DNA interface are highlighted in dark blue.

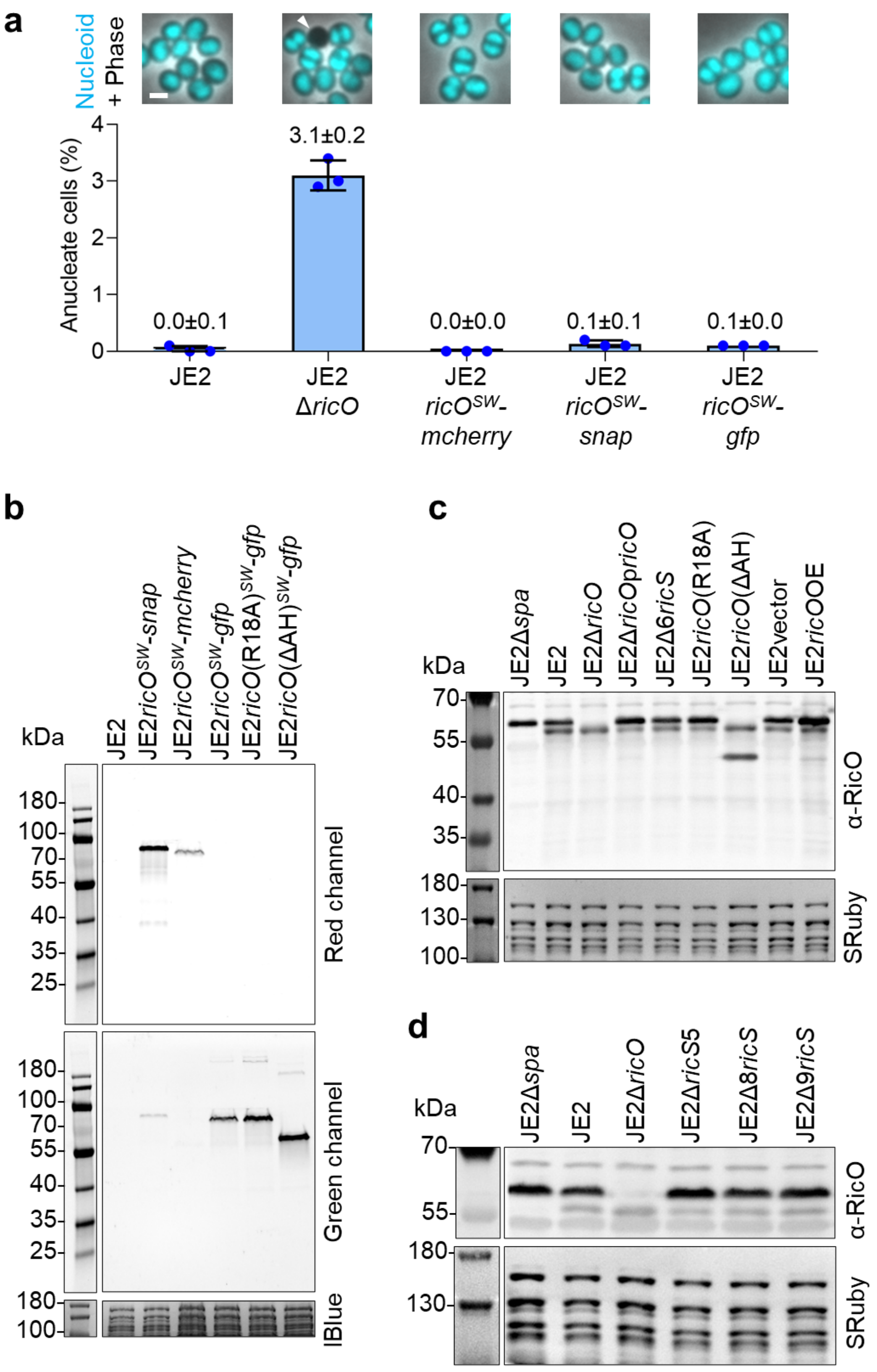

**Figure S7. Functionality and stability of RicO variants used in this study.**

a) Insertion of protein tags between valine 182 and threonine 183 in the RicO protein does not affect its function. The bar graph shows the percentage of anucleate cells measured for the indicated strains, each labelled with DNA dye Hoechst 33342. Bars represent the mean and lines indicate the standard deviation of three independent experiments. At least 779 cells were analysed for each biological replicate. Fluorescence micrographs overlaid with phase contrast images show cells with labelled DNA (cyan), and anucleate cells are indicated by white arrowheads. Scale bar, 1 µm.

b) RicO protein fusions are subject to minor or no apparent proteolytic cleavage. Fluorescence gel analysis of non-boiled total protein extracts obtained from the indicated strains. JE2 and JE2*ricO*$^{SW}$-*snap* were labelled with 83 nM of the SNAP-tag ligand JF549-cpSTL. SNAP-, mCherry- and GFP-tags were inserted between valine 182 and threonine 183 in the RicO protein. Images show in-gel protein fluorescence detected in red (top) and green (middle) channels. Gel was post-stained with Instant Blue (IBlue) to visualize total proteins (bottom). Theoretical molecular weights (kDa): RicO$^{SW}$-SNAP, 56.6. RicO$^{SW}$-mCherry, 63.1. RicO$^{SW}$-GFP, 64.2. RicO(R18A)$^{SW}$-GFP, 64.1. RicO(ΔAH)$^{SW}$-GFP, 62.5.

c,d) Western Blot analysis of total protein extracts obtained from the indicated strains using anti-RicO antibody (top). Membrane was stained with SyproRuby (SRuby) to visualize total proteins that migrated above the 100-kDa size marker band (bottom). Theoretical molecular weights (kDa): RicO, 35.6. RicO(R18A), 35.5. RicO(ΔAH), 33.9. Spa, 55.6.

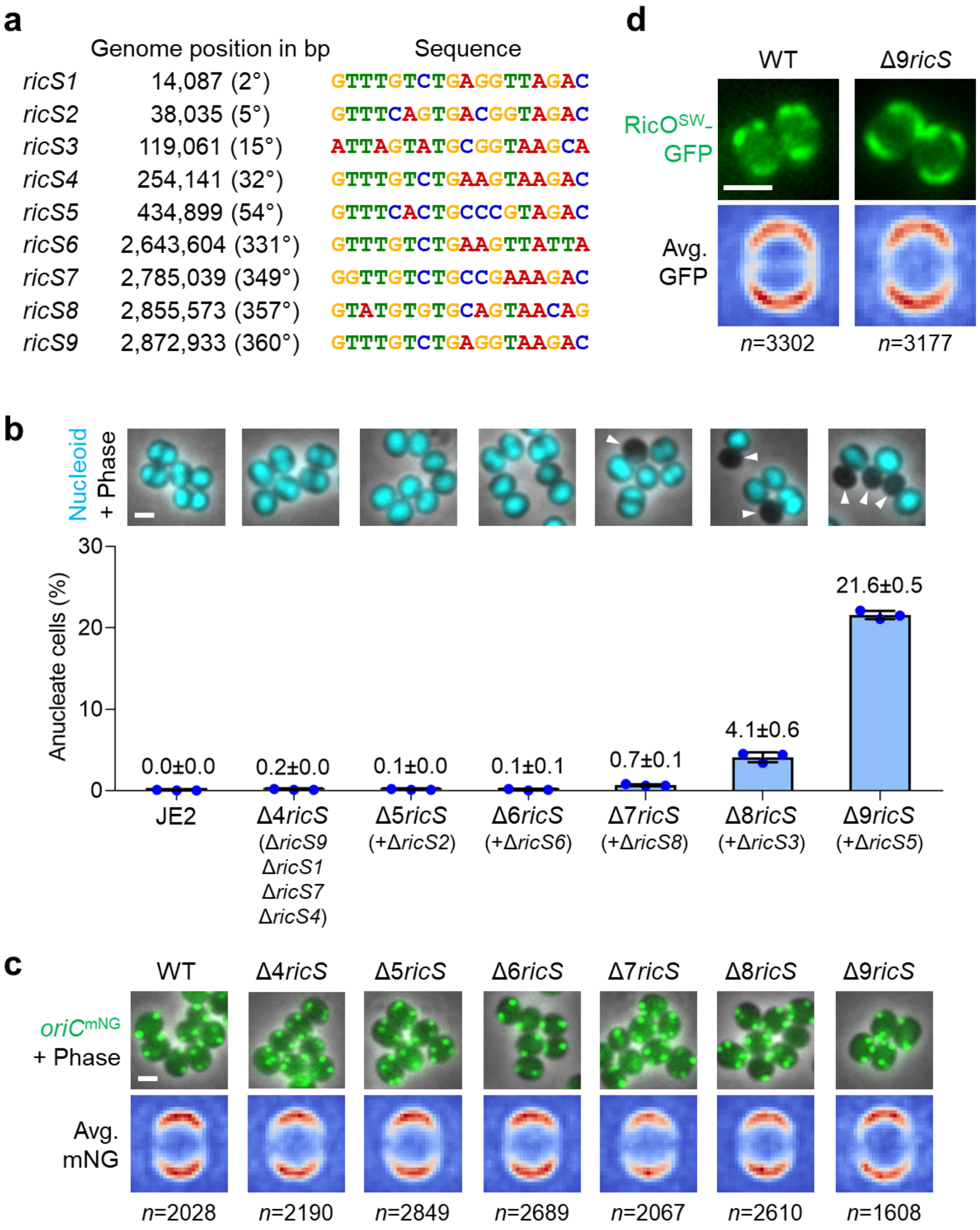

**Figure S8. Multiple *ricS* deletions cause defects in chromosome segregation but no apparent changes in localization of replication origins or RicO.**

a) Conserved sequences identified at the nine loci of the *S. aureus* JE2 genome targeted by RicO.

b) *ricS* sites are required for normal chromosome segregation. Bar graph shows the percentage of anucleate cells measured for the strains (from left to right) JE2, JE2Δ4*ricS,* JE2Δ5*ricS,* JE2Δ6*ricS,* JE2Δ7*ricS,* JE2Δ8*ricS* and JE2Δ9*ricS*. Bars represent the mean and lines indicate the standard deviation of three independent experiments. At least 1594 cells were analysed for each biological replicate. Fluorescence micrographs overlaid with phase contrast images show cells labelled with DNA dye Hoechst 33342 (cyan), and anucleate cells are indicated by white arrowheads. Scale bar, 1 µm.

c) *ricS* sites are dispensable for polar localization of chromosomal origins. Fluorescence micrographs overlaid with phase contrast images of the strains (from left to right): JE2_FROS$^{Ori}$ (WT), JE2*oriC*$^{mNG}$Δ4*ricS,* JE2*oriC*$^{mNG}$Δ5*ricS,* JE2*oriC*$^{mNG}$Δ6*ricS,* JE2*oriC*$^{mNG}$Δ7*ricS,* JE2*oriC*$^{mNG}$Δ8*ricS,* JE2*oriC*$^{mNG}$Δ9*ricS.* Replication origins (green) were fluorescently labelled by producing TetR-mNG and introducing a *tetO* array at position 358° of the circular genome. Lower row shows heat maps of the average localization of detected fluorescent foci corresponding to replication origins of the same strains as cells shown in the upper row. Blue-white-red color code in heat maps represents spot density from low to high. *n*, number of analysed cells. Scale bar, 1 µm.

d) *ricS* sites are dispensable for polar localization of RicO. Fluorescence micrographs overlaid with phase contrast images of the strains JE2*ricO*$^{SW}$*-gfp* (WT) and JE2Δ9*ricSricO*$^{SW}$*-gfp* (Δ9*ricS*). Lower row shows heat maps of the average localization of detected fluorescent foci corresponding to RicO of the same strains as cells shown in the upper row. Blue-white-red color code in heat maps represents spot density from low to high, respectively. Data shown are from three biological replicates. *n*, number of analysed cells. Scale bar, 1 µm.

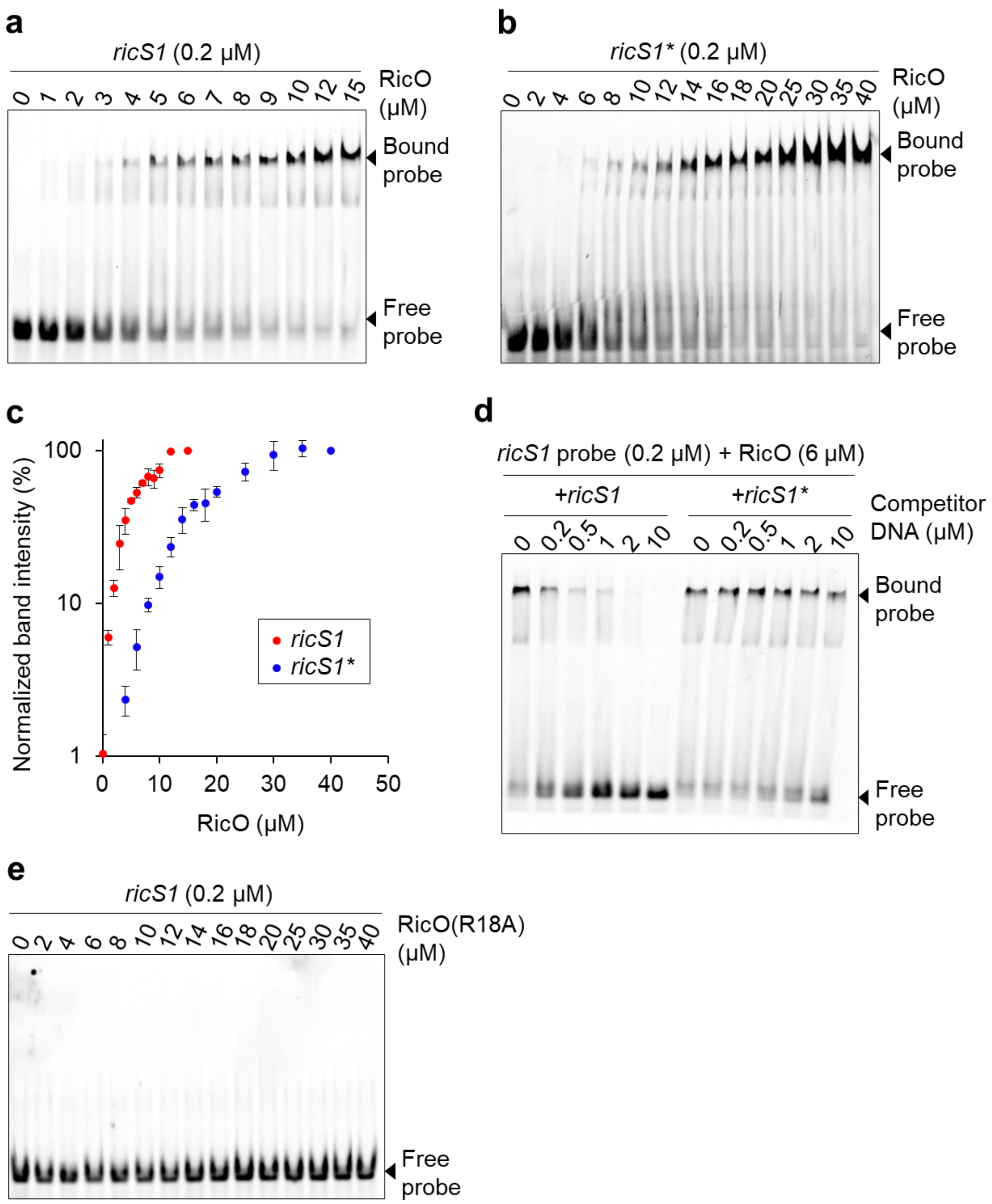

**Figure S9. RicO preferentially binds DNA containing the *ricS* motif *in vitro*.**

a-c) Electrophoretic mobility shift assay using purified RicO$^{1\text{-}174}$ at different concentrations and a 54-bp DNA fragment containing either *ricS1* (panel a) or its mutated derivative *ricS1** (panel b) (Supplementary Table 3). The graph in panel c shows band intensities obtained from image data represented in panels a and b. Error bars indicate the range values obtained from two to three biological replicates.

d) Electrophoretic mobility shift assay using purified RicO$^{1\text{-}174}$ and a fluorescently labelled 54-bp DNA fragment containing the *ricS1* sequence. The RicO-*ricS* interaction was challenged by titrating an unlabelled 54-bp DNA fragment either with the identical sequence or its mutated derivative *ricS1** into the reaction.

e) Electrophoretic mobility shift assay using different concentrations of purified RicO$^{1\text{-}174}$ containing amino acid substitution R18A and a 54-bp DNA fragment containing *ricS1*.

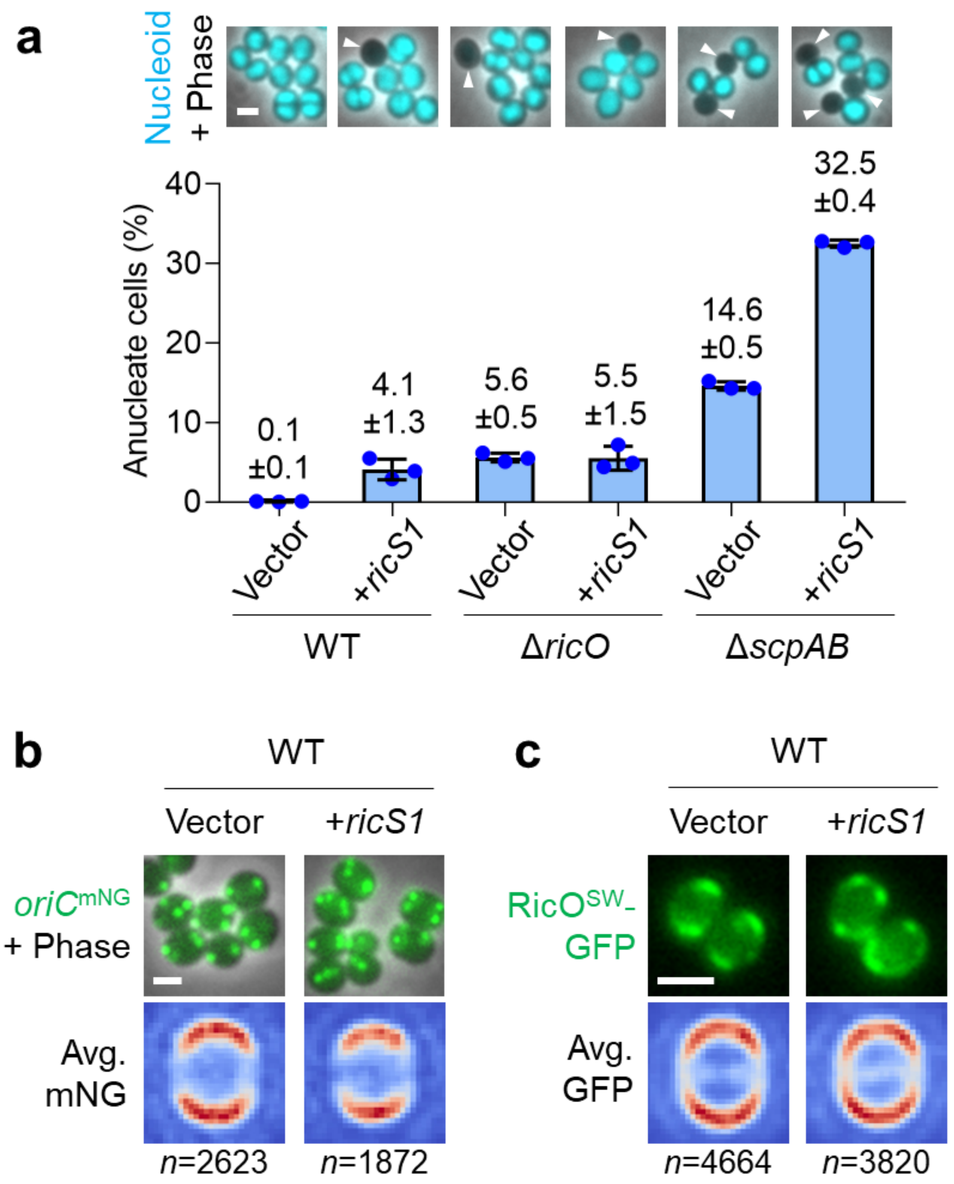

**Figure S10. Extra *ricS* copies cause defects in chromosome segregation but no apparent changes in localization of replication origins or RicO.**

a) Ectopic *ricS* sites impair chromosome segregation. Bar graph shows the percentage of anucleate cells measured for the strains (from left to right) JE2pEPSA5, JE2pEPSA5-*ricS1,* JE2Δ*ricO*pEPSA5, JE2Δ*ricO*pEPSA5-*ricS1,* JE2Δ*scpAB*pEPSA5 and JE2Δ*scpAB*pEPSA5-*ricS1*. Strains harbouring pEPSA5-*ricS1* have the *ricS1* sequence in a replicative plasmid. Bars represent the mean and lines indicate the standard deviation of three independent experiments. At least 710 cells were analysed for each biological replicate. Fluorescence micrographs overlaid with phase contrast images show cells labelled with DNA dye Hoechst 33342 (cyan), and anucleate cells are indicated by white arrowheads. Scale bar, 1 μm.

b) Ectopic *ricS* sites do not affect polar localization of chromosomal origins. Fluorescence micrographs overlaid with phase contrast images of the strains JE2*oriC*$^{mNG}$pEPSA5 (left) and JE2*oriC*$^{mNG}$pEPSA5-*ricS1* (right). Replication origins (green) were fluorescently labelled by producing TetR-mNG and introducing a *tetO* array at position 358° of the circular genome. Lower row shows heat maps of the average localization of detected fluorescent foci corresponding to replication origins of the same strains as cells shown in the upper row. Blue-white-red color code in heat maps represents spot density from low to high, respectively. *n*, number of analysed cells. Scale bar, 1 μm.

c) Ectopic *ricS* sites do not affect polar localization of RicO. Fluorescence micrographs overlaid with phase contrast images of the strains JE2*ricO*$^{SW}$-*gfp*pEPSA5 (left) and JE2*ricO*$^{SW}$-*gfp*pEPSA5-*ricS1* (right). Lower row shows heat maps of the average localization of detected fluorescent foci corresponding to RicO of the same strains as cells shown in the upper row. Blue-white-red color code in heat maps represents spot density from low to high, respectively. Data shown are from three biological replicates. *n*, number of analysed cells. Scale bar, 1 µm.

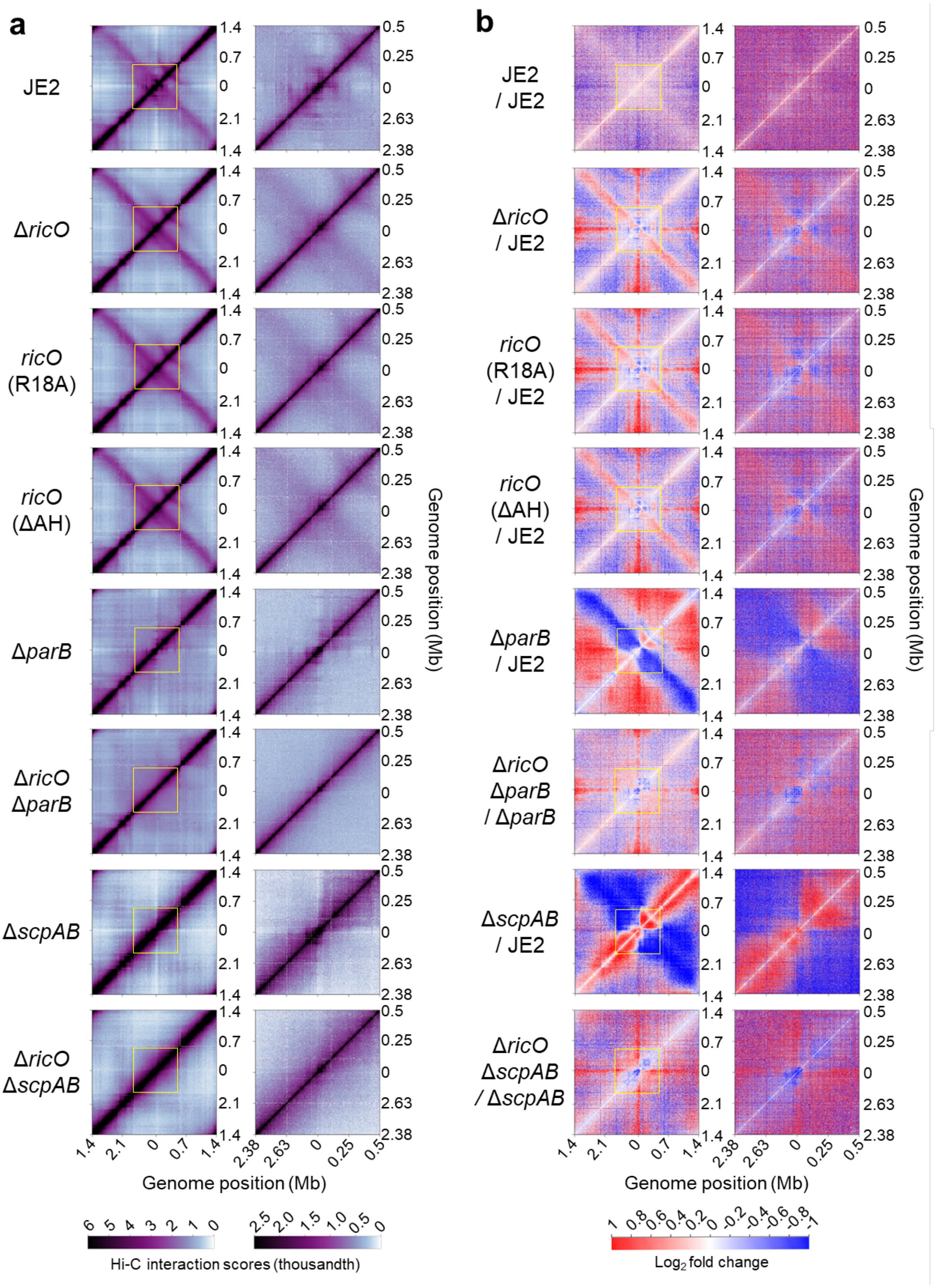

**Figure S11. RicO-dependent genomic interactions in the chromosomal origin region are maintained upon deletion of *parB* and *scpAB*.**

a) Normalized Hi-C contact maps of the strains (from top to bottom) JE2, JE2Δ*ricO*, JE2*ricO*(R18A), JE2*ricO*(ΔAH), JE2_Δ*parB,* JE2Δ*ricO*Δ*parB,* JE2_Δ*scpAB* and JE2Δ*ricO*Δ*scpAB*. Hi-C contact maps display contact frequencies for pairs of 5-kb bins. Contact maps on the right show a zoom into the 1-

Mb region flanking the replication origin (highlighted by yellow squares in the corresponding whole-genome maps) and display frequencies for pairs of 1-kb bins. Data obtained from two biological replicates were combined into one contact map for each strain, while a single biological replicate was analysed for the JE2*ricO*(ΔAH) and JE2Δ*ricO*Δ*scpAB* strains. Data for strains JE2, JE2_Δ*parB* and JE2_Δ*scpAB* were previously published [5], and derive from samples prepared simultaneously with the remaining samples included in this figure.

b) Ratio plots of the Hi-C contact maps shown in panel a. Red and blue indicate increased and reduced chromosomal interactions, respectively, detected in JE2 (the first biological replicate) vs. JE2 (the second biological replicate), JE2Δ*ricO* vs. JE2 (the first and second biological replicates combined), JE2*ricO*(R18A) vs. JE2, JE2*ricO*(ΔAH) vs. JE2, JE2_Δ*parB* vs. JE2, JE2Δ*ricO*Δ*parB* vs. JE2_Δ*parB*, JE2_Δ*scpAB* vs. JE2, and JE2Δ*ricO*Δ*scpAB* vs. JE2_Δ*scpAB*.

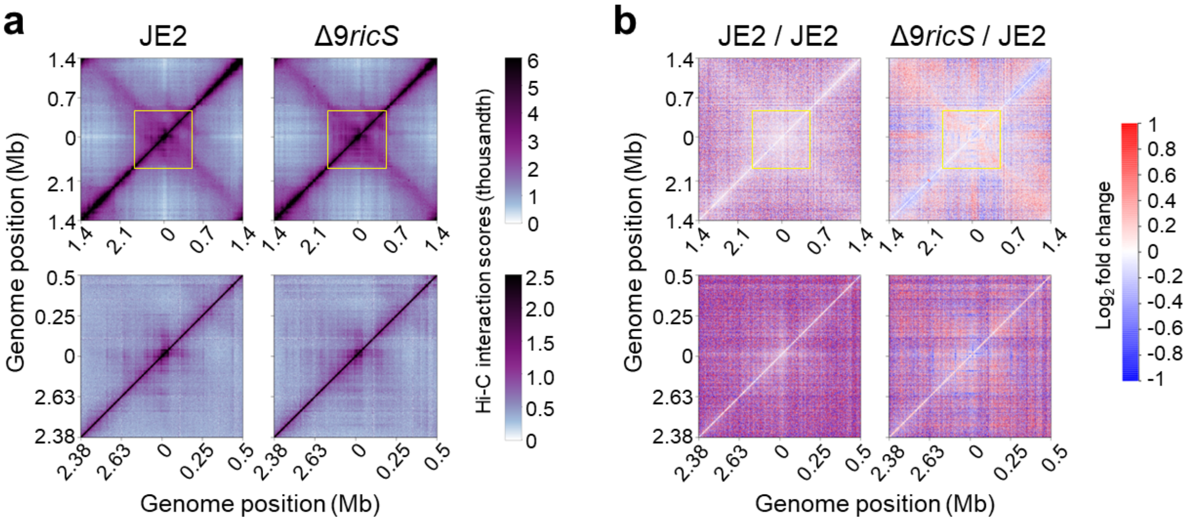

**Figure S12. RicO-dependent genomic interactions in the chromosomal origin region are only marginally reduced upon deletion of nine *ricS* sites.**

a) Normalized Hi-C contact maps of the strains JE2 (left) and JE2Δ9*ricS* (right) displaying contact frequencies for pairs of 5-kb bins. Contact maps in the lower row show a zoom into the 1-Mb region flanking the replication origin (highlighted by yellow squares in the corresponding whole-genome maps) and display frequencies for pairs of 1-kb bins. Data obtained from two biological replicates were combined into one contact map for each strain.

b) Ratio plots of the Hi-C contact maps shown in panel a. Red and blue indicate increased and reduced chromosomal interactions, respectively, detected in JE2 (the third biological replicate) vs. JE2 (the fourth biological replicate) and JE2Δ9*ricS* vs. JE2 (the third and fourth biological replicates combined).

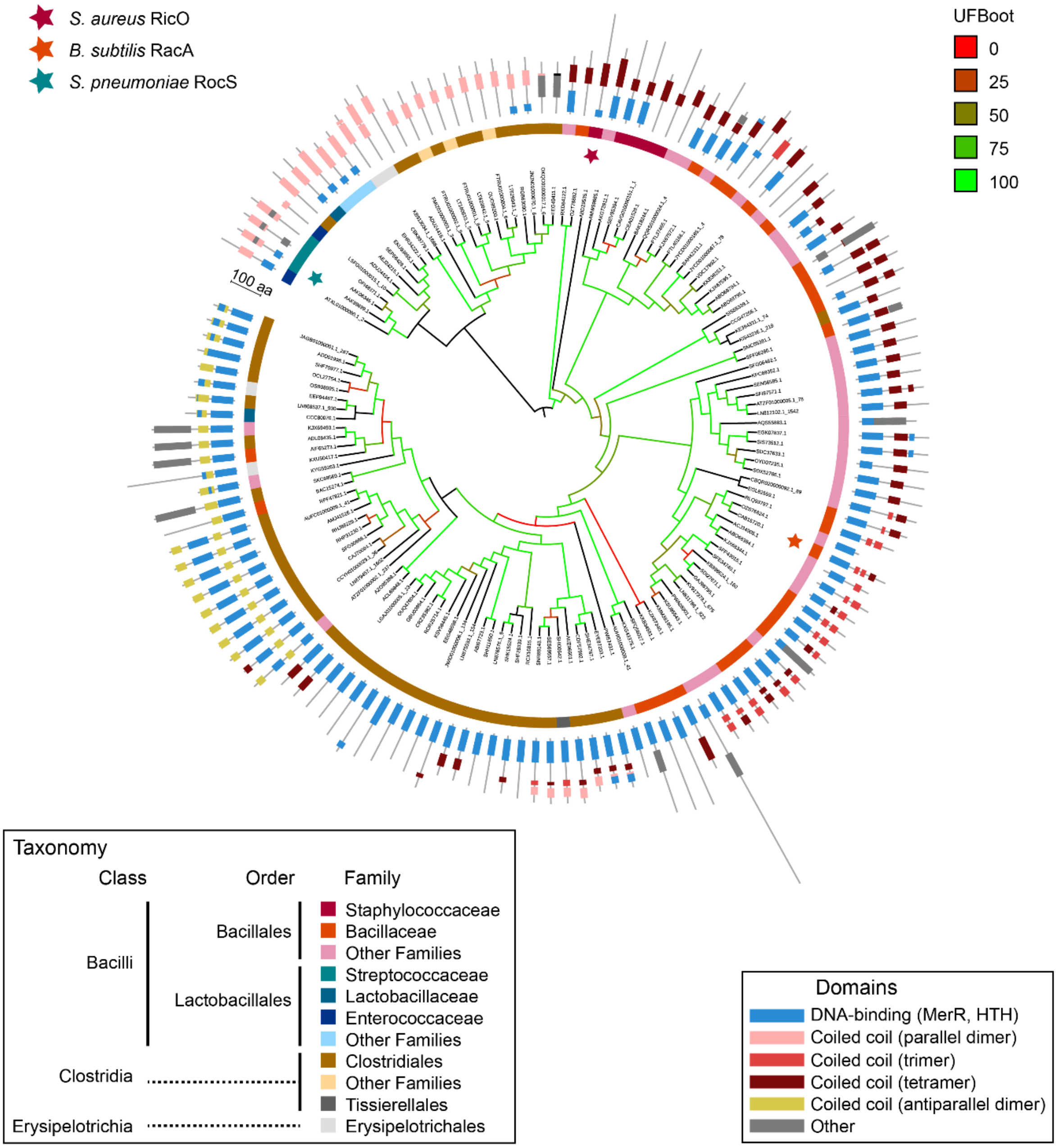

**Figure S13. Phylogenetic tree of potential RicO homologs.**

A maximum likelihood phylogenetic tree based on an alignment (185 amino acid positions) of the 137 homologs of RicO. The tree was inferred with IQ-TREE v3.0.1 using the model Q.PFAM+F+G4. Ultra fast bootstrapping values are colored according to the scale in the upper right corner. In front of each leaf is represented, from inside to outside, (1) a colored bar indicating the taxonomy of the sequence (legend in the lower left corner), (2) the Pfam domains organisation and coiled-coil domain prediction by CoCoNat (legend in the lower right corner). In the absence of an outgroup, the tree is tentatively rooted between two clades, one of them containing only proteins predicted to form parallel dimers. The star symbols indicate the locations of *S. aureus* RicO (dark red), *B. subtilis* RacA (orange) and *S. pneumoniae* RocS (teal). The scale indicates a length of 100 amino acids in the tree gap in the upper left area.

Supplementary Figure 7b

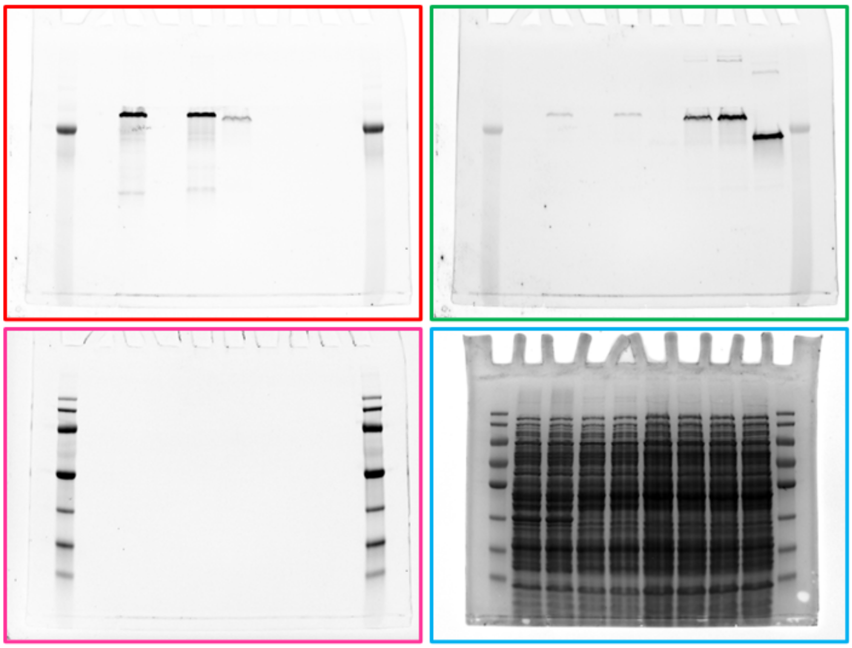

Supplementary Figure 7c

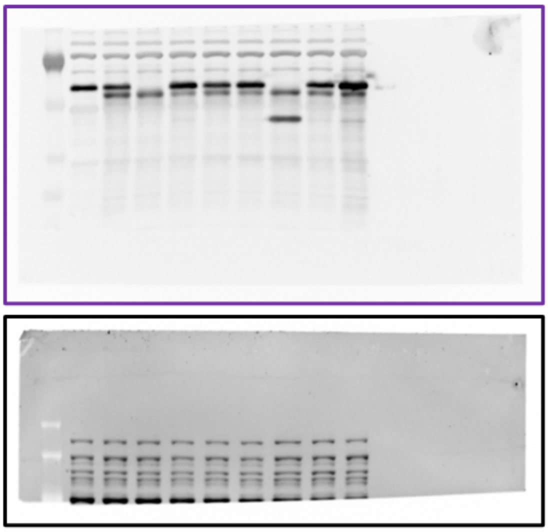

Supplementary Figure 7d

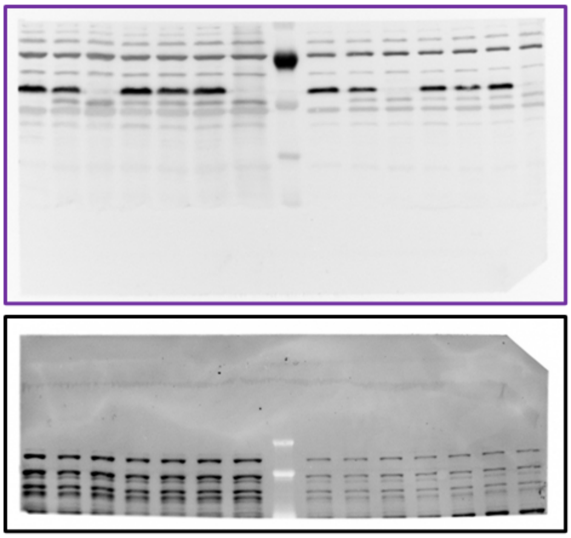

**Figure S14. Uncropped images of gels and blotted membranes.**

Upper panel, RicO$^{SW}$-SNAP labelled with JF549, RicO$^{SW}$-mCherry, RicO$^{SW}$-GFP, RicO(R18A)$^{SW}$-GFP, RicO(ΔAH)$^{SW}$-GFP and molecular weight markers visualized by red (outlined in red), green (outlined in green) and far-red (outlined in magenta) fluorescence detections. Total proteins were visualized by post-staining with Instant Blue Coomassie (outlined in blue). Lower panels, RicO labelled with Star-Bright (outlined in purple) and total proteins stained with Sypro-Ruby (outlined in black).
